# Intratumoral heterogeneity and clonal evolution induced by HPV integration

**DOI:** 10.1101/2022.08.11.503647

**Authors:** Keiko Akagi, David E. Symer, Medhat Mahmoud, Bo Jiang, Sarah Goodwin, Darawalee Wangsa, Zhengke Li, Weihong Xiao, Joe Dan Dunn, Thomas Ried, Kevin R. Coombes, Fritz J. Sedlazeck, Maura L. Gillison

## Abstract

The human papillomavirus (HPV) genome is integrated into host DNA in most HPV-positive cancers, but the consequences for chromosomal integrity are unknown. Continuous long-read sequencing of oropharyngeal cancers and cancer cell lines revealed a unique form of structural variation, i.e., heterocateny, characterized by diverse, interrelated, and repetitive patterns of concatemerized virus and host DNA segments within a cancer. Unique breakpoint sequences shared across structural variants facilitated stepwise reconstruction of their evolution from a common molecular ancestor. This analysis revealed that virus and virus-host concatemers are unstable and, upon insertion into and excision from chromosomes, facilitate capture, amplification, and recombination of host DNA and chromosomal rearrangements. Evidence of heterocateny was detected in extrachromosomal and intrachromosomal DNA. The data indicate that heterocateny is driven by the dynamic, aberrant replication and recombination of an oncogenic DNA virus, thereby extending known consequences of HPV integration to include promotion of intratumoral heterogeneity and clonal evolution.

## INTRODUCTION

Human papillomavirus (HPV) causes more than 630,000 cancers worldwide each year, including anogenital and oropharyngeal squamous cell carcinomas (1). Early in infection, the viral genome is maintained as an ∼8-kilobase pair (kb) extrachromosomal circular DNA (ecDNA), i.e., an episome. In a majority of subsequent cancers, HPV DNA has integrated into the host genome, connecting viral and cellular DNAs at breakpoints (2-5) in intrachromosomal DNA (icDNA) and/or ecDNA forms (6). HPV integration promotes tumorigenesis by increasing expression and stability of transcripts encoding the E6 and E7 oncoproteins (7), which target tumor suppressors p53 and pRb for degradation, respectively (8, 9). Recent whole genome sequencing (WGS) analyses of cervical and oropharyngeal cancers revealed that HPV integrants are enriched in genomic regions with structural variants (SVs) and copy number variants (CNVs) (2, 4, 5, 10). Diverse genetic consequences of HPV integration have been identified, including dysregulated host gene expression near integrants (2-5).

An improved understanding of the mechanisms by which HPV integration leads to SVs, CNVs, and aberrant host gene expression depends upon enhanced resolution of genomic sequence variants and their connectivity. To resolve the structures of genomic rearrangements flanking HPV integration sites, we conducted continuous long-read sequencing (LR-seq) of HPV-positive oropharyngeal cancers and human cell lines. This analysis revealed a unique form of structural variation, which we named “heterocateny” (for “variable chain”), characterized by diverse, interrelated, and repetitive patterns of concatemerized virus and host DNA segments within a tumor. Evolutionary models based on LR-seq data explained heterocateny as the consequence of aberrant host DNA replication and recombination induced by HPV integration. We conclude that HPV integration promotes intratumoral heterogeneity and clonal evolution.

## RESULTS

Our analysis of 105 HPV-positive oropharyngeal cancers by WGS identified HPV-host breakpoints that directly flank, bridge, or map within host genomic regions enriched with SVs and CNVs (5). To resolve genomic rearrangements at sites of HPV integration, here we used Illumina WGS and two forms of LR-seq, PacBio HiFi and Oxford Nanopore Technologies (ONT). These methods were chosen because they yield high resolution reads with few errors at a single nucleotide level (WGS, PacBio) or longer, continuous reads that can span across genomic features, including repetitive elements, for tens of kilobases (ONT). Given the expected lengths of ONT reads, we selected five HPV-positive primary oropharyngeal cancers and four cell lines, each with virus-host breakpoints that were mapped by WGS to target regions with CNVs ≥4n and SVs with breakpoints <60 kb apart. Cell lines included 93-VU-147T (hereafter VU147) (11), GUMC-395 (12), HeLa (13), and HTEC (14). Details about distributions of read lengths and depths of sequencing coverage are provided in Supplementary Information (**Supp Figs. 1.1-1.4, Supp Tables 1.1,1.2**).

### Extensive concatemerization and variation of HPV genomic DNA

After initial infection, HPV is maintained as an ∼7.9-kb ecDNA episome in cell nuclei. Therefore, we evaluated the technical ability of both LR-seq approaches to capture and identify small circular DNA molecules, using the ∼16.5-kb circular mitochondrial (mt)DNA genome as a proxy. Histograms of ONT mtDNA reads displayed frequency peaks at 16.5 and 33 kb (**Ext Data Fig. 1, Supp Table 1.1**). Plots of the distance between the 5’ and 3’ ends of mapped reads were within 100 base pairs (bp) in the reference mtDNA genome, indicating predominantly one- and two-unit circular mtDNA genomes. This analysis confirmed the ability of LR-seq to detect ecDNAs, determine their lengths, and identify head-to-tail tandem repeats.

Comparable analysis of ONT reads mapped to the HPV reference genome revealed read lengths frequently exceeding ∼7.9 kb (**Fig. 1A-D**, *top*, **Supp Fig. 1.5**). Plots of the distance between the 5’ and 3’ ends of mapped reads (**Fig. 1E**) indicated a predominance of single-unit HPV episomes in one primary cancer (**Fig. 1A**, *bottom*, Tumor 1) and multi-unit, head-to-tail virus-virus concatemers in others (**Fig. 1A-D**, *bottom*, **Fig. 1F, Supp Fig. 1.5**), consistent with recent reports (15, 16).

**Figure 1.**
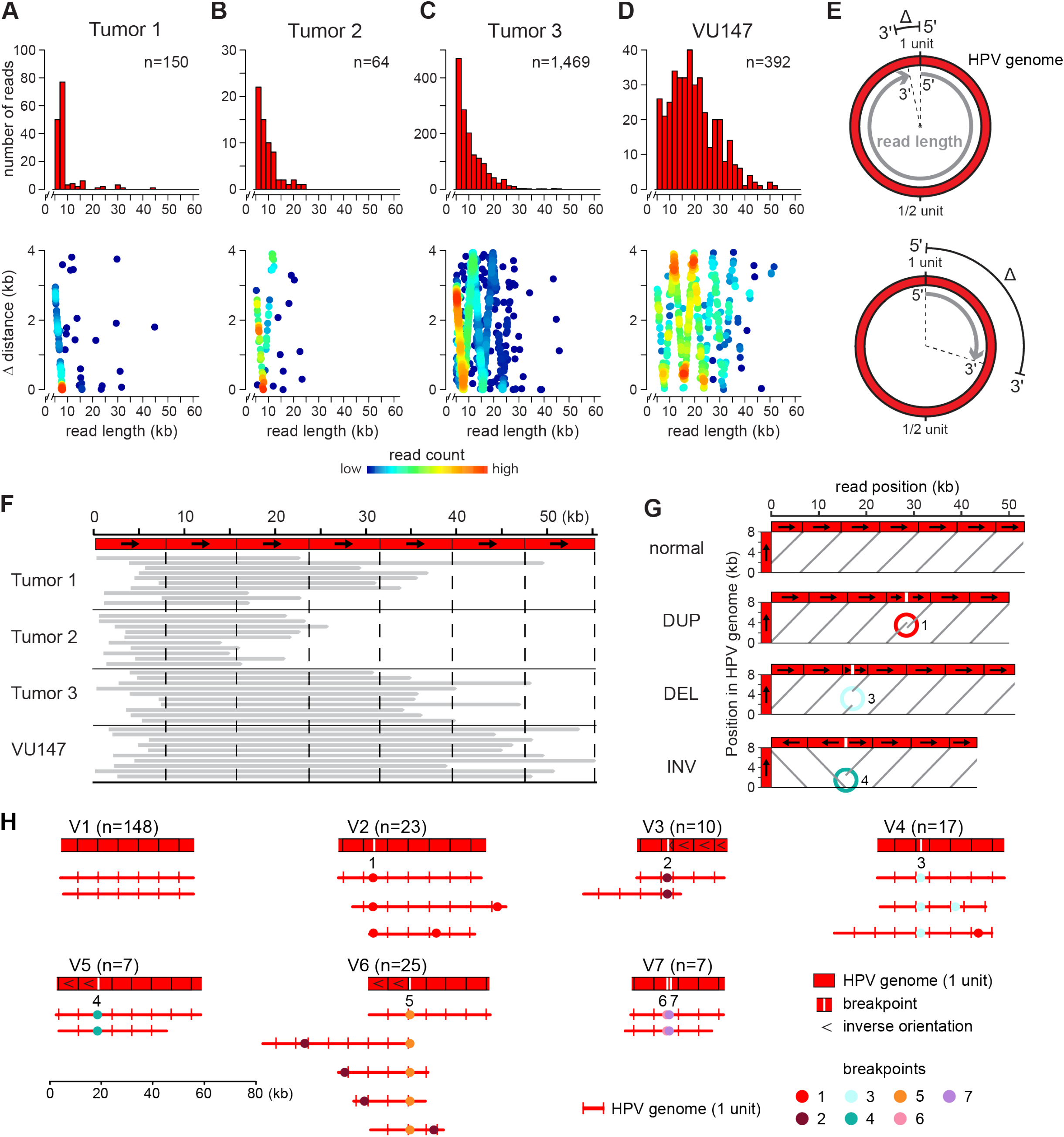
Detection of HPV concatemers and SVs by LR-seq reads. LR-seq (ONT) reads containing only HPV sequences revealed frequent HPV concatemers with and without structural variants (SVs) in multiple cancers and cell lines. (A-D) Shown are (*top, y-axis*) read count histograms and (*bottom, y-axis*) plots of the distance (Δ) between 5’ and 3’ mapped coordinates when HPV-only reads were aligned against the HPV16 reference genome for (A) Tumor 1, (B) Tumor 2, (C) Tumor 3, and (D) VU147 cell line. *X-axis, top and bottom panels*, ONT read lengths in kilobase pairs (kb); *n*, number of aligned ONT reads. *Bottom, heatmap*, read counts. (E) Schematic depicting distance Δ between read 5’ and 3’ ends (based on half-maximal genome unit circumference, 7906 ÷ 2 bp). *Grey, top and bottom*, two ONT reads aligned against (*red*) one-unit circle of the HPV16 genome. (F) Representative ONT reads from samples in panels A-D, aligned against (*x-axis, dashed lines*, ∼7.9-kb HPV genome unit length) concatemers of HPV genome. *Black arrows*, orientation of HPV genome from coordinate 1 to 7906. (G) Dotplots depict (*light gray*) alignments of *(x-axis)* representative ONT reads from VU147 cells of variable lengths against *(y-axis, arrow)* one ∼7.9-kb unit of HPV genome. *DUP*, duplications; *DEL*, deletions; *INV*, inversions. (H) Virus only VU147 ONT reads shown as (*top*) block diagrams and (*bottom*) breakpoint plots, grouped by presence of unique virus-virus breakpoints. *Red lines*, HPV genome (vertical black ticks, HPV reference coordinate 0; vertical white ticks, HPV rearrangment); *colored dots, numbers*, breakpoints (*see inset key, panel H*); *numbers below block diagrams*, group-defining breakpoints. See also **Supp Figs. 1.5,1.6**.

In contrast to mtDNA, ONT reads of HPV genomes deviated more frequently from the expected unit lengths (i.e., multiples of ∼7.9 kb; **Fig. 1A-D**, *bottom*), revealing rearrangements in virus DNA. All unique virus-virus breakpoints were confirmed by at least two and almost always by all three sequencing platforms, arguing against technical artifacts (**Supp Tables 2-5**) (17). Alignment of ONT reads from VU147, Tumor 2 and Tumor 3 against an HPV16 template model revealed rearrangements including tandem duplications, deletions, and inversions (**Fig. 1G, Supp Fig. 1.6**). The seven unique virus-virus breakpoints detected in VU147 were each assigned a numerical identifier (i.e., 1-7). To facilitate pattern recognition, DNA segments in LR-seq reads were visualized using block diagrams, and breakpoints were visualized using breakpoint plots (**Fig. 1H**). The HPV reference genome coordinate of 0 was depicted as black or red vertical lines in these and all subsequent visualizations (**Figs. 1-6**). Numerous, diverse rearrangements in HPV DNA were apparent in VU147 (**Fig. 1H**), indicating genetic instability of the concatemerized virus genomes.

### Identification of heterocateny, a unique form of structural variation

Extending our analysis beyond virus-only LR-seq reads, we found that Tumor 4 harbored a total of 22 unique breakpoints flanking regions of CNVs and SVs on Chrs. 5p13, 5q14, and Xp22, including 14 HPV-host, five host-host, and three virus-virus breakpoints (**Fig. 2A, Supp Figs. 1.1,1.2, Supp Table 2.1**). Rearrangements included two chromosomal translocations -- t(5;X)(p13;p22) and t(5;X)(q14;p22) (**Supp Table 2.1**). To facilitate resolution of genomic structural rearrangements as covered by ONT reads, the breakpoints that were best-supported by discordant or split WGS and/or LR-seq reads were selected as segment-defining breakpoints. This allowed us to delineate host or virus DNA segments based on the reference human and HPV type-specific genomes (**Supp Table 2.2**).

**Figure 2.**
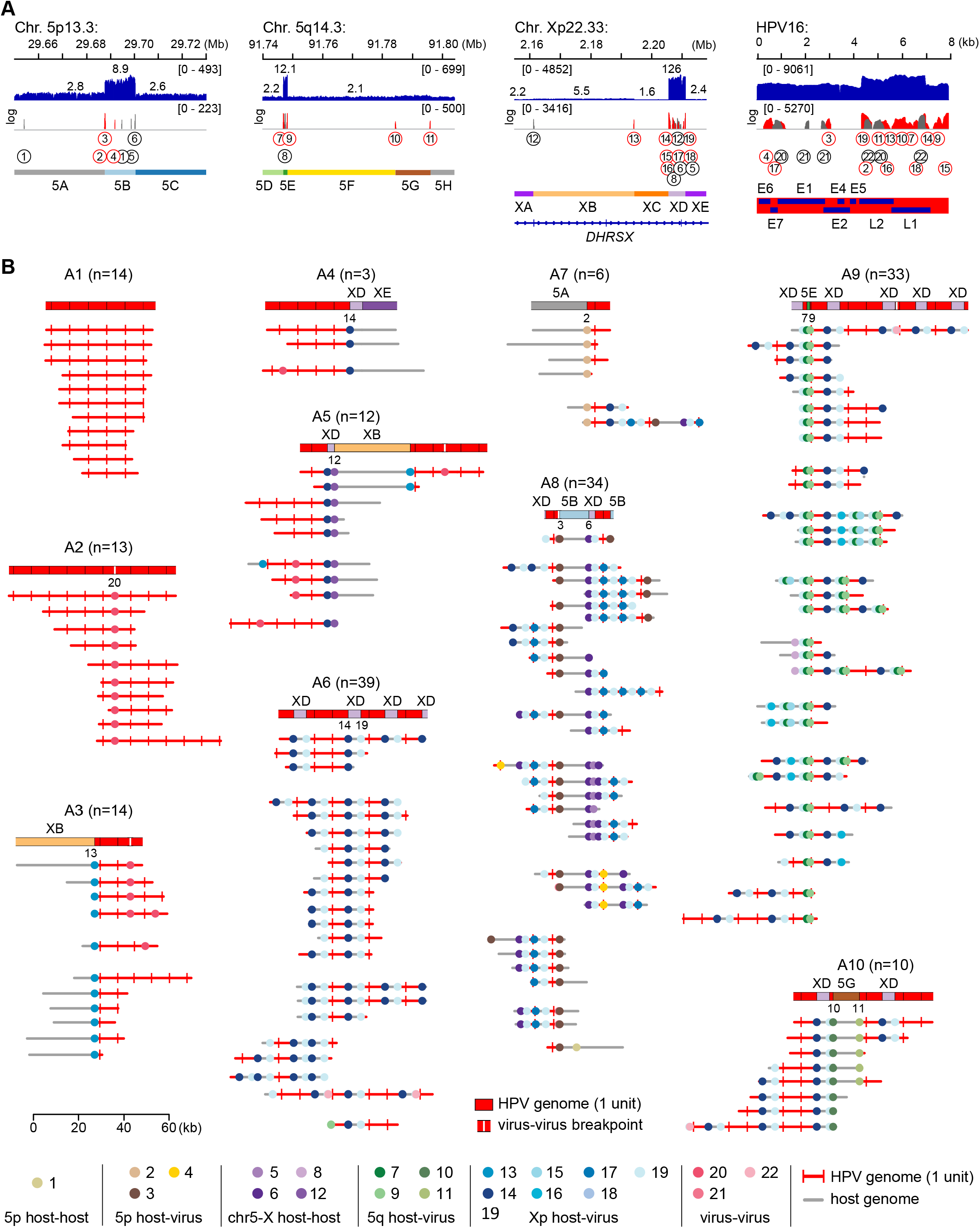
Intratumoral heterogeneity and clonal evolution induced by HPV integration. Analysis of LR-seq reads from Tumor 4 revealed shared breakpoint patterns and extensive heterogeneity in virus-virus and virus-host DNA structures. (A) Depths of sequencing coverage and breakpoints at HPV integration sites at (*left to right*) Chrs. 5p, 5q, and Xp and in the HPV16 genome, as indicated. *Top*, IGV browser display of (*y-axis, blue*) WGS coverage; *middle* (*red*) virus-host and (*gray*) host-host or virus-virus breakpoints. *Circles, numbers*, identifiers of each (*top*) segment-defining and (*bottom*) segment non-defining breakpoint (see **Supp Table 2.1**). *Bottom, left*, genomic segments defined by breakpoints (see **Supp Table 2.2**); *right*, HPV genes. (B) Pairs of schematics display (*top*) block diagrams depicting ONT reads of length >20 kb and (*bottom*) breakpoint plots, supporting groups of various HPV-host structures sharing patterns of breakpoints and genomic segments. Groups A1-A10 are defined by shared breakpoint patterns per breakpoint IDs specified below block diagrams. Breakpoint plots within groups also display further heterogeneity characteristic of heterocateny. *Red lines*, HPV genome (*vertical red ticks*, HPV reference coordinate 0; *vertical white ticks*, HPV rearrangements); *gray lines*, host DNA segments; *colored dots, numbers*, breakpoints (*see inset key, panel B*). *Parentheses*, count of reads in group.

Tumor 4 harbored ∼171 HPV16 genome copies per haploid genome, as estimated from WGS. Virus-virus concatemers comprising up to 6 tandem HPV16 genomes were detected (**Fig. 2B**, group A1), but HPV nucleotides 5144-7906 plus 1-776 were deleted intermittently from adjoining viral genome units, forming a unique, recurring virus-virus breakpoint (i.e., breakpoint 20; **Fig. 2B**, group 2, **Supp Table 2.1**). ONT reads with lengths ≥20 kb (N = 178) revealed rearranged virus-host structures in which distinct segments of Chr. X (e.g., XB, XD) and/or Chr. 5 (e.g., 5B, 5E, 5G) were inserted precisely where viral genome segments were deleted (**Fig. 2B**, groups A3-10). Individual molecules displayed specific patterns of virus and/or host DNA segments and breakpoints, which in some cases were repeated in series (**Fig. 2B**, groups A6, 9, 10). These diverse patterns were analogous to those observed in the virus-only ONT reads of VU147 (**Fig. 1H**). We clustered Tumor 4 reads into groups based on key distinct breakpoints (**Fig. 2B**, groups A1-10). Both within and between these read groups, breakpoint patterns diverged markedly, demonstrating extensive intermolecular heterogeneity. Distinct patterns of breakpoints and segments defining a group were occasionally linked with other group patterns in individual molecules. For example, breakpoint 13 in group A3 was also connected to breakpoints 12 and 14 in group A5 (**Fig. 2B**).

We used the unique breakpoints and patterns shared across heterogeneous structures in Tumor 4 as molecular barcodes to reconstruct genomic structural evolution from a common molecular ancestor (**Methods**). According to the resulting model, insertion of concatemerized HPV genomes initially occurred at the host DNA segment XC deletion site on Chr. X (**Fig. 3**). During their subsequent excision from Chr. X, these concatemerized HPV genomes captured host DNA and formed ecDNA, which then inserted into Chr. 5p and 5q (**Fig. 3**). Shared virus and host DNA segments and breakpoints were linked in series in recurrent patterns but lacked single nucleotide variants or small insertions/deletions (**Ex Data Fig. 2**), consistent with intermittent high-fidelity amplification by rolling-circle replication or recombination-dependent replication (RDR) (18-21).

**Figure 3.**
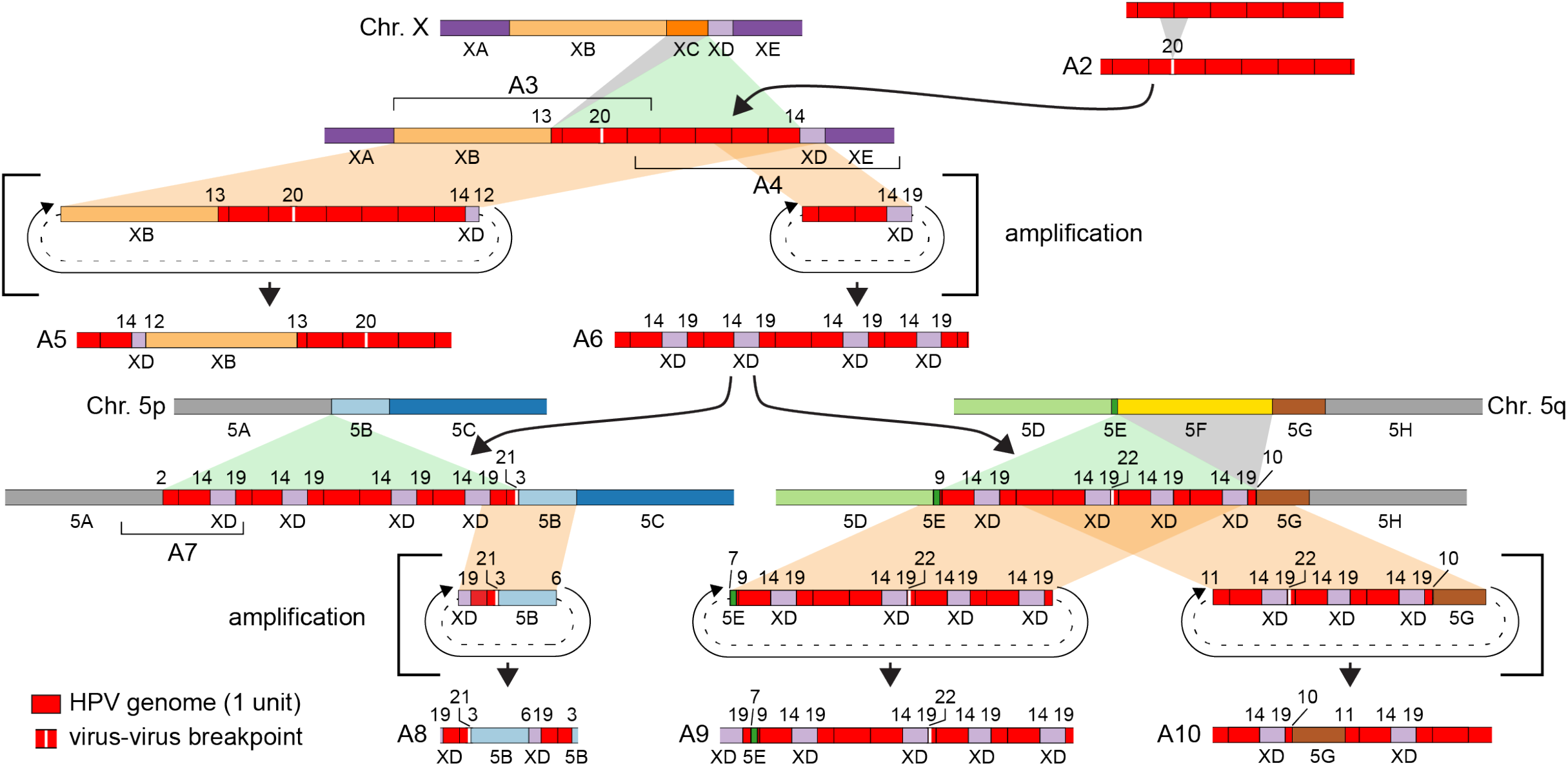
Evolution of heterocateny in Tumor 4. A model depicts how various groups of structural variants evolved from a common molecular ancestor (see **Methods**). *Block diagrams* (e.g., A1, A2, A3), representative ONT reads as in **Fig. 2B**; *brackets*, hypothetical intermediate structures; *gray*, deletions; *green*, insertions; *tan*, ecDNA excisions; *dashed lines*, circularized segments; *circular arrow*, amplification; *block colors*, segments defined in **Fig. 2A,B**.

Counts of reads supporting integration of rearranged virus-host concatemers into flanking chromosomal DNA were very low in Tumor 4. Therefore, we inferred that the numerous virus-host concatemers observed in this tumor mostly occurred as ecDNA. Notably, predictions for ecDNA as made by AmpliconArchitect software (22) were oversimplified and inaccurate compared to the virus-host concatemers we detected by LR-seq (**Supp Fig. 2, Fig. 2**), likely due to inherent limitations of short-read WGS data.

In sum, Tumor 4 LR-seq data revealed a unique form of structural variation flanking virus-host breakpoints, characterized by diverse, interrelated, and repetitive patterns of virus and host DNA segments and breakpoints (**Fig. 2B**). We named this unique form of structural variation “heterocateny.” Multiple independent lines of evidence for heterocateny were observed in all cancers and cell lines studied, as described below.

Tumor 2 harbored a total of 23 breakpoints, including 14 virus-host, four host-host, and five virus-virus, at the ∼60 kb *EP300* locus on Chr. 22q13.2 (**Fig. 4A, Supp Table 3.1**). *EP300* is frequently inactivated by somatic mutation in HPV-positive oropharyngeal cancers (23). Of the breakpoints, 14 were chosen as segment-defining breakpoints (**Fig. 4A, Supp Table 3.2**). As done for Tumor 4, we used key breakpoints to sort ONT reads into groups (**Fig. 4B**). ONT reads (N = 154) supported concatemers with multiple tandem full-length HPV genome units interspersed with tandems lacking nucleotides 7065-7906 and 1-2312 (breakpoint 17; **Fig. 4B**, group B2). Virus concatemers containing breakpoint 17 were detected in series with *EP300* segments (**Fig. 4B**, groups B3-10). Structural heterogeneity within and between read groups, analogous to that in Tumor 4 (**Fig. 2B**), provided further evidence of heterocateny. Based on LR-seq data, these structures evolved from a clonal ancestor by sequential events, including insertion of concatemerized HPV genomes, ecDNA excision, copy number amplifications, and additional rearrangements such as serial deletions (**Ext Data Fig. 3**). No WGS or LR-seq reads supported integration of virus-host structures into Chr. 22q13.2, suggesting that virus-host concatemers containing *EP300* fragments occurred predominantly or exclusively as ecDNA.

**Figure 4.**
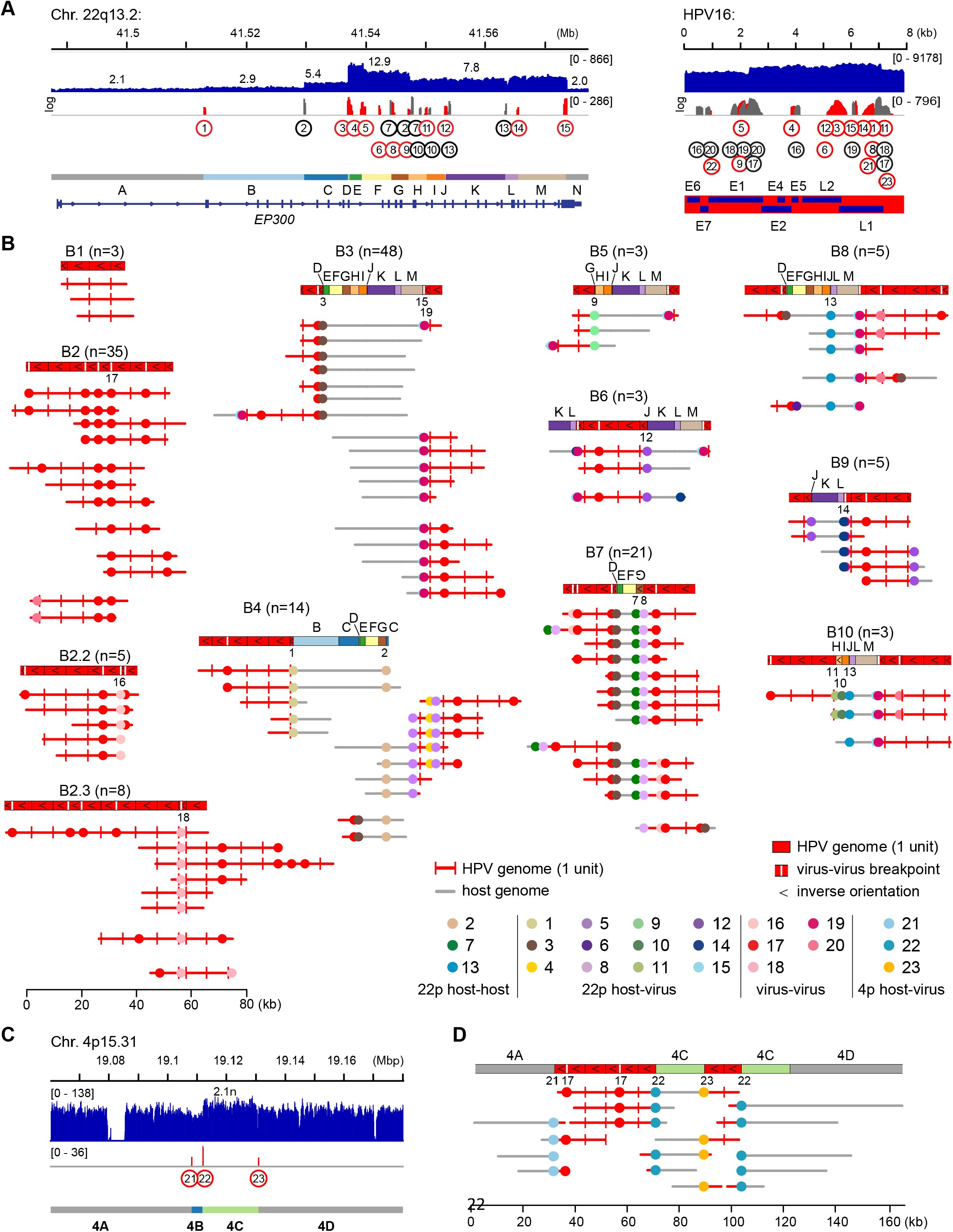
Heterocateny involving the *EP300* locus and Chr. 4p in Tumor 2. The *EP300* gene locus is disrupted by HPV integration and development of heterocateny in Tumor 2. (A-B) The EP300 locus and (C-D) Chr. 4p15, both in Tumor 2. (A) Depths of sequencing coverage and HPV insertional breakpoints at (*left to right*) the *EP300* gene locus at Chr. 22q13.2 and in the HPV16 genome, as indicated (see legend of **Figure 2**, panel A and **Supp Tables 3.1,3.2** for more details). (B) Pairs of schematics display (*top*) block diagrams depicting ONT reads of length >20 kb and (*bottom*) breakpoint plots, supporting groups of various HPV-host structures sharing patterns of breakpoints and genomic segments. Groups B1-B10 are defined by the breakpoint patterns represented. Breakpoint plots within groups also display further heterogeneity characteristic of heterocateny. (C) Depths of sequencing coverage and virus-host breakpoints at Chr. 4p15 in Tumor 2 as per panel A. (D) Block diagram (*top*) for a virus-host concatemer in icDNA in Chr. 4 supported by (*bottom*) representative LR-seq reads >20 kb in breakpoint plots. Breakpoint 17 is shared by concatemers at both chromosomal loci. *Red lines*, HPV genome (*vertical red ticks*, HPV reference coordinate 0; *vertical white ticks*, HPV rearrangements); *gray lines*, host DNA segments; *colored dots, numbers*, breakpoints (*see inset key, panel B*). *Parentheses*, count of reads in group.

Interestingly, ONT reads in Tumor 2 independently identified integration of virus-host concatemers into flanking host sequences at Chr. 4p15.31 (**Fig. 4C, Supp Table 3.1**). Detection of the same breakpoint 17 both in Chr. 4 (**Fig. 4D**) and at the Chr. 22 *EP300* locus indicated that the virus-host concatemers at these two distinct sites were clonally related. This example also demonstrated that concatemers can coexist as ecDNA and as icDNA integrants within the same tumor.

Virus-host concatemers detected in Tumor 5 mapped near or included the cancer driver gene *MYC* on Chr. 8q24.21, a hotspot for HPV integration in oropharyngeal (5) and cervical cancers (24). We identified six breakpoints, three virus-host and three host-host (**Supp Table 3.3**), which were selected to delineate host segments A through J at *MYC* (**Ex Data Fig. 4A, Supp Table 3.4**). While HPV concatemers were not detected, a deletion at HPV nucleotides 1803-2170 was identified. We detected 110 ONT reads (each ≥20 kb) defining SVs at the *MYC* locus. Of these, 98 (88%) supported a genomic rearrangement in which *MYC* was duplicated at least twice in tandem (segment E, **Ext Data Fig. 4B**). Less common but related SVs were derived from this ancestral molecule by recombination events (**Ext Data Fig. 4B**). Because no reads supported integration of virus-host concatemers into adjacent chromosomal DNA, they likely existed predominantly in ecDNA form. As this tumor harbored ∼20 HPV16 genome copies per haploid genome, each cell may have contained a range of ecDNA molecules with lengths commensurate with numbers of linked HPV units (**Ext Data Fig. 4C**). The relative homogeneity of ecDNA structure in Tumor 5, relative to Tumors 4 and 2 above, is consistent with a selective, clonal growth advantage conferred by the captured *MYC* oncogene.

In Tumor 3, single- to six-unit HPV16 episomes predominated (**Fig.1C,F**), but 5 virus-host breakpoints mapped to a gene rich region on Chr. 3q27.1 (**Ext Data Fig. 4D, Supp Tables 3.5,3.6**). LR-seq data supported insertion of a virus concatemer at this locus (**Ext Data Fig. 4E,F**), but low read counts and few derivative rearrangements (**Ext Data Fig. 4E**) suggested that ecDNA excision and recombination likely occurred in a subclonal population. Tumors 3 and 5 thus demonstrate early subclonal versus dominant clonal populations induced by HPV integration, respectively.

### Heterocateny in cancer cell lines

The GUMC-395 cell line was derived from a liver metastasis of an aggressive cervical neuroendocrine carcinoma (12). GUMC-395 cells harbored 13 breakpoints, including five virus-host and seven host-host, clustered within an ∼200-kb region of extreme hyper-amplification (up to ∼225n) and structural rearrangements at the *MYC* locus (**Fig. 5A, Supp Table 4.1**). Eight of the breakpoints defined sequential host DNA segments A - L (**Supp Table 4.2**). Segments B and C encompassed *MYC*. This cell line had ∼112 HPV copies per haploid genome.

**Figure 5.**
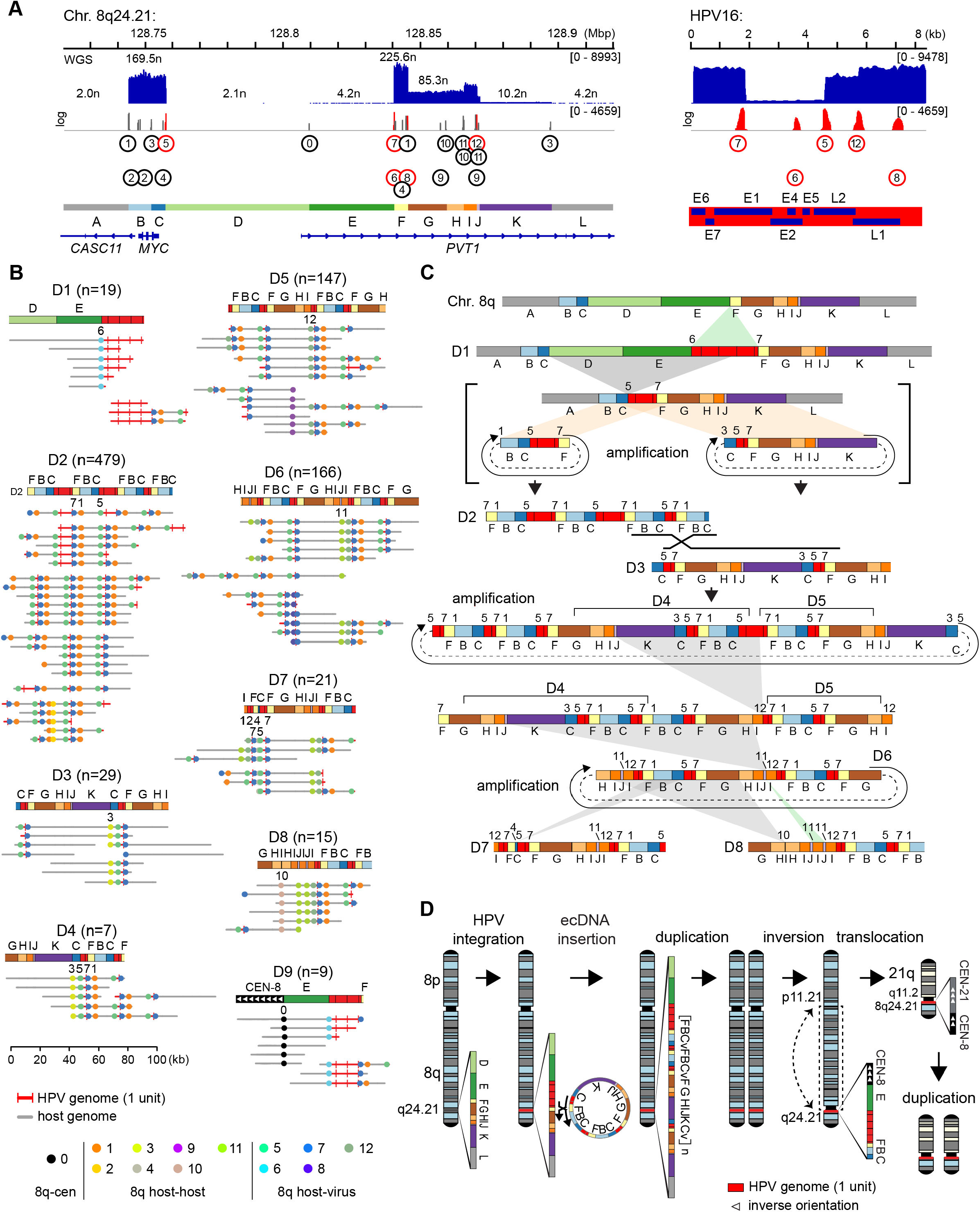
Intratumoral heterogeneity and clonal evolution at *MYC* in GUMC-395 cells. LR-seq (ONT) reads support marked heterocateny in GUMC-395 cells. (A) Depths of sequencing coverage and breakpoints at HPV integration sites at (*left to right*) Chr. 8q24.21 (*MYC* and *PVT1* genes) and in HPV16, as indicated (see legend of **Fig. 2A** and **Supp Tables 4.1,4.2** for more details). (B) *Block diagrams* (e.g., D1, D2, D3), representative ONT reads (see legend of **Fig. 2B**). Schematics of representative ONT reads of length >20 kb grouped by recurrent breakpoint patterns also display heterogeneity and incorporation of other patterns within groups. *Parentheses*, count of reads in group; *key, inset*, breakpoints and virus segments. (C) Schematic depicts potential evolution of groups of structural variants from a common molecular ancestor. *Black X*, site of potential homologous recombination. *brackets*, hypothetical intermediate structures; *gray*, deletions; *green*, insertions; *tan*, ecDNA excisions; *dashed lines*, circularized segments; *circular arrow*, amplification; *block colors*, segments defined in panel A. (D) Schematic supported by LR-seq reads depicts a stepwise model by which insertion of a virus-host concatemer containing *MYC* is followed by Chr. 8 duplication, inversion of Chr. 8q, chromosomal translocation between centromeres of Chr. 8 and Chr. 21 resulting in t(8;21)(q24;q11), and duplication of this translocation.

Analogous to our observations in primary cancers, marked heterogeneity in patterns of virus and host DNA segments and breakpoints was observed in ONT reads from GUMC-395 cells (**Fig. 5B**). Insertion of a virus concatemer was detected in Chr. 8 between segments E and F (**Fig. 5A**), defining breakpoints 6 and 7. Interestingly, no sequence data supported a normal allele connecting host DNA segments E to F (**Supp Table 4.1**), indicating loss of heterozygosity. Most virus-host concatemers identified in ONT reads (N = 774, ≥20 kb) contained breakpoint 7, nominating this insertion as an early, likely tumorigenic event. Moreover, many structural variants shared the same V-F-B-C pattern containing *MYC* and deletion of host segments D and E (**Fig. 5B**), consistent with evolution from a common molecular ancestor. In our evolutionary model for GUMC-395, ecDNAs were generated from concatemerized HPV genomes integrated at the *MYC* locus and then underwent subsequent amplification and recombination (**Fig. 5C**). These HPV-host concatemers continued to evolve via secondary recombination and deletion events (**Fig. 5C**) and ultimately gave rise to diverse but related variant structures indicative of heterocateny (**Fig. 5B**). The model provided a potential explanation for the step changes in CNVs identified from WGS data at several segment junctions, including F to G, H to I, J to K, and K to L (**Fig. 5A**). We concluded that HPV integration was responsible for hyper-amplification of *MYC* in GUMC-395, a seminal event which likely promoted the development and growth of that lethal cancer.

### Chromosomal translocations mediated by virus-host concatemers

FISH analysis of GUMC-395 cells using an HPV16 probe localized the virus to two icDNA copies of Chr. 8q and two copies of Chr. 21 in all metaphase spreads examined, due to a t(8;21)(q24.21;q11.2) translocation involving the centromere of Chr. 21 (**Ext Data Fig. 5A**). Consistent with this observation, LR-seq data showed virus-host concatemers integrated adjacent to host segment E on Chr. 8q24.21 (**Fig. 5B**, group D1) and into a second site joining host segment E to the centromere of Chr. 8 (**Fig. 5B**, group D9). In addition, numerous ONT reads that joined centromeric repeats of Chrs. 8 and 21 over several kb were detected (data not shown). We inferred that these concatemers (likely as ecDNA) were inserted by homologous recombination at the *MYC* locus, followed by Chr. 8 duplication, intrachromosomal Chr. 8q inversion, t(8;21)(q24.21;q11.2) translocation, and duplication of this translocation (**Fig. 5D**).

Numerous HeLa ONT reads supported virus-host concatemers integrated upstream of *MYC* on Chr. 8q24.21 (**Fig. 6A,B, Supp Tables 5.1,5.2**), corroborating previous analyses (25-27). A chromosomal translocation in HeLa at t(8;22)(q24;q13) was initially identified by spectral karyotyping (27) but was not detected using WGS or haplotype-resolved data (25, 26). Its relationship with HPV integration, if any, was not reported previously. Our LR-seq data uniquely confirmed and resolved this translocation (**Fig. 6C**). We identified virus-host concatemers with breakpoints identical to those integrated in Chr. 8 but connecting the 5’ end of HeLa genomic segment C with a 2-kb segment of repeated telomeric sequences (i.e., 5’-TTAGGG) on Chr. 22, forming breakpoint 2 (**Fig. 6C**). Consistent with ONT data, HPV18 FISH probes hybridized to two of three copies of Chr. 8, a t(8;22)(q24;q13) translocation, and a complex der(5)t(5;22;8)(q11;q11q13;q24) rearrangement (**Ext Data Fig. 5B**) (27). WGS data indicated that four of the five copies of Chr. 8q extended from the HPV integration site to a telomere. Thus, we inferred that virus-host concatemers first integrated into Chr. 8, followed by Chr. 8 duplication, translocation to the telomere of Chr. 22, and translocation from the Chr. 22 centromere to the Chr. 5 centromere (**Fig. 6D**).

**Figure 6.**
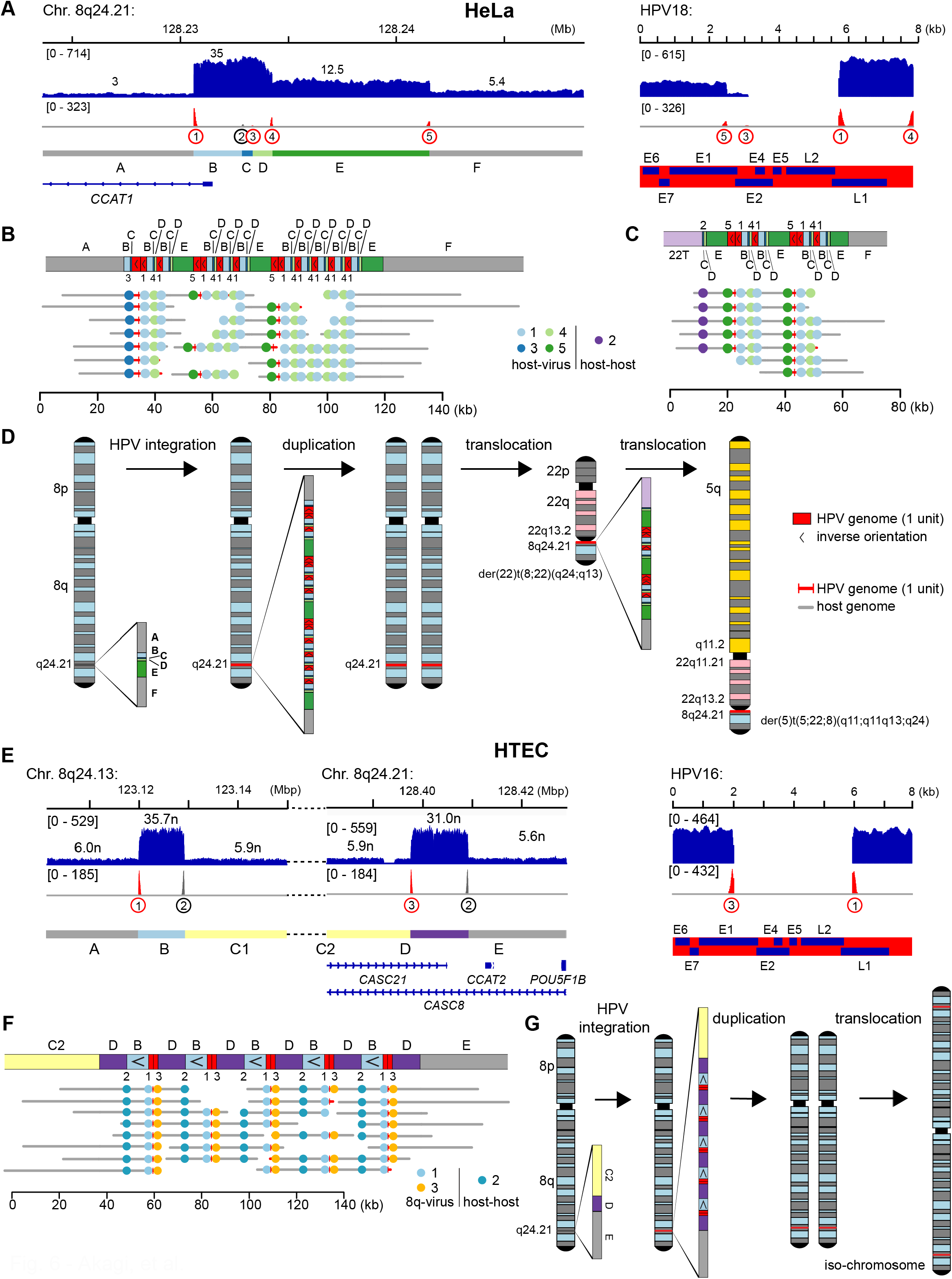
HPV integration in HeLa cells and human tonsillar epithelial cells (HTEC) induced CNV, SV, and intrachromosomal rearrangements. Virus-host concatemers in icDNA lead to chromosomal instability in HeLa (A-D) and HTEC (E-G) cells. (A) Depths of sequencing coverage and breakpoints at HPV integration sites in HeLa at (*left to right*) Chr. 8q24.21 (upstream of *MYC*, not shown) and in the HPV18 genome, as indicated (see legend of **Fig. 2A** and **Supp Tables 5.1,5.2** for more details). (B, C) *Top*, block diagrams depicting concatemerized HPV integrants and rearrangements (B) integrated into flanking intrachromosomal segments at Chr. 8q24, and (C) joining Chr. 22 and Chr. 8 at a translocation. *Bottom*, breakpoint plots depicting (*x-axis*) representative ONT reads of length >20 kb, as annotated in legend, **Fig. 2B**. Many of the ONT reads demonstrate intrachromosomal integration as they directly connect concatemers with flanking host DNA segments A (*left*) and F (*right*). *Key, inset*, breakpoints, virus segments as defined in **Supp Table 5.1**. (D) Stepwise model depicting molecular evolution of Chr. 8, starting with insertion of a virus-host concatemer (inset) into Chr. 8q24.21, likely by homologous recombination, followed by chromosomal translocation to the telomere of Chr. 22 and then to the centromere of Chr. 5. (E) Depths of sequencing coverage and breakpoints at HPV integration sites in HTEC at (*left to right*) Chr. 8q24.13 (upstream of *MYC*, not shown) and in the HPV16 genome, as indicated (see legend of **Fig. 2A** and **Supp Tables 5.5,5.6** for more details. (F) ONT reads (*bottom, breakpoint plots*) harboring breakpoints (*key, inset*) support integration of a virus-host concatemer in icDNA at Chr. 8q24.13 (G) *Left to right*, stepwise model depicting molecular evolution of Chr. 8 in HTEC *in vitro*, starting with insertion of a virus-host concatemer (inset) into Chr. 8q24.13, likely by homologous recombination, followed by chromosomal duplication and development of isochromosome 8.

Collectively, our combined analysis of HeLa and GUMC-395 cells revealed that integrated virus-host concatemers are unstable and can induce chromosomal translocations and other forms of genomic structural variation.

### HPV integrants were detected in icDNA and ecDNA in cell lines

The GUMC-395 and HeLa ONT data supported integration of virus-host concatemers into icDNA. In contrast, VU147 ONT data revealed virus-host concatemers containing Chr. 17q12 segments in ecDNA form (**Ext Data Fig. 6A-C, Supp Tables 5.3,5.4**) and virus-host concatemers anchored into icDNA at Chr. Xp21.1 (**Ext Data Fig. 6D,E**). To evaluate the possible occurrence of virus-host concatemers in ecDNA form in GUMC-395, HeLa, and VU147, we performed metaphase fluorescence *in situ* hybridization (FISH) with HPV probes and Circle-seq. Both methods identified HPV-containing ecDNA in subsets of cells in all cell lines examined (**Ext Data Fig. 7**). GUMC-395 and HeLa Circle-seq data supported ecDNA from the *MYC* locus on Chr. 8q24.21 that aligned well to their respective amplified regions (**Ext Data Fig. 8A,B**). Comparable data were observed for VU147 (**Ext Data Fig. 8C,D**). This analysis confirmed structurally similar virus-host concatemers in ecDNA and icDNA forms in cell lines, indicating excision from and insertion into chromosomes.

**Figure 7.**
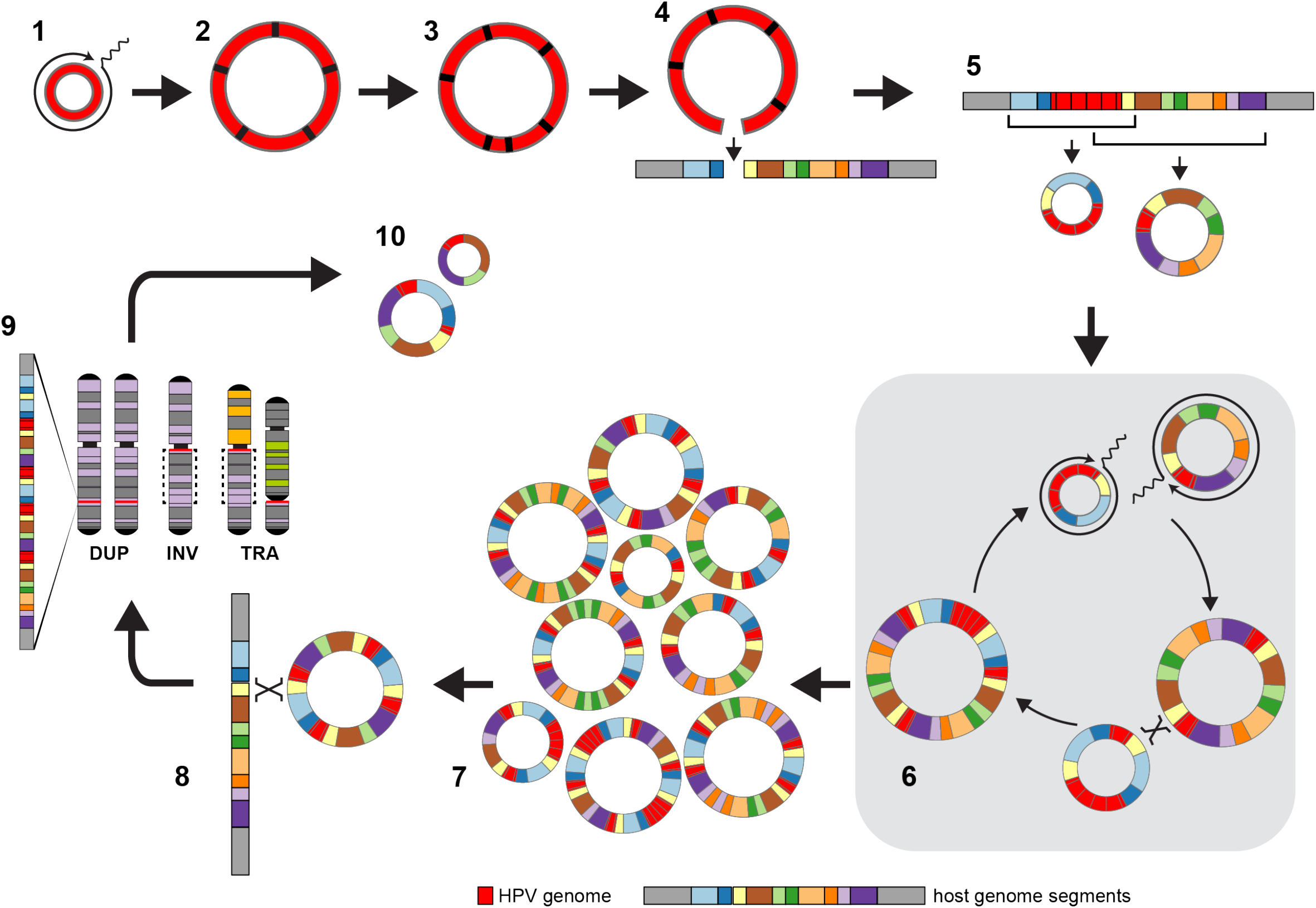
Model of HPV heterocateny, derived from multiple lines of evidence. A general model for development of HPV heterocateny with highly diverse but related genomic rearrangements, including CNV and SV, at HPV integration sites. (1) Rolling-circle replication of HPV episomes results in (2) unstable virus genome ecDNA concatemers that (3) acquire structural rearrangements and (4) integrate into chromosomes at sites of double-strand DNA breaks. (5) Dynamic excision of virus with captured host DNA leads to (6) serial rounds of amplification of ecDNA by rolling-circle or recombination-dependent replication and recombination events between host and/or HPV segments in the same cells, driving (7) HPV heterocateny and thus intratumoral heterogeneity and clonal evolution. (8) Insertion of ecDNA by recombination into chromosomes (likely through homology-directed repair) can induce (9) chromosomal inversions (INV) and translocations (TRA). (10) Occasional additional rounds of excision may produce more diverse HPV ecDNAs.

### HPV integration *in vitro* recapitulates clonal selection at the *MYC* locus

The human tonsillar epithelial cell line (HTEC) was created upon transfection of primary cells with HPV16 episomal DNA *in vitro*, followed by clonal selection in cell culture (14). Virus integration and formation of HPV-host concatemers occurred solely during cell culture *in vitro*. LR-seq data revealed striking similarities between HPV integration sites and genomic rearrangements at the *MYC* locus in HTEC and those in both GUMC-395 and HeLa cells. In HTEC, two virus-host breakpoints flanked the 5’ ends of two amplified genomic loci (i.e.,16-19n) ∼350 kb upstream of *MYC* (**Fig. 6E, Supp Tables 5.5,5.6**), analogous to HeLa in location (**Fig. 6A**) and to GUMC-395 in structural variation (**Fig. 5A**). ONT reads demonstrated integrated virus-host concatemers that displayed homology to host DNA segments captured in the concatemer (**Fig. 6F**), supporting a mechanism of insertion induced by homologous recombination comparable to that of HeLa (**Fig. 6B, Ext Data Fig. 5D,E**). In serially passaged HTEC cells, Circle-seq reads aligning at this locus showed structural variation and additional discordant rearrangements, suggesting instability of intrachromosomal insertions resulting in occasional excisions of ecDNA from this site (**Ext Data Fig. 8E**). HPV16 FISH probes localized to Chr. 8q and to both ends of isochromosome i(8q) in all metaphase spreads examined (**Ext Data Fig. 5C**, indicating viral integration preceded formation of this chromosomal abnormality as the epithelial cells evolved *in vitro* (**Fig. 6G**).

### HPV genomic structures and transcripts in the context of heterocateny

Virtually all primary tumor and cell line ONT reads containing HPV sequences included at least one copy of the viral origin of replication (HPV16 nucleotides 7838-7906 and 1 to 100) and the region encoding E6/E7 (nucleotides 83 to 858), even when other HPV genomic deletions were observed. RNA-Seq analysis revealed high levels of E6 and E7 transcripts in all cases (**Ext Data Fig. 9**) (23). Except for Tumor 5, in which E1 was deleted, the primary tumors with a predominance of virus-host concatemers in ecDNA form contained full-length HPV genomes and expressed E1 and E2 transcripts. In contrast, the cell lines with a predominance of virus-host concatemers in icDNA, i.e., HeLa, GUMC-395, and HTEC, had deletions in E1 and/or E2 and the corresponding transcripts were poorly expressed. Thus, E6 and E7 were expressed regardless of whether E2 had been disrupted.

## DISCUSSION

Here we identified heterocateny, a unique form of structural variation induced by HPV integration in human cancers, characterized by highly diverse, interrelated, and repetitive patterns of virus and host DNA segments and breakpoints that coexist within a tumor. We detected strong evidence of heterocateny in HPV-containing ecDNA, icDNA, or both across all cancers and cell lines evaluated. Evolutionary models based on LR-seq data explained heterocateny as the consequence of aberrant host DNA replication and recombination, induced by HPV integration and frequently involving concatemerized, circularized DNA. We inferred that virus-virus and virus-host genomic structural rearrangements characteristic of heterocateny are unstable, whether present in ecDNA or icDNA, leading to further intratumoral heterogeneity and clonal evolution. For this reason, we also use the term heterocateny to describe the stepwise process by which HPV induces formation of this unique form of genomic heterogeneity.

Our previous WGS analyses of cell lines (2) and primary tumors (5) prompted us to develop the mechanistic “looping” model to explain extensive genomic structural variation observed at HPV integration sites (2). Our HPV looping model proposed that double-stranded HPV DNA breaks facilitate capture of host DNA, resulting in insertional breakpoints followed by amplification, recombination, repair, and integration of virus-host concatemers in icDNA (2). However, short WGS reads limited our ability to connect genomic segments and breakpoints over longer genomic distances. The new insights gained here have enabled expansion of this HPV looping model to include generation and insertion of unstable, concatemerized HPV genomes into icDNA; capture and rearrangement of host DNA during excision of HPV ecDNAs from icDNA and their insertion back into icDNA; HPV-host segment amplification by rolling-circle replication or RDR; recombination between repetitive or homologous segments, likely by homology-directed repair or template-switching during replication, resulting in novel combinations of breakpoint and segment patterns; and formation of chromosomal inversions and translocations between repeats in concatemers and telomeres and/or centromeres (**Fig. 7**).

Steps 1-5 in our evolutionary model (**Fig. 7**) are consistent with the existing literature. For example, Southern blotting and two-dimensional electrophoresis provided low-resolution evidence of integrated and/or episomal HPV concatemers in cervical cancers and cell lines (28, 29). Excision of HPV integrants from icDNA after unlicensed replication from the HPV origin was proposed after analysis of short-read WGS data from TCGA (30). Although virus-host concatemers (2) and hybrid episomes (30) were reported previously, the discovery and characteristics of heterocateny as illustrated in steps 6-10 of our model (**Fig. 7**) are novel.

Cancer evolution involves two essential processes: genetic variation and clonal selection (31). Comparisons of LR-Seq reads, as visualized in breakpoint plots in **Figs. 2B, 4B, and 5B**, for example, demonstrated extensive intratumoral genomic structural variation directly linked with HPV integrants. Our evolutionary models strongly implicated these HPV integrants as the inducers of heterocateny across individual cells and subclones in each tumor. Collectively, our data and resulting models describe the selection of DNA segments containing a host oncogene or its regulatory elements by HPV integrants, e.g., *MYC* in Tumor 5 and cell lines GUMC-395 and HeLa, in addition to the viral oncogenes expressed in all HPV-positive cancers. Similarities between the structural variants observed at the *MYC* locus in HTEC, which was immortalized and clonally selected upon transfection with HPV16 *in vitro*, and those in Tumor 5, HeLa, and GUMC-395 provide consistent and compelling experimental evidence implicating hetercateny as a driver event in the evolution of human tumors.

Overall, virus-virus and virus-host concatemers in ecDNA form showed more extensive heterocateny than those anchored into icDNA, implicating circular forms of ecDNA in the generation of heterocateny. Across primary tumors, several chromosomal loci affected by HPV integration lacked LR-seq support for anchoring into icDNA, suggesting they harbored ecDNA predominantly. In contrast, FISH analysis of cell lines demonstrated HPV integration in chromosomal DNA in every cell examined (2). Nevertheless, FISH, LR-seq and Circle-seq data from cell lines consistently suggested that integrated virus-virus and virus-host concatemers occasionally undergo excision, forming HPV ecDNAs. Distinctions observed between primary cancers and cultured cells may be attributable to differences in numbers of ecDNAs maintained in different cellular contexts. For example, essential factors required to replicate and maintain HPV ecDNA may be downregulated or lost upon derivation and growth of cell lines *in vitro*. Alternatively, primary cancer subclones harboring icDNA HPV integrants may benefit from selective growth advantages during cell line development.

We note both similarities and differences between HPV-containing ecDNAs and HPV-negative ecDNAs observed in neuroblastoma (32), glioma (33), and other cancers. The latter ecDNAs comprise very large (>1 megabase pair) circles (34) with unknown mechanisms of replication (35). Like HPV-containing ecDNAs, they frequently contain cellular oncogenes (e.g., *MYC, EGFRVIII*) (33, 36). Such ecDNAs can increase intratumoral heterogeneity and facilitate rapid adaptation to selective environmental pressures, attributed to unequal replication and segregation of ecDNAs in daughter cells during mitosis (32, 33, 36, 37). In contrast, HPV-containing ecDNAs have the viral origin of replication and encode viral proteins including oncoproteins E6 and E7. These features may increase their stable maintenance as ecDNAs by facilitating replication, segregation, and tethering onto chromosomes during mitosis (38, 39).

Loss of HPV-containing ecDNAs would likely undergo strong negative selection because expression of E6 and E7 is necessary for the malignant phenotype.

HPV undergoes two predominant modes of replication that depend upon the differentiation status of the infected cell (40-42). Maintenance replication in the basal epithelium occurs in S phase by bidirectional theta replication initiated from the viral origin and depends upon HPV E1 helicase and E2 transcriptional regulatory proteins. In contrast, rolling-circle replication and RDR occur in the G2/M phase, are less dependent on the viral origin, and are unidirectional (18, 40, 41). The latter two modes of replication depend upon E7- or E1-induced activation of the ATM-mediated DNA repair pathway (43). The virus-virus and virus-host concatemers observed here, which lacked SNVs or indels at the unit junctions (**Ext Data Fig. 2**), likely resulted from E6/E7 expression, abrogation of the G1-S checkpoint, prolonged stalling of the cell cycle in G2, and rolling-circle replication or RDR (18).

Each primary cancer and cell line analyzed here provided a snapshot in time to inform our model for heterocateny (**Fig. 7)** (2). We acknowledge a lack of longitudinally collected cancers and data to validate the sequence of events. To date, we have not demonstrated that HPV ecDNA-mediated amplification of host oncogenes contributes directly to cancer formation or progression (5). Furthermore, despite many advantages over WGS data, including longer read length distributions and continuous sequences, LR-seq data still cannot determine whether the heterogeneous, repetitive virus-host concatemerized structures detected here were linked within the same, very long (>100 kb) molecules, co-existed within the same cells, and/or were segregated among distinct subclones. Our rigorous requirement for validation by supporting reads from at least two of three independent sequencing platforms may have underestimated the extent of molecular heterogeneity in each cancer.

The model shown in **Fig. 7** proposes mechanisms by which HPV integration induces formation of CNVs and SVs, extensive diversity, and heterocateny. We conclude that this structural variation is caused by HPV integration and does not reflect a preference for HPV integration at sites of pre-existing SVs and CNVs. Our data extend our understanding of the consequences of HPV integration to include promotion of intratumoral heterogeneity and clonal evolution in human cancers.

## METHODS

### Cancer cell lines and primary tumors

HeLa, 93-VU-147T (VU147), GUMC-395 and HTEC cell lines were obtained from ATCC, Drs. RD Steenbergen, Richard Schlegel, and John Lee, respectively. Primary oropharyngeal cancer specimens were obtained with informed consent from human subjects enrolled in a genomics study at Ohio State University and studied under approved Institutional Review Board protocols (OSU, MDACC) as described (5, 23).

### Sequencing libraries and data generation

Genomic DNA was extracted from cancer samples as previously described (23). For WGS, all samples were prepared for 2 × 150 bp paired end libraries for Illumina WGS sequencing (5).

For LR-seq libraries, molecular weight distributions of genomic DNA samples were evaluated using a Femto Pulse pulse-field capillary electrophoresis system (Agilent). To prepare PacBio libraries, genomic DNA was sheared with a Megaruptor (Diagenode) or Covaris g-tube to obtain >15-25 kb fragments. Resulting sheared DNA fragments were re-assessed using the Femto Pulse. Up to 5 μgs of DNA was used to prepare a SMRTbell library with a PacBio SMRTbell Express Template prep kit 2.0 (Pacific Biosciences of California). Briefly, single-stranded DNA overhangs were removed, DNA damage was repaired by end-repair and A-tailing, PB adapters were ligated, desired size fragments were purified using AMpure PB beads, and resulting CCS HiFi libraries were sized-selected in the 10-50 kb fragment range using a Blue Pippin system (Sage Science). LR-seq data were generated on one SMRT cell 8M with v2.0/v2.0 chemistry on a PacBio Sequel II instrument (Pacific Biosciences) with movie length of 30 hours. Circular consensus sequence (CCS) data files and high accuracy subreads were generated using SMRTLink software, v. 9.0.0 to 10.1.0. If yield was < 10x fold coverage, additional library aliquots were re-sequenced.

For ONT libraries, samples containing high molecular-weight DNA fragments were sheared by passage 2-5 times (depending on starting material size distribution) through a 26.5-gauge needle. DNA size distributions were assessed again with Femto Pulse. Five μgs of DNA were used to prepare each ONT library with an Oxford Nanopore SQK-LSK-110 Kit. Libraries were size-selected to remove shorter fragments using a Short Read Eliminator (SRE) kit (Circulomics). Sized libraries were sequenced on a PromethION 24 cell PROM0002 instrument for 3 days, including a nuclease flush performed at 24 h to increase yield. Base-calling, trimming of adapters and quality checking were performed using Guppy (Oxford Nanopore), resulting in FASTQ files.

To prepare Circle-seq libraries from cultured cancer cells, we followed a published protocol (44). Briefly, 5 mcg genomic DNA was purified from serial passages of each cell line by proteinase K digestion and phenol/chloroform extraction. DNA was treated with 0.2 units/ul Plasmid-Safe ATP-Dependent DNase (Epicentre) for 5 days at 37 degrees. A SYBR Green quantitative (q)PCR (Thermo Fisher Scientific) assay of a 173bp *HBB* amplicon and TaqMan qPCR (Life Technologies) assay of a 153bp *ERV3* amplicon were used to confirm degradation of linear chromosomal DNA (i.e., expected cycle threshold values >35). Remaining circular DNA was amplified by Multiple Displacement Amplification using φ29 DNA polymerase and random hexamer primers using the Qiagen REPLI-g Mini Kit (Qiagen). Magnetic bead-based purification was used to remove the polymerase and primers. Amplified circular DNA was sheared with ten cycles (on/off, 30/30) using a Bioruptor Pico with a cooler (Diagenode). Sequencing libraries were prepared using a NEBNext DNA Library prep kit (New England Biolabs) resulting in a target insert size of 250 bp as confirmed by TapeStation (Agilent). Resulting DNA libraries were pooled at 10 nM and sequenced in 2 × 76-bp format (Illumina), resulting in >35 million read pairs per library.

### Bioinformatics analysis of sequencing data

#### Global sequence alignment and analysis

WGS data (Illumina) were aligned against a hybrid human-HPV reference genome comprised of GRCh37 + 15 high-risk HPV type genomes (GRCh37 + HPV) (5). SVs and breakpoints were detected as described (45). We previously validated our WGS pipeline for virus-host breakpoint calls with Sanger Sequencing, which confirmed ∼100% (2,5).

PacBio and ONT reads were aligned globally against a hybrid GRCh37 + HPV16 reference using Minimap2 version 2.17 (46), as part of PRINCESS version 1.0 (47). We selected default options appropriate to each sequencing platform (-x map-pb and -x map-ont, respectively). For HeLa cell analysis, we used a hybrid GRCh37 + HPV18 reference. Resulting alignments were compared against those from LRA version 1.3.2 (48), based on the same hybrid reference genomes indexed using the commands lra global, with lra align and option - CCC for PacBio HiFi data and with -ONT for ONT data. Comparable results were observed. SVs were identified from these global alignments using Sniffles v1.0.12 (49) with or without a VCF file generated by Lumpy analysis of WGS short reads (option –Ivcf). This step identified reads covering target regions of interest including clustered HPV insertional breakpoints (**Supp Tables 2-5**).

#### Local realignments and analysis

Breakpoints (i.e., virus-virus, virus-host, or host-host) that were detected with ≥20 Illumina short reads, ≥5 PacBio, and/or ≥5 ONT reads, and called by two or more of these platforms, were selected for further analysis. We defined boundaries of genomic segments by identifying sites of copy number transitions or discontinuous read alignments. Particular breakpoints that were best-supported by discordant or split WGS and/or LR-seq reads were selected as segment-defining breakpoints to delineate host or virus DNA segments based on the reference human and HPV type-specific genomes. By contrast, other breakpoints included those that did not flank a copy number transition site or were <1 kb from a segment-defining breakpoint due to alignment constraints (**Supp Tables 2-5**).

Target regions of interest were defined by alignment of virus-host breakpoints against the human reference genome, and then we added +/-50 kb of flanking genomic sequences. For local realignments, we extracted long reads that aligned in part or in total to the target regions. To facilitate local alignment, target regions of interest were extended by adding 1 Mbp of reference sequences up- and downstream (referred to as “pad” in **Supp Tables 2-5**). We used these coordinates to create a local reference sequence model for each sample locus as template for local re-alignments. Genomic coordinates of segments used for local realignments are listed in **Supp Tables 2-5**.

Realignments of extracted long reads against extended target regions were performed using Minimap2 (46). Reads with at least one segmental alignment > 1 kbp were included for further analysis. SVs in the realigned long reads were confirmed using Sniffles by alignment with these custom local sequence models (**Supp Tables 2-5**). Further local realignments were evaluated using a custom script to count numbers of long reads supporting individual segments and/or breakpoint junctions. Local realignments and qualities were visualized in alignment dotplots (e.g., **Fig. 1**) generated using pafR package (https://github.com/dwinter/pafr).

#### Reconstructing clonal evolution of virus-host concatemers and rearrangements

This analysis was restricted to informative ONT reads ≥20 kb in length that contained HPV and host DNA segments and breakpoints in a target region of interest. All breakpoints and segments in each read and their order in sequence were annotated. Further analysis was restricted to annotated patterns supported by three or more reads. To facilitate manual curation, DNA segments in LR-seq reads were visualized using block diagrams and breakpoints were visualized using breakpoint plots. LR-seq reads were then sorted into groups based upon differences in annotated patterns of segments and breakpoints.

To elucidate how annotated patterns in grouped LR-Seq reads from target regions of interest may be interrelated, grouped LR-Seq reads were serially ordered based upon the minimal number of additional DNA segments or breakpoints present when compared to the previous and subsequent group. The analysis was repeated until all LR-Seq groups from target regions of interested were included.

After the grouped LR-Seq reads were ordered, differences in annotated patterns and genomic coordinates between groups were manually inspected at single-base pair resolution, using breakpoints as molecular barcodes, to infer a mechanism by which one group could be derived from the previously ordered group with the minimal number of events, including deletion, insertion, inversion, ecDNA excision, amplification by rolling-circle or recombination-dependent replication, recombination, or translocation. We applied this examination within and across ordered groups of LR-Seq reads. This analysis was predicated upon a reasonable statistical assumption that a unique individual breakpoint occurred only once in time and would remain in downstream genomic structures unless they were deleted. Such a deletion would result in a novel breakpoint, prompting us to trace its molecular lineage. For some models, hypothetical intermediate structures were proposed to explain stepwise evolution of breakpoint patterns observed in LR-Seq reads. The sequence of inferred, ordered events were then used to create evolutionary models for each tumor or cell line.

#### Bioinformatics analysis for ecDNA detection using Circle-seq data

To increase the accuracy of structural variant (SV) detection, we merged paired-end reads having ≥15 nt overlap between them to form longer, continuous single reads using BBMAP (https://sourceforge.net/projects/bbmap/) before alignment. Resulting merged reads were aligned to human reference genome GHCh37 + HPV16/18 genome by BWA v0.7.17 (50). SVs including duplications were called by Lumpy v 0.3.0 (45). Candidate circular DNAs were detected by the following criteria: SVs (duplications as a marker of circular DNA) with ≥2 supporting reads; 95% coverage of regions flanked by SVs; and the mean depth of sequencing coverage in the amplified SV region was greater than that in the flanking region of the same length (51).

#### Prediction of ecDNA and rearrangement structures by AmpliconArchitect (AA)

We used 20x coverage Illumina paired-end WGS data as input for AmpliconArchitect (v1.2) (22). First, reads were aligned against human GRCh37 + HPV reference genome using BWA, and highly amplified regions were selected using amplified_intervals script (option --gain: 4n, --cnsize: 1000 bp). We ran AmpliconArchitect using both EXPLORE mode and VIRAL mode and comparatively predicted virus-associated amplicons. We also ran AmpliconArchitect on virus-associated amplification regions using VIRAL_CLUSTERED mode for further resolution. Amplicon types were annotated using AmpliconClassifier (v0.3.8) and amplicons predicted as ecDNA-like circular structures were visualized using CycleViz (0.1.1).

### RNA-seq analysis

Total RNA was extracted and strand-specific RNA-seq libraries were prepared and sequenced as previously described (23). RNA-seq reads (2 × 150 nt) were aligned against a custom, hybrid genome comprised of human GRCh37 reference with 13 appended HPV type genome sequences (2) using STAR aligner version 2.7.2 (52). For HPV transcript analysis, we calculated mean depth of coverage every 10 bp along the HPV16 or -18 reference genomes (NC_001526.3 and NC_001357.1) and normalized against total aligned read count per million.

### Fluorescence in situ hybridization (FISH)

Metaphase chromosomes were prepared from cultured cells by incubating them in 0.02 mg/ml Colcemid (Invitrogen; Grand Island, NY) for ∼2 h. Cells then were incubated in hypotonic (0.075M) KCl solution and fixed in methanol/acetic acid (3:1). Slides were incubated at 37°C before FISH was performed. Biotinylated HPV probes were purchased from Enzo Life Sciences. Whole chromosome paint probes were generated in-house using PCR labeling techniques (53). To increase the signal of the HPV probe, the Tyramide SuperBoost kit (ThermoFisher Scientific) was used during detection. Slides were imaged on a Leica DM-RXA fluorescence microscope equipped with appropriate optical filters (Chroma) and a 63X fluorescence objective. Slides then were counterstained with 4′,6-diamidino-2-phenylindole (DAPI) or with YOYO-1 (ThermoFisher). When HPV probe signal co-localized with YOYO-1 signal detecting DNA at 63x magnification, HPV-containing ecDNA was counted. In a proof-of-principle experiment, 293T cells were transfected with a pGEM-T vector containing or lacking full-length HPV16. Colocalized HPV and YOYO-1 DNA signals were observed only when cells were transfected with HPV, but not when empty vector DNA was used in the transfection (**Ext Data Fig. 7A**).

## Date availability

Both Illumina WGS data and LR-seq data were deposited at European Genome Archive (EGA; https://ega-archive.org/). The accession numbers are EGAD00001009630, EGAD00001009631, and EGAD00001009632.

## ACKNOWLEDGMENTS

The authors thank the cancer patients who enrolled on our genomics study, and members of the Gillison and Symer laboratories for helpful comments. The authors acknowledge computational resources from the High Performance Computing for Research facility at the University of Texas MD Anderson Cancer Center.

---

**Extended Data Figure 1.**
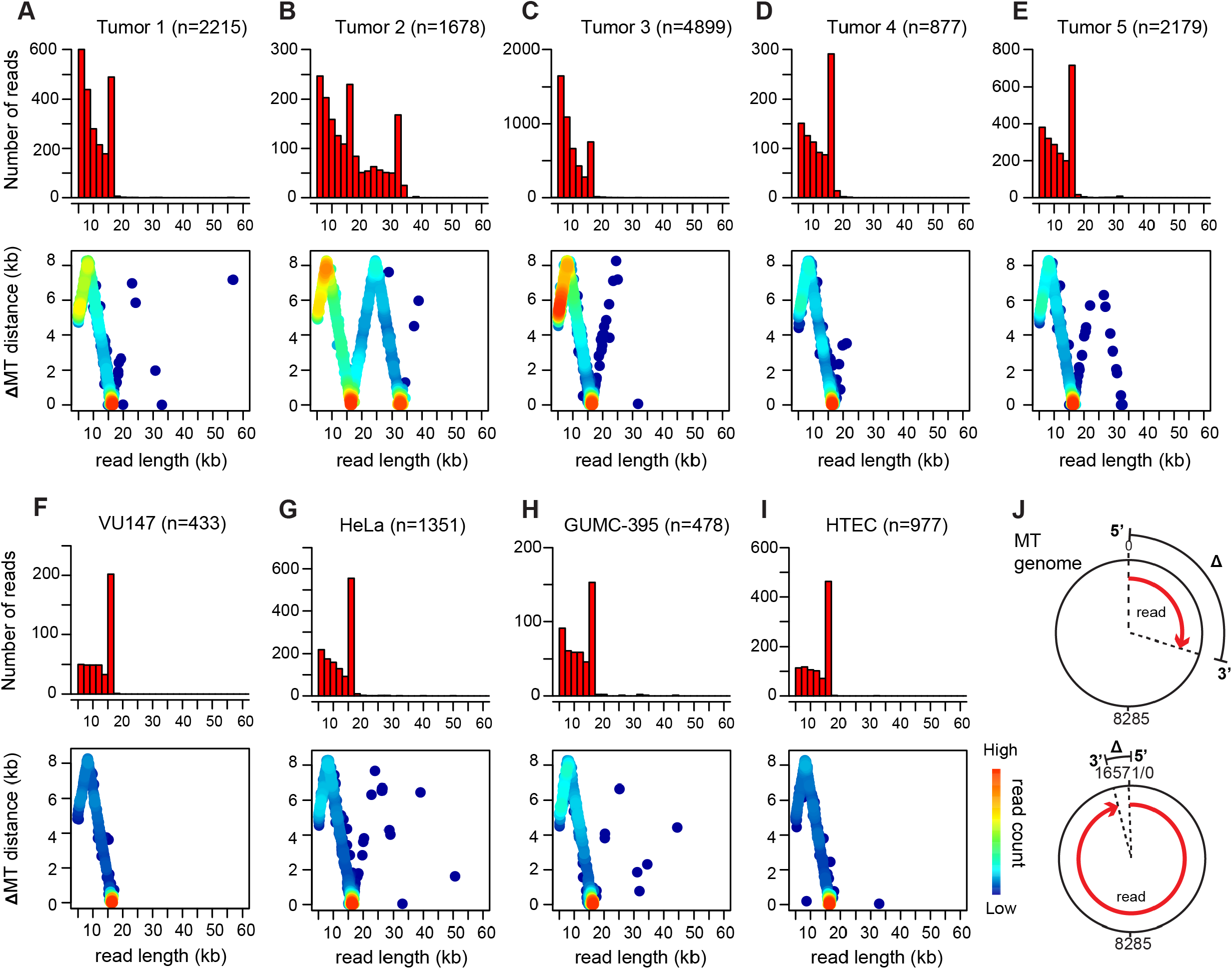
Detection of circularized mitochondrial DNA by LR-seq reads. We used the ∼16.5-kb circular mitochondrial (mt)DNA genome to evaluate the technical ability of ONT LR-seq to capture and identify small circular DNA molecules. (A-I) Shown are (*top, y-axis*) read count histograms and (*bottom, y-axis*) distance (Δ) between 5’ and 3’ coordinates of read ends when aligned to the mtDNA genome for (A) Tumor 1, (B) Tumor 2, (C) Tumor 3, (D) Tumor 4, (E) Tumor 5, and cell lines (F) VU147, (G) HeLa, (H) GUMC-395, and (I) HTEC. *X-axis, top and bottom panels*, ONT read lengths in kilobase pairs (kb); n, number of aligned ONT reads; *heatmap*, read counts. (J) Schematic depicting distance (Δ) between read 5’ and 3’ ends (based on half-maximal genome unit circumference, 16,569 ÷ 2, bp). *Red, top and bottom*, two ONT reads aligned against (*black*) one unit circle of mtDNA genome. See **Table S1.1** for additional details.

**Extended Data Figure 2.**
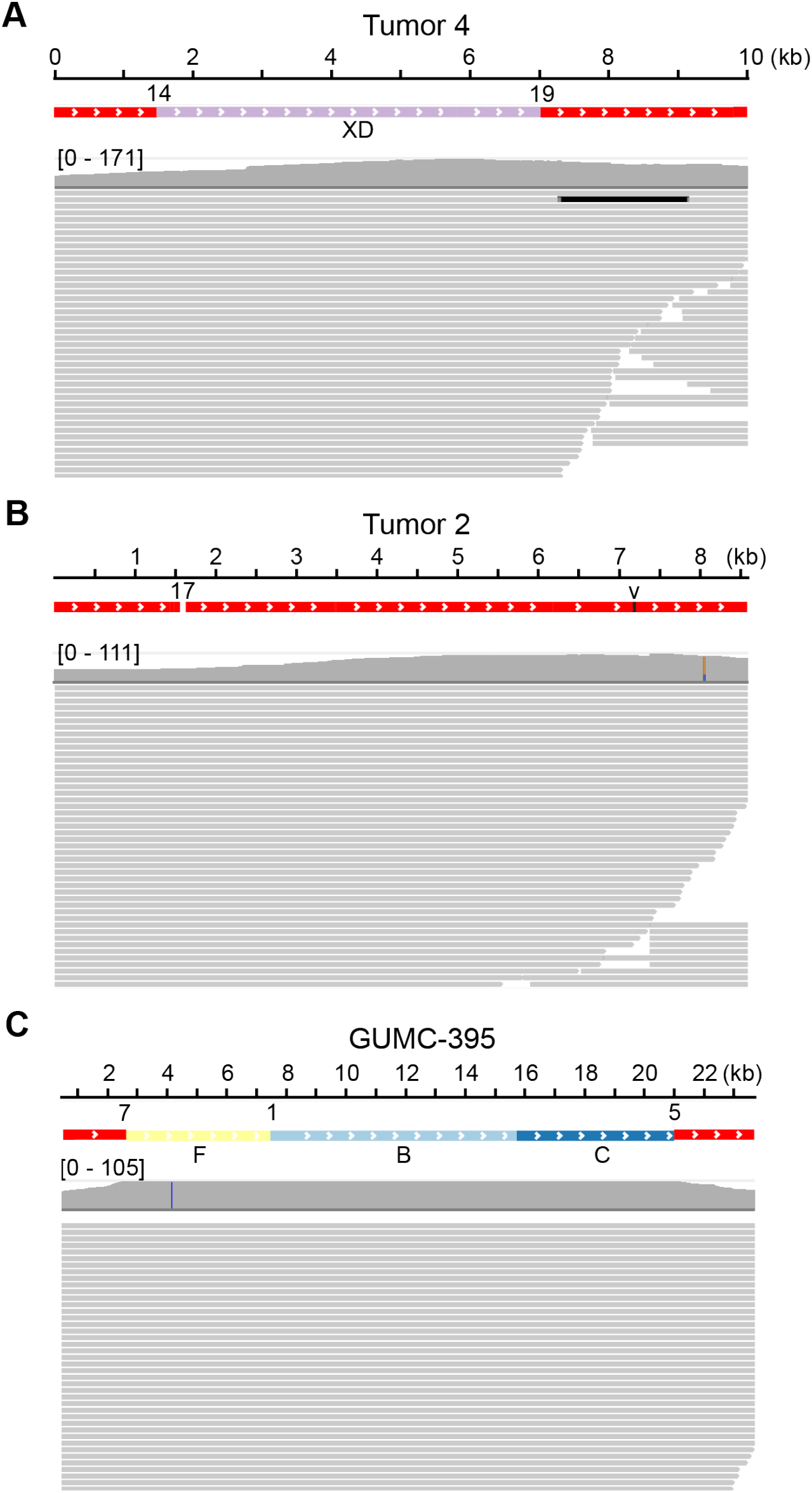
Absence of sequence variants at virus-host and virus-virus breakpoints. The absence of small indels and single nucleotide variants at virus-host and virus-virus breakpoints supports the hypothesis that rolling-circle or recombination-dependent replication is the most likely mechanism of concatemerization and amplification of virus and/or host sequences. PacBio HiFi reads were aligned to genomic sequences spanning (A) virus-host breakpoints 14 and 19 in Tumor 4 (see **Fig. 2**); (B) virus-virus breakpoint 17 in Tumor 2 (see **Fig.4A,B**); and (C) virus-host breakpoints 7 and 5 and host-host breakpoint 1 in GUMC-395 (see **Fig. 5A,B**). *Gray lines*, PacBio HiFi reads; *X-axis*, genome coordinates; *tiles*, HPV (red) or host (other colors) genome segments; *gray histogram*, depth of sequencing coverage; *numbers above tiles*, breakpoint IDs; v, normal virus-virus head-to-tail junction, *letters below tiles*, segment IDs.

**Extended Data Figure 3.**
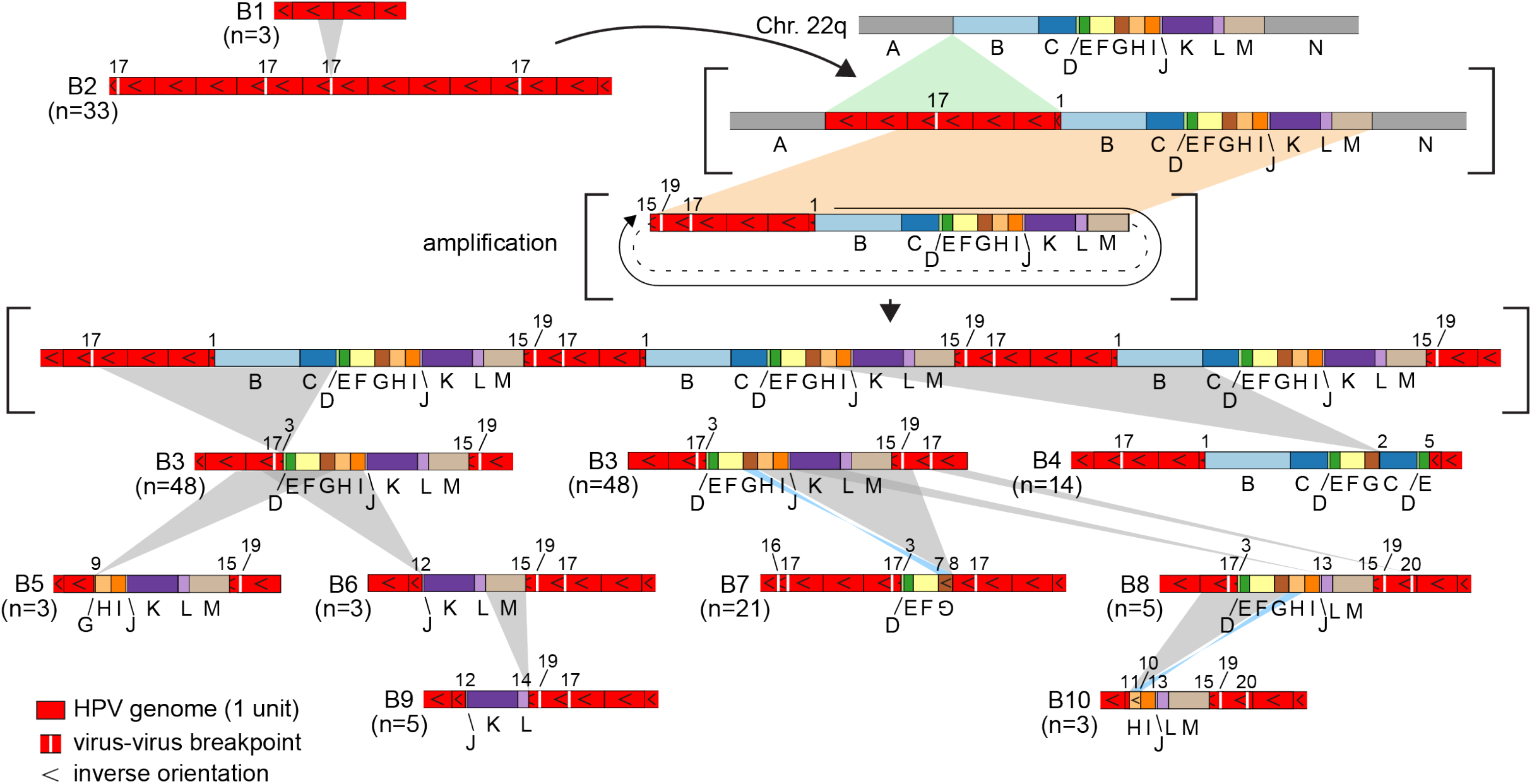
Evolution of structural variants at Chr. 22 of Tumor 2. Schematic depicts how various groups of structural variants evolved from a common molecular ancestor. *Block diagrams* (e.g., B1, B2, B3), representative ONT reads as in **Figure 4B**. *brackets*, hypothetical intermediate structures; *gray*, deletions; *green*, insertions; *tan*, ecDNA excisions; *blue*, inversions; *dashed lines*, circularized segments; *circular arrow*, amplification; *block colors*, segments defined in **Fig. 4A,B**.

**Extended Data Figure 4.**
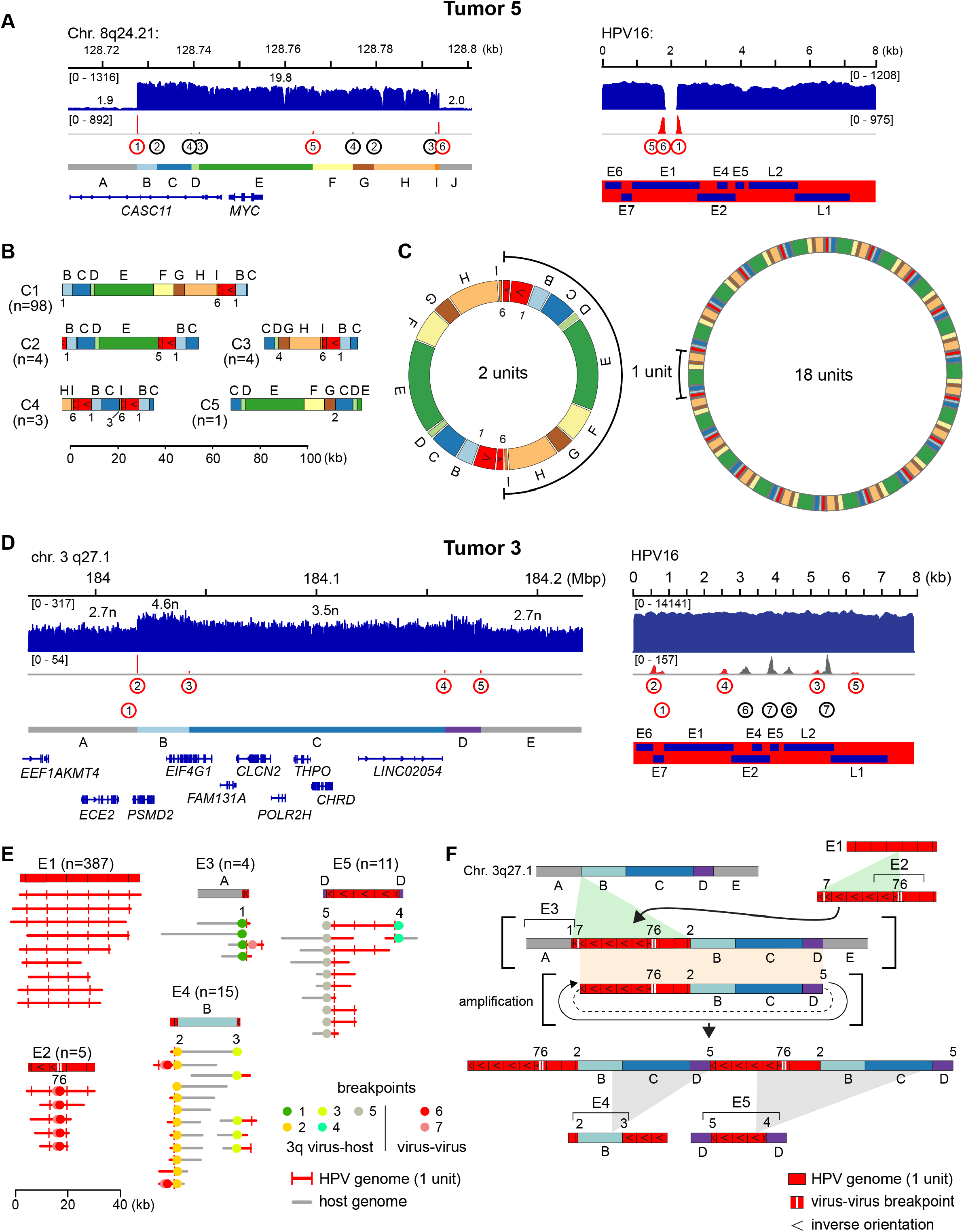
Evidence of heterocateny in Tumor 5 and Tumor 3. Analysis of LR-seq reads from Tumor 5 (A-C) and Tumor 3 (D-F) revealed additional evidence for heterocateny. (A) Depths of sequencing coverage and breakpoints at HPV integration sites at (*left*) the *MYC* locus at Chr. 8q and (*right*) in the HPV16 genome. *Top*, IGV browser display of (*y-axis, blue*) WGS coverage. *Middle*, (*red*) virus-host and (*gray*) host-host or virus-virus breakpoints; *circled numbers*, identifiers of segment-defining breakpoints (see **Supp Table 3.3**). *Bottom left*, genomic segments defined by breakpoints (see **Supp Tables 3.4**) and genes therein. *Bottom right*, HPV genes. (B) Tumor 5 ONT reads ≥20 kb were sorted into groups C1-C5 based on shared breakpoints. Block diagrams depict an ONT read representative of each group. Host segments (*letters, above*) and breakpoints (*numbers, below*) defined in panel A. n, number of ONT reads per group; *X-axis*, read length. (C) Tumor 5 WGS data indicated ∼20 HPV16 genome copies per cell and 9-fold amplification of *MYC*. Models show potential numbers of unit chains in ecDNAs, ranging from 2 (*left*) to 18 (*right*) linked units per molecule. Many combinations per concatemer are possible but cannot be distinguished using current methods. (D) Depths of sequencing coverage and breakpoints at HPV integration sites at (*left*) Chr. 3q and (*right*) in the HPV16 genome. *Top*, IGV browser display of (*y-axis, blue*) WGS coverage. *Middle* (*red*) virus-host and (*gray*) host-host or virus-virus breakpoints; *circled numbers*, identifiers of segment-defining (*upper row*) and nondefining (*lower row*) breakpoints (see **Supp Table 3.5**). *Bottom left*, genomic segments defined by breakpoints (see **Supp Table 3.6**) and genes therein. *Bottom right*, HPV genes. (E) Tumor 3 ONT reads ≥10 kb containing HPV sequences were sorted into groups E1-E5. Block diagram (*top*) depicts virus and host segments in an ONT read representative of each group and breakpoint plots (*below*) depict patterns of breakpoints and genomic segments in reads belonging to the group. Groups E1-E5 were defined by shared breakpoint patterns per breakpoint IDs (*numbers below block diagrams*). n, number of ONT reads per group; *X-axis*, read length. (F) Model depicts the evolution of structural variants shown in panel E from a common molecular ancestor. *Brackets*, hypothetical intermediate structures; *gray*, deletions; *green*, insertions; *tan*, ecDNA excisions; *dashed lines*, circularized segments; *circular arrow*, amplification; *block colors*, segments defined in panel D.

**Extended Data Figure 5.**
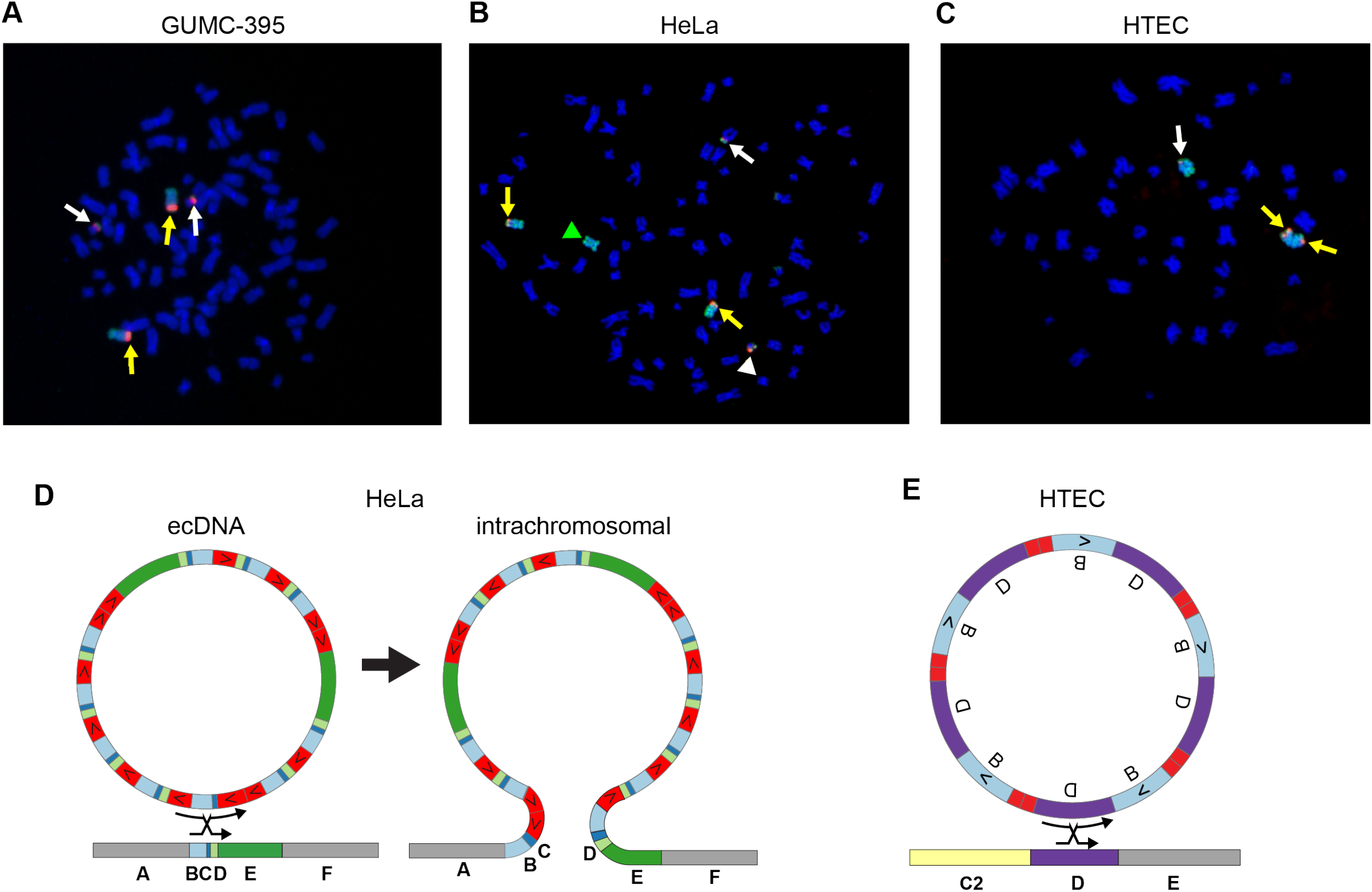
HPV integration into icDNA in GUMC-395, HeLa, and HTEC. To visualize HPV integrants in intrachromosomal DNA (icDNA), we performed fluorescence in situ hybridization on metaphase chromosomes of GUMC-395, HeLa, and HTEC cells using probes for HPV and for Chr. 8 (A-C). (A) In GUMC-395, HPV16 probe (red) labeled two copies of Chr. 8 (*yellow arrow*s) and two copies of t(8;21)(q24:q11) chromosomal translocation (*white arrows*). (B) In HeLa, HPV18 probe (red) labeled two copies of Chr. 8 (*yellow arrows*) but not a third copy (*green arrowhead*) and translocations on Chr. 22 (*white arrowhead*) and 5 (*white arrow*). (C) In HTEC, HPV16 probe labeled Chr. 8 (*white arrow*) and chromosome 8q iso-chromosome (*yellow arrows*). (D,E) Models showing the integration of virus-host concatemers in ecDNA form into Chr. 8 icDNA in (D) HeLa or (E) HTEC by homologous recombination. Virus-host concatemers and host gene segments as in **Fig. 6B** for HeLa and **Fig. 6F** for HTEC. *X*, site of homologous recombination between ecDNA and icDNA.

**Extended Data Figure 6.**
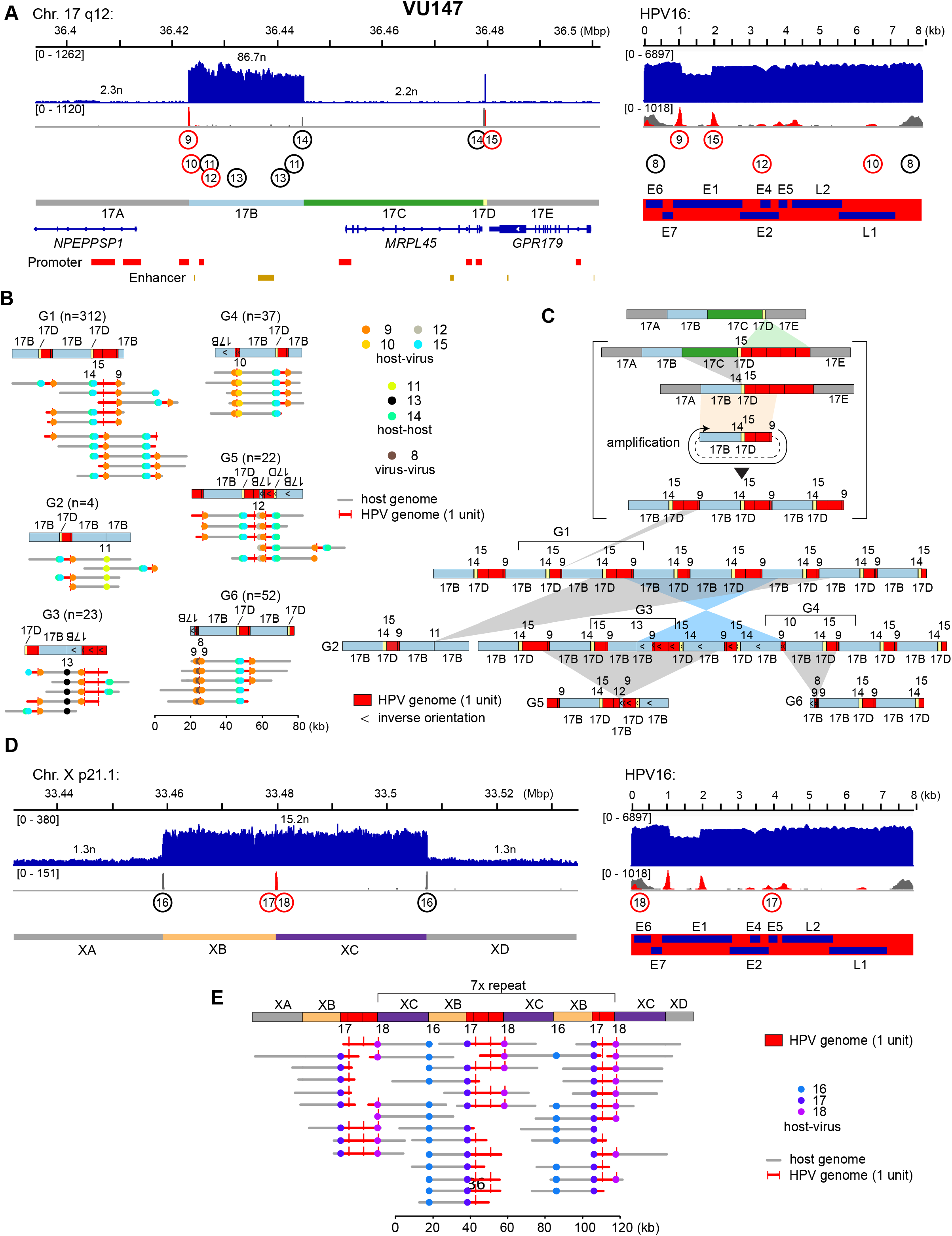
HPV integrants at Chr. 17 and Chr. X of VU147. Analysis of LR-seq reads from VU147 revealed HPV integrants at Chrs. 17 (A-C) and X (D,E). (A) Depths of sequencing coverage and breakpoints at HPV integration sites at (*left*) Chr. 17q and (*right*) in the HPV16 genome. *Top*, IGV browser display of (*y-axis, blue*) WGS coverage. *Middle*, (*red*) virus-host and (*gray*) host-host or virus-virus breakpoints; *circled numbers*, identifiers of segment-defining (*upper row*) and nondefining (*lower row*) breakpoints (see **Supp Table 5.3**). *Bottom left*, genomic segments defined by breakpoints (see **Supp Table 5.4**) and genes (*blue lines*), promoters (red), and enhancers (yellow) reported in the Ensembl database. *Bottom right*, HPV genes. (B) VU147 ONT reads were sorted into groups G1-G6. Block diagram (*top*) depicts virus and host segments in an ONT read representative of each group and breakpoint plots (*below*) depict patterns of breakpoints and genomic segments in reads belonging to the group. Groups G1-G6 were defined by shared breakpoint patterns per breakpoint IDs (*numbers below block diagrams*). n, number of ONT reads per group; *X-axis*, read length. (C) A model depicts how Chr. 17 structural variants evolved from a common molecular ancestor. *Block diagrams* (e.g., G1-G6), representative ONT reads as in panel B; *brackets*, hypothetical intermediate structures; *gray*, deletions; *green*, insertions; *tan*, ecDNA excisions; *blue*, inversions; *dashed lines*, circularized segments; *circular arrow*, amplification; block colors, segments defined in panel A. (D) Depths of sequencing coverage and breakpoints at HPV integration sites at (*left*) Chr. Xp and (*right*) in the HPV16 genome. *Top*, IGV browser display of (*y-axis, blue*) WGS coverage. *Middle*, (*red*) virus-host and (*gray*) host-host or virus-virus breakpoints; *circled numbers*, identifiers of segment-defining breakpoints (see **Supp Table 5.3**). *Bottom left*, genomic segments defined by breakpoints (see **Supp Table 5.4**). *Bottom right*, HPV genes. (E) *Top*, block diagrams depicting concatemerized HPV integrants and rearrangements integrated into flanking intrachromosomal segments at Chr. Xp21.1. *Bottom*, breakpoint plots depicting representative ONT reads of length >20 kb, as annotated in legend. Multiple ONT reads demonstrate intrachromosomal integration as they directly connect concatemers with flanking host DNA segments XA (*left*) and XD (*right*).

**Extended Data Figure 7.**
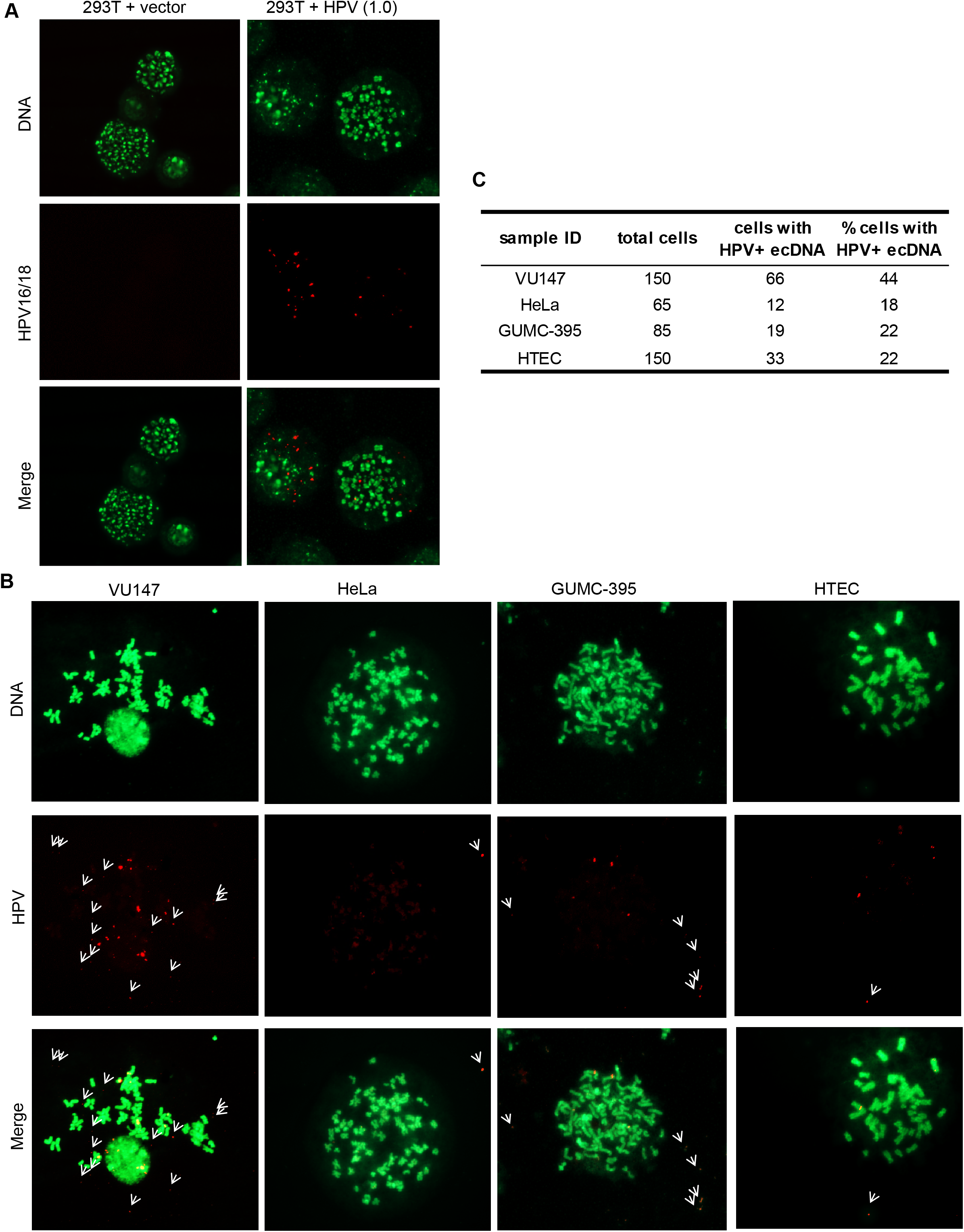
Metaphase FISH identifies HPV-containing ecDNAs in cell lines. To test for HPV integrants in ecDNA, we performed metaphase fluorescence *in situ* hybridization using probes for HPV16 or -18 on VU147, HeLa, GUMC-395, HTEC, and, as a positive control, 293T cells transfected with a plasmid containing the HPV16 genome. (A,B) *Red*, HPV16 (293T, VU147, GUMC-395, and HTEC) or HPV18 (HeLa) probe. *Green*, YOYO-1 DNA dye. (A) Micrographs (63x) of 293T cells transfected with vector control (vector) or HPV16 genome (HPC). (B) Micrographs of VU147, HeLa, GUMC-395, and HTEC cell lines. *White arrows*, ecDNA labeled with HPV probe. (C) Summary table indicating for each cell line (sample ID) the total number of cells evaluated (total cells), the number of cells with ecDNA labeled with HPV probe (cells with HPV+ ecDNA), and the percentage of cells with ecDNA labeled with HPV probe (% cells with HPV+ ecDNA).

**Extended Data Figure 8.**
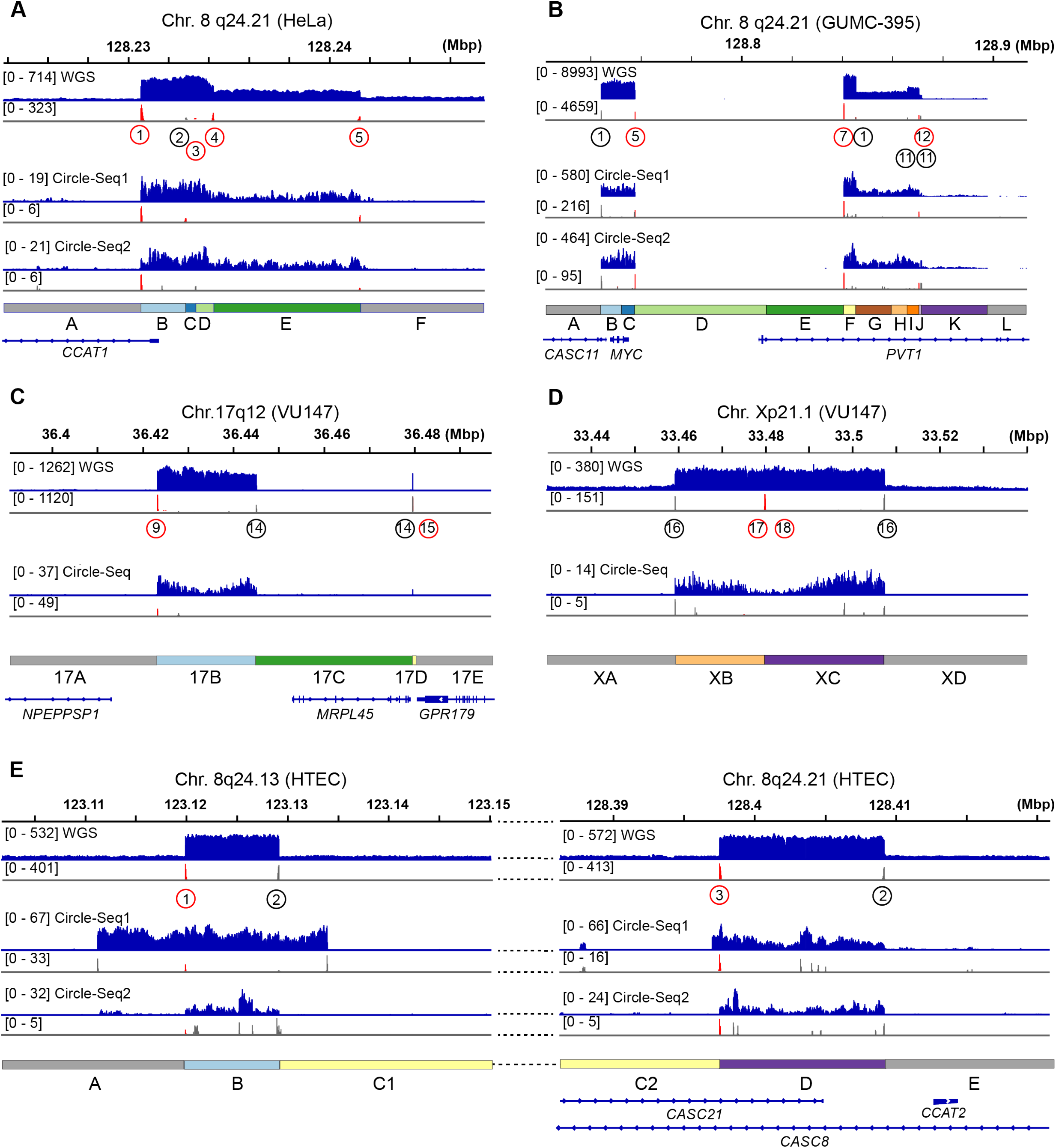
Circle-seq analysis of HeLa, GUMC-395, VU147, and HTEC cells. To address the possibility of co-existence of ecDNA in cell lines in which LR-Seq data and FISH indicated integration of virus-host concatemers in icDNA, we performed Circle-seq on HeLa, GUMC-395, VU147, and HTEC cells. (A-E) IGV browser views of (*top track*) WGS and (*middle, lower tracks*) Circle-seq replicate experimental data from (A) HeLa; (B) GUMC-395; (C,D) VU147; and (E) HTEC cells. Sequencing reads were mapped to HPV integration sites at (A) Chr. 8q24.21; (B) Chr. 8q24.21; (C) Chr. 17q12; (D) Xp21; and (E) *left*, Chr. 8q24.13; *right*, Chr. 8q24.21. *X-axis, top and bottom*, genomic coordinates, segments and gene schematic; *navy histogram*, depth of sequencing coverage (WGS and circle-seq replicates 1 and 2, respectively); *numbers in brackets*, counts of reads; *red spikes*, counts of HPV breakpoint-specific reads; *gray spikes*, counts of host-host breakpoint reads.

**Extended Data Figure 9.**
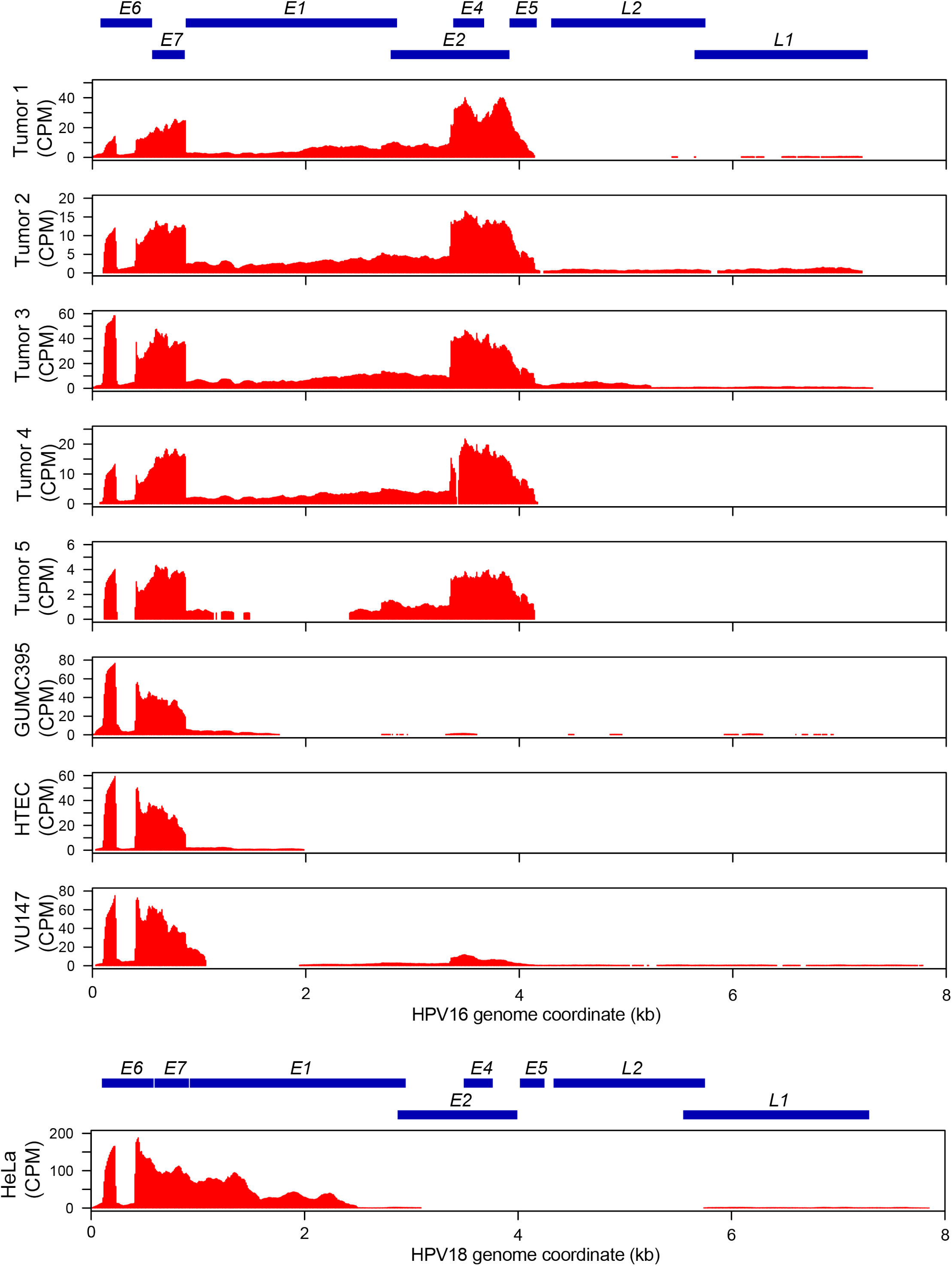
Transcription levels of HPV genes. Plots show expression levels of HPV16 or 18 transcripts using Illumina RNA-Seq data. *X-axis*, HPV genome coordinates. *Y-axis*, transcription level normalized to library size (CPM; counts per million reads). *Navy tiles*, coding regions of HPV genes.

## REFERENCES

1. Forman D, de Martel C, Lacey CJ, Soerjomataram I, Lortet-Tieulent J, Bruni L, et al. Global burden of human papillomavirus and related diseases. Vaccine. 20121;30 Suppl 5(2):F12–23.

2. Akagi K, Li J, Broutian TR, Padilla-Nash H, Xiao W, Jiang B, et al. Genome-wide analysis of HPV integration in human cancers reveals recurrent, focal genomic instability. Genome Res. 2014;24(2):185–99.

3. Cancer Genome Atlas Research N, Albert Einstein College of M, Analytical Biological S, Barretos Cancer H, Baylor College of M, Beckman Research Institute of City of H, et al. Integrated genomic and molecular characterization of cervical cancer. Nature. 2017;543(7645):378–84.

4. Parfenov M, Pedamallu CS, Gehlenborg N, Freeman SS, Danilova L, Bristow CA, et al. Characterization of HPV and host genome interactions in primary head and neck cancers. Proc Natl Acad Sci U S A. 2014;111(43):15544–9.

5. Symer DE, Akagi K, Geiger HM, Song Y, Li G, Emde AK, et al. Diverse tumorigenic consequences of human papillomavirus integration in primary oropharyngeal cancers. Genome Res. 2022;32(1):55–70.

6. Pang J, Nguyen N, Luebeck J, Ball L, Finegersh A, Ren S, et al. Extrachromosomal DNA in HPV-Mediated Oropharyngeal Cancer Drives Diverse Oncogene Transcription. Clin Cancer Res. 2021;27(24):6772–86.

7. Jeon S, Lambert PF. Integration of human papillomavirus type 16 DNA into the human genome leads to increased stability of E6 and E7 mRNAs: implications for cervical carcinogenesis. Proc Natl Acad Sci U S A. 1995;92(5):1654–8.

8. Crook T, Tidy JA, Vousden KH. Degradation of p53 can be targeted by HPV E6 sequences distinct from those required for p53 binding and trans-activation. Cell. 1991;67(3):547–56.

9. Gonzalez SL, Stremlau M, He X, Basile JR, Münger K. Degradation of the retinoblastoma tumor suppressor by the human papillomavirus type 16 E7 oncoprotein is important for functional inactivation and is separable from proteasomal degradation of E7. J Virol. 2001;75(16):7583–91.

10. Ojesina AI, Lichtenstein L, Freeman SS, Pedamallu CS, Imaz-Rosshandler I, Pugh TJ, et al. Landscape of genomic alterations in cervical carcinomas. Nature. 2014;506(7488):371–5.

11. Steenbergen RD, Hermsen MA, Walboomers JM, Joenje H, Arwert F, Meijer CJ, et al. Integrated human papillomavirus type 16 and loss of heterozygosity at 11q22 and 18q21 in an oral carcinoma and its derivative cell line. Cancer Res. 1995;55(22):5465–71.

12. Yuan H, Krawczyk E, Blancato J, Albanese C, Zhou D, Wang N, et al. HPV positive neuroendocrine cervical cancer cells are dependent on Myc but not E6/E7 viral oncogenes. Sci Rep. 2017;7:45617.

13. Schneider-Gadicke A, Schwarz E. Different human cervical carcinoma cell lines show similar transcription patterns of human papillomavirus type 18 early genes. EMBO J. 1986;5(9):2285–92.

14. Lace MJ, Anson JR, Klingelhutz AJ, Lee JH, Bossler AD, Haugen TH, et al. Human papillomavirus (HPV) type 18 induces extended growth in primary human cervical, tonsillar, or foreskin keratinocytes more effectively than other high-risk mucosal HPVs. J Virol. 2009;83(22):11784–94.

15. Zhou L, Qiu Q, Zhou Q, Li J, Yu M, Li K, et al. Long-read sequencing unveils high-resolution HPV integration and its oncogenic progression in cervical cancer. Nat Commun. 2022;13(1):2563.

16. Rossi NM, Dai J, Xie Y, Lou H, Boland JF, Yeager M, et al. Extrachromosomal Amplification of Human Papillomavirus Episomes as a Mechanism of Cervical Carcinogenesis. bioRxiv. 2021:2021.10.22.465367.

17. Dyer N, Young L, Ott S. Artifacts in the data of Hu et al. tNat Genet. 2016;48(1):2–4.

18. Sakakibara N, Chen D, McBride AA. Papillomaviruses use recombination-dependent replication to vegetatively amplify their genomes in differentiated cells. PLoS Pathog. 2013;9(7):e1003321.

19. McBride AA. Mechanisms and strategies of papillomavirus replication. Biol Chem. 2017;398(8):919–27.

20. Liblekas L, Piirsoo A, Laanemets A, Tombak EM, Laaneväli A, Ustav E, et al. Analysis of the Replication Mechanisms of the Human Papillomavirus Genomes. Front Microbiol. 2021;12:738125.

21. Chang HHY, Pannunzio NR, Adachi N, Lieber MR. Non-homologous DNA end joining and alternative pathways to double-strand break repair. Nat Rev Mol Cell Biol. 2017;18(8):495–506.

22. Deshpande V, Luebeck J, Nguyen ND, Bakhtiari M, Turner KM, Schwab R, et al. Exploring the landscape of focal amplifications in cancer using AmpliconArchitect. Nat Commun. 2019;10(1):392.

23. Gillison ML, Akagi K, Xiao W, Jiang B, Pickard RKL, Li J, et al. Human papillomavirus and the landscape of secondary genetic alterations in oral cancers. Genome Res. 2019;29(1):1–17.

24. Bodelon C, Untereiner ME, Machiela MJ, Vinokurova S, Wentzensen N. Genomic characterization of viral integration sites in HPV-related cancers. Int J Cancer. 2016;139(9):2001–11.

25. Adey A, Burton JN, Kitzman JO, Hiatt JB, Lewis AP, Martin BK, et al. The haplotype-resolved genome and epigenome of the aneuploid HeLa cancer cell line. Nature. 2013;500(7461):207–11.

26. Landry JJ, Pyl PT, Rausch T, Zichner T, Tekkedil MM, Stutz AM, et al. The genomic and transcriptomic landscape of a HeLa cell line. G3 (Bethesda). 2013;3(8):1213–24.

27. Macville M, Schrock E, Padilla-Nash H, Keck C, Ghadimi BM, Zimonjic D, et al. Comprehensive and definitive molecular cytogenetic characterization of HeLa cells by spectral karyotyping. Cancer Res. 1999;59(1):141–50.

28. Dürst M, Croce CM, Gissmann L, Schwarz E, Huebner K. Papillomavirus sequences integrate near cellular oncogenes in some cervical carcinomas. Proc Natl Acad Sci U S A. 1987;84(4):1070–4.

29. Kristiansen E, Jenkins A, Holm R. Coexistence of episomal and integrated HPV16 DNA in squamous cell carcinoma of the cervix. J Clin Pathol. 1994;47(3):253–6.

30. Nulton TJ, Olex AL, Dozmorov M, Morgan IM, Windle B. Analysis of The Cancer Genome Atlas sequencing data reveals novel properties of the human papillomavirus 16 genome in head and neck squamous cell carcinoma. Oncotarget. 2017;8(11):17684–99.

31. Greaves M, Maley CC. Clonal evolution in cancer. Nature. 2012;481(7381):306–13.

32. Koche RP, Rodriguez-Fos E, Helmsauer K, Burkert M, MacArthur IC, Maag J, et al. Publisher Correction: Extrachromosomal circular DNA drives oncogenic genome remodeling in neuroblastoma. Nat Genet. 2020;52(4):464.

33. deCarvalho AC, Kim H, Poisson LM, Winn ME, Mueller C, Cherba D, et al. Discordant inheritance of chromosomal and extrachromosomal DNA elements contributes to dynamic disease evolution in glioblastoma. Nat Genet. 2018;50(5):708–17.

34. Wu S, Turner KM, Nguyen N, Raviram R, Erb M, Santini J, et al. Circular ecDNA promotes accessible chromatin and high oncogene expression. Nature. 2019;575(7784):699–703.

35. Bailey C, Shoura MJ, Mischel PS, Swanton C. Extrachromosomal DNA-relieving heredity constraints, accelerating tumour evolution. Ann Oncol. 2020;31(7):884–93.

36. Nathanson DA, Gini B, Mottahedeh J, Visnyei K, Koga T, Gomez G, et al. Targeted therapy resistance mediated by dynamic regulation of extrachromosomal mutant EGFR DNA. Science. 2014;343(6166):72–6.

37. Verhaak RGW, Bafna V, Mischel PS. Extrachromosomal oncogene amplification in tumour pathogenesis and evolution. Nat Rev Cancer. 2019;19(5):283–8.

38. McBride AA. Replication and partitioning of papillomavirus genomes. Adv Virus Res. 2008;72:155–205.

39. Pittayakhajonwut D, Angeletti PC. Analysis of cis-elements that facilitate extrachromosomal persistence of human papillomavirus genomes. Virology. 2008;374(2):304–14.

40. Flores ER, Lambert PF. Evidence for a switch in the mode of human papillomavirus type 16 DNA replication during the viral life cycle. J Virol. 1997;71(10):7167–79.

41. Orav M, Geimanen J, Sepp EM, Henno L, Ustav E, Ustav M. Initial amplification of the HPV18 genome proceeds via two distinct replication mechanisms. Sci Rep. 2015;5:15952.

42. Hoffmann R, Hirt B, Bechtold V, Beard P, Raj K. Different modes of human papillomavirus DNA replication during maintenance. Virol. 2006;80(9):4431–9.

43. Moody CA, Laimins LA. Human papillomaviruses activate the ATM DNA damage pathway for viral genome amplification upon differentiation. PLoS Pathog. 2009;5(10):e1000605.

44. Henssen A, MacArthur I, Koche R, Dorado García H. Purification and Sequencing of Large Circular DNA from Human Cells. Protocol Exchange. 2019.

45. Layer RM, Chiang C, Quinlan AR, Hall IM. LUMPY: a probabilistic framework for structural variant discovery. Genome Biol. 2014;15(6):R84.

46. Li H. Minimap2: pairwise alignment for nucleotide sequences. Bioinformatics. 2018;34(18):3094–100.

47. Mahmoud M, Doddapaneni H, Timp W, Sedlazeck FJ. PRINCESS: comprehensive detection of haplotype resolved SNVs, SVs, and methylation. Genome Biol. 2021;22(1):268.

48. Ren J, Chaisson MJP. lra: A long read aligner for sequences and contigs. PLoS Comput Biol. 2021;17(6):e1009078.

49. Sedlazeck FJ, Rescheneder P, Smolka M, Fang H, Nattestad M, von Haeseler A, et al. Accurate detection of complex structural variations using single-molecule sequencing. Nat Methods. 2018;15(6):461–8.

50. Li H, Durbin R. Fast and accurate short read alignment with Burrows-Wheeler transform. Bioinformatics. 2009;25(14):1754–60.

51. Møller HD, Mohiyuddin M, Prada-Luengo I, Sailani MR, Halling JF, Plomgaard P, et al. Circular DNA elements of chromosomal origin are common in healthy human somatic tissue. Nat Commun. 2018;9(1):1069.

52. Dobin A, Davis CA, Schlesinger F, Drenkow J, Zaleski C, Jha S, et al. STAR: ultrafast universal RNA-seq aligner. Bioinformatics. 2013;29(1):15–21.

53. Schrock E, du Manoir S, Veldman T, Schoell B, Wienberg J, Ferguson-Smith MA, et al. Multicolor spectral karyotyping of human chromosomes. Science. 1996;273(5274):494–7.

