## Supplementary Tables for "Intratumoral heterogeneity and clonal evolution induced by HPV integration"

**Akagi, Symer, et al.**

### SUPPLEMENTARY TABLES

| ID | sample ID | WGS, Illumina | | | LR-seq, ONT | | | LR-seq, pacBio HiFi | | |
|---|---|---|---|---|---|---|---|---|---|---|
| | | autosome coverage (x) | MT coverage (x) | MT copy number | autosome coverage (x) | MT coverage (x) | MT copy number | autosome coverage (x) | MT coverage (x) | MT copy number |
| GS18097 | Tumor 1 | 95.9 | 28639.4 | 597.3 | 27.0 | 2226.0 | 164.9 | NA | NA | NA |
| GS18047 | Tumor 2 | 109.4 | 22013.6 | 402.4 | 24.5 | 2008.3 | 163.9 | 10.1 | 793.9 | 157.2 |
| GS18006 | Tumor 3 | 115.5 | 20196.3 | 349.7 | 30.8 | 5069.1 | 329.2 | 9.5 | 379.3 | 79.9 |
| GS18001 | Tumor 4 | 107.1 | 19234.7 | 359.2 | 19.4 | 823.0 | 84.8 | 9.2 | 236.5 | 51.4 |
| GS18074 | Tumor 5 | 98.4 | 38740.9 | 787.4 | 20.5 | 2211.8 | 215.8 | 9.4 | 445.7 | 94.8 |
| 93-VU-147T | VU147 | 50.9 | 22675.7 | 891.0 | 26.4 | 445.8 | 33.8 | 7.7 | 213.1 | 55.4 |
| HeLa | HeLa | 73.4 | 31174.7 | 849.4 | 31.3 | 1349.3 | 86.2 | 7.1 | 205.1 | 57.8 |
| GUMC-395 | GUMC-395 | 105.8 | 45036.6 | 851.4 | 25.5 | 508.3 | 39.9 | 15.8 | 410.2 | 51.9 |
| HTEC | HTEC | 52.2 | 36313.6 | 1391.3 | 15.0 | 912.9 | 121.7 | 9.1 | 859.0 | 188.8 |

**Supplementary Table 1.1. Depths of autosomal and mitochondrial genome sequencing coverage from WGS and LR-seq.** Five HPV-positive primary oropharyngeal cancers (Tumors 1-5) and four HPV-positive cell lines were evaluated by whole genome sequencing (WGS) on the Illumina platform and by long range sequencing (LR-seq) on the Oxford Nanopore Technologies (ONT) and PacBio HiFi platforms. Shown for each platform are depths of sequencing coverage for human autosomes and the mitochondrial (MT) genome and estimated mitochondria genome copy number per haploid genome, calculated by dividing MT depth of coverage by autosome depth of coverage x 2. We previously analyzed the five primary tumors (Tumor 1-5) by Illumina WGS, as described in Symer et al. *Genome Research* 2022 (PMID: 34903527).

| sample ID | WGS, Illumina | | | LR-seq, ONT | | | LR-seq, PacBio HiFi | | |
|---|---|---|---|---|---|---|---|---|---|
| | autosome coverage (x) | HPV coverage (x) | HPV copy number | autosome coverage (x) | HPV coverage (x) | HPV copy number | autosome coverage (x) | HPV coverage (x) | HPV copy number |
| Tumor 1 | 95.9 | 7512.6 | 156.7 | 27.0 | 408.8 | 30.3 | NA | NA | NA |
| Tumor 2 | 109.4 | 9106.9 | 166.5 | 24.5 | 1724.5 | 140.8 | 10.1 | 780.4 | 154.5 |
| Tumor 3 | 115.5 | 13421.8 | 232.4 | 30.8 | 3706.2 | 240.7 | 9.5 | 726.5 | 152.9 |
| Tumor 4 | 107.1 | 9191.5 | 171.6 | 19.4 | 1029.4 | 106.1 | 9.2 | 410.5 | 89.2 |
| Tumor 5 | 98.4 | 1062.6 | 21.6 | 20.5 | 201.1 | 19.6 | 9.4 | 103.4 | 22.0 |
| VU147 | 50.9 | 6314.3 | 248.1 | 26.4 | 3103.9 | 235.1 | 7.7 | 820.9 | 213.2 |
| HeLa | 73.4 | 235.7 | 6.4 | 31.3 | 110.9 | 7.1 | 7.1 | 12.4 | 3.5 |
| GUMC-395 | 105.8 | 5907.9 | 111.7 | 25.5 | 1378.9 | 108.1 | 15.8 | 784.8 | 99.3 |
| HTEC | 52.2 | 225.5 | 8.6 | 15.0 | 66.3 | 8.8 | 9.1 | 55.8 | 12.3 |

**Supplementary Table 1.2. Depths of autosomal and HPV genome sequencing coverage from WGS and LR-seq.** Five HPV-positive primary oropharyngeal cancers (Tumors 1-5) and four HPV-positive cell lines were evaluated by whole genome sequencing (WGS) on the Illumina platform and by long range sequencing (LR-seq) on the Oxford Nanopore Technologies (ONT) and PacBio HiFi platforms. Shown for each platform are depths of sequencing coverage for human autosomes and the HPV genome and estimated HPV genome copy number, calculated by dividing HPV depth of coverage by autosome depth of coverage x 2. We previously analyzed the five primary tumors (Tumor 1-5) by Illumina WGS, as described in Symer et al. *Genome Research* 2022 (PMID: 34903527).

| ID | chrom1 | start1 | stop1 | chrom2 | start2 | stop2 | description | normal | type | strand1 | strand2 | # reads ILMN | # reads PacBio | # reads ONT | segment defining |
|---|---|---|---|---|---|---|---|---|---|---|---|---|---|---|---|
| 1 | chr5 | 29653971 | 29653974 | chr5 | 29694739 | 29694759 | 5B-5A | N | DUP | - | + | 26 | 4 | 5 | N |
| 2 | chr5 | 29687607 | 29687627 | HPV16 | 4523 | 4525 | 5A-V | N | BND | + | - | 35 | 0 | 11 | N |
| 3 | chr5 | 29687635 | 29687636 | HPV16 | 3003 | 3004 | V-5B | N | BND | - | + | 231 | 36 | 50 | Y |
| 4 | chr5 | 29691736 | 29691756 | HPV16 | 431 | 434 | V-5B | N | BND | - | + | 28 | 0 | 6 | N |
| 5 | chr5 | 29698118 | 29698131 | chrX | 2211159 | 2211178 | XD-5B | N | BND | - | + | 31 | 10 | 17 | N |
| 6 | chr5 | 29700377 | 29700378 | chrX | 2208484 | 2208485 | 5B-XD | N | BND | + | - | 240 | 47 | 79 | Y |
| 7 | chr5 | 91746855 | 91746856 | HPV16 | 6100 | 6101 | V-5D | N | BND | - | + | 506 | 56 | 31 | Y |
| 8 | chr5 | 91746906 | 91746910 | chrX | 2207470 | 2207490 | 5D-XD | N | BND | + | - | 24 | 4 | 5 | N |
| 9 | chr5 | 91748270 | 91748271 | HPV16 | 7466 | 7467 | 5E-V | N | BND | + | - | 500 | 67 | 90 | Y |
| 10 | chr5 | 91784329 | 91784349 | HPV16 | 5897 | 5917 | V-5G | N | BND | - | + | 23 | 6 | 11 | Y |
| 11 | chr5 | 91795911 | 91795931 | HPV16 | 4901 | 4906 | 5G-V | N | BND | + | - | 21 | 2 | 9 | Y |
| 12 | chrX | 2161219 | 2161220 | chrX | 2208698 | 2208699 | XD-XB | N | DUP | - | + | 149 | 38 | 28 | Y |
| 13 | chrX | 2194280 | 2194281 | HPV16 | 5711 | 5712 | XB-V | N | BND | + | - | 151 | 26 | 28 | Y |
| 14 | chrX | 2205635 | 2205636 | HPV16 | 6940 | 6941 | V-XD | N | BND | - | + | 3123 | 185 | 386 | Y |
| 15 | chrX | 2205833 | 2205836 | HPV16 | 7849 | 7852 | V-XD | N | BND | - | + | 32 | 5 | 7 | N |
| 16 | chrX | 2206003 | 2206004 | HPV16 | 5341 | 5342 | V-XD | N | BND | - | + | 68 | 18 | 20 | N |
| 17 | chrX | 2206937 | 2206938 | HPV16 | 605 | 606 | V-XD | N | BND | - | + | 303 | 43 | 18 | N |
| 18 | chrX | 2210490 | 2210491 | HPV16 | 6539 | 6559 | XD-(V) | N | BND | - | - | 52 | 0 | 6 | N |
| 19 | chrX | 2211168 | 2211169 | HPV16 | 4329 | 4330 | XD-V | N | BND | + | - | 3998 | 448 | 511 | Y |
| 20 | HPV16 | 775 | 776 | HPV16 | 5144 | 5145 | V-V | N | DUP | - | + | 150 | 30 | 45 | N |
| 21 | HPV16 | 2174 | 2175 | HPV16 | 2709 | 2710 | V-V | N | DEL | + | - | 105 | 0 | 14 | N |
| 22 | HPV16 | 4621 | 4641 | HPV16 | 6437 | 6457 | V-V | N | DEL | + | - | 23 | 14 | 4 | N |
| a | chr5 | 29687633 | 29687634 | chr5 | 29687635 | 29687636 | 5A-5B | Y | LANDMARK | + | - | 63 | 10 | 22 | NA |
| b | chr5 | 29700377 | 29700378 | chr5 | 29700379 | 29700380 | 5B-5C | Y | LANDMARK | + | - | 40 | 24 | 19 | NA |
| d | chr5 | 91746854 | 91746855 | chr5 | 91746856 | 91746857 | 5D-5E | Y | LANDMARK | + | - | 47 | 82 | 44 | NA |
| e | chr5 | 91748268 | 91748269 | chr5 | 91748270 | 91748271 | 5E-5F | Y | LANDMARK | + | - | 72 | 28 | 21 | NA |
| f | chr5 | 91784328 | 91784329 | chr5 | 91784330 | 91784331 | 5F-5G | Y | LANDMARK | + | - | 76 | 6 | 14 | NA |
| g | chr5 | 91795909 | 91795910 | chr5 | 91795911 | 91795912 | 5G-5H | Y | LANDMARK | + | - | 55 | 14 | 15 | NA |
| xa | chrX | 2161219 | 2161220 | chrX | 2161221 | 2161222 | XA-XB | Y | LANDMARK | + | - | 44 | 14 | 12 | NA |
| xb | chrX | 2194278 | 2194279 | chrX | 2194280 | 2194281 | XB-XC | Y | LANDMARK | + | - | 26 | 13 | 8 | NA |

| | | | | | | | | | | | | | | | |
|---|---|---|---|---|---|---|---|---|---|---|---|---|---|---|---|
| xc | chrX | 2205633 | 2205634 | chrX | 2205635 | 2205636 | XC-XD | Y | LANDMARK | + | - | 17 | 10 | 16 | NA |
| xd | chrX | 2211168 | 2211169 | chrX | 2211170 | 2211171 | XD-XE | Y | LANDMARK | + | - | 47 | 11 | 22 | NA |
| v | HPV16 | 0 | 1 | HPV16 | 7905 | 7906 | V-V | Y | LANDMARK | - | + | 5921 | 682 | 818 | NA |

**Supplementary Table 2.1. Breakpoints and normal segment junctions in Tumor 4.** Genomic coordinates and other features of virus-virus, virus-host, and host-host breakpoints and of normal segment junctions detected in Tumor 4 upon analysis of WGS and LR-seq data. Description, defined segments connected at junction (see **Supplementary Table 2.2**); normal, presence (Y) or absence (N) of junction in reference genome sequence; type, junction classification by the Sniffles algorithm as duplication (DUP), deletion (DEL), inversion (INV), structural variation breakpoint (BND), or normal reference genome feature (LANDMARK); strand 1 and strand 2, + or - strand as per BEDPE format for WGS data; # reads, number of reads from indicated platform supporting junction; segment-defining, (Y) breakpoint used to define the termini of genomic segments typically >1 kb in length or (N) breakpoint not used to define genomic segments.

| chrom | start | stop | ID | size |
|---|---|---|---|---|
| chr5 | 28637633 | 29637633 | pad1 | 1000000 |
| chr5 | 29637634 | 29687635 | 5A | 50001 |
| chr5 | 29687636 | 29700378 | 5B | 12742 |
| chr5 | 29700379 | 29750379 | 5C | 50000 |
| chr5 | 29750380 | 30750380 | pad2 | 1000000 |
| chr5 | 90696909 | 91696909 | pad3 | 1000000 |
| chr5 | 91696910 | 91746855 | 5D | 49945 |
| chr5 | 91746856 | 91748269 | 5E | 1413 |
| chr5 | 91748270 | 91784329 | 5F | 36059 |
| chr5 | 91784330 | 91795910 | 5G | 11580 |
| chr5 | 91795911 | 91845911 | 5H | 50000 |
| chr5 | 91845912 | 92845912 | pad4 | 1000000 |
| chrX | 1111219 | 2111219 | pad5 | 1000000 |
| chrX | 2111220 | 2161220 | XA | 50000 |
| chrX | 2161221 | 2194279 | XB | 33058 |
| chrX | 2194280 | 2205634 | XC | 11354 |
| chrX | 2205635 | 2211169 | XD | 5534 |
| chrX | 2211170 | 2261170 | XE | 50000 |
| chrX | 2261171 | 3261171 | pad6 | 1000000 |
| HPV16 | 0 | 7906 | V | 7906 |

**Supplementary Table 2.2. Definition of genomic segments flanking HPV integrants in Tumor 4.** Shown here are chromosome coordinates for segments used for the analysis of Tumor 4 LR-seq and WGS data. Chromosome, segment start and stop positions, segment ID, and segment size are indicated. Coordinates are based on GRCh37 human reference genome sequence and HPV16 reference genome (NC_001526.3).

| ID | chrom1 | start1 | stop1 | chrom2 | start2 | stop2 | description | normal | type | strand1 | strand2 | #ILMN support | #pacBio support | #ONT support | segment defining |
|---|---|---|---|---|---|---|---|---|---|---|---|---|---|---|---|
| 1 | chr22 | 41512975 | 41512978 | HPV16 | 6766 | 6771 | (V)-B | N | BND | - | - | 27 | 2 | 6 | Y |
| 2 | chr22 | 41529727 | 41529728 | chr22 | 41547182 | 41547183 | G-C | N | DUP | - | + | 155 | 36 | 17 | Y |
| 3 | chr22 | 41537063 | 41537064 | HPV16 | 5395 | 5396 | (V)-D | N | BND | - | - | 487 | 58 | 91 | Y |
| 4 | chr22 | 41537402 | 41537422 | HPV16 | 3827 | 3830 | (V)-E | N | BND | - | - | 48 | 10 | 9 | Y |
| 5 | chr22 | 41539467 | 41539485 | HPV16 | 2302 | 2322 | E-(V) | N | BND | + | + | 69 | 0 | 22 | Y |
| 6 | chr22 | 41541002 | 41541022 | HPV16 | 5114 | 5134 | (V)-F | N | BND | - | - | 16 | 7 | 5 | N |
| 7 | chr22 | 41544346 | 41544347 | chr22 | 41547198 | 41547199 | F-(G) | N | BND | + | + | 143 | 18 | 32 | Y |
| 8 | chr22 | 41544375 | 41544376 | HPV16 | 6823 | 6824 | V-G | N | BND | - | + | 129 | 20 | 30 | N |
| 9 | chr22 | 41547096 | 41547116 | HPV16 | 2063 | 2082 | (V)-G | N | BND | - | - | 12 | 8 | 5 | N |
| 10 | chr22 | 41548059 | 41548069 | chr22 | 41550669 | 41550680 | (I)-H | N | BND | - | - | 12 | 3 | 5 | N |
| 11 | chr22 | 41550251 | 41550271 | HPV16 | 7024 | 7029 | H-V | N | BND | + | - | 24 | 6 | 5 | Y |
| 12 | chr22 | 41553267 | 41553270 | HPV16 | 5261 | 5264 | (V)-J | N | BND | - | - | 49 | 8 | 10 | Y |
| 13 | chr22 | 41553458 | 41553475 | chr22 | 41563386 | 41563389 | I-J | N | DEL | + | - | 45 | 16 | 16 | Y |
| 14 | chr22 | 41565565 | 41565573 | HPV16 | 6200 | 6208 | L-(V) | N | BND | + | + | 9 | 2 | 5 | Y |
| 15 | chr22 | 41573527 | 41573528 | HPV16 | 6196 | 6197 | M-(V) | N | BND | + | + | 139 | 82 | 47 | Y |
| 16 | HPV16 | 425 | 426 | HPV16 | 4014 | 4015 | V-V | N | DUP | - | + | 132 | 11 | 30 | N |
| 17 | HPV16 | 2312 | 2313 | HPV16 | 7065 | 7066 | V-V | N | DUP | - | + | 2401 | 420 | 434 | N |
| 18 | HPV16 | 1858 | 1859 | HPV16 | 7065 | 7066 | V-V | N | DEL | + | - | 166 | 62 | 35 | N |
| 19 | HPV16 | 2164 | 2165 | HPV16 | 6137 | 6138 | V-V | N | DEL | + | - | 167 | 42 | 53 | N |
| 20 | HPV16 | 917 | 918 | HPV16 | 2474 | 2490 | V-V | N | DEL | + | - | 69 | 9 | 12 | N |
| 21 | chr4 | 19108228 | 19108234 | HPV16 | 6852 | 6858 | 4A-(V) | N | BND | + | + | 20 | 6 | 4 | Y |
| 22 | chr4 | 19111938 | 19111958 | HPV16 | 1023 | 1026 | (V)-4C | N | BND | - | - | 45 | 12 | 12 | Y |
| 23 | chr4 | 19130806 | 19130826 | HPV16 | 7353 | 7359 | 4C-(V) | N | BND | + | + | 22 | 2 | 5 | Y |
| a | chr22 | 41512973 | 41512974 | chr22 | 41512975 | 41512976 | A-B | Y | LANDMARK | + | - | 110 | 30 | 23 | NA |
| b | chr22 | 41529727 | 41529728 | chr22 | 41529729 | 41529730 | B-C | Y | LANDMARK | + | - | 157 | 34 | 33 | NA |
| c | chr22 | 41537062 | 41537063 | chr22 | 41537064 | 41537065 | C-D | Y | LANDMARK | + | - | 324 | 52 | 45 | NA |
| d | chr22 | 41537401 | 41537402 | chr22 | 41537403 | 41537404 | D-E | Y | LANDMARK | + | - | 455 | 110 | 137 | NA |
| e | chr22 | 41539484 | 41539485 | chr22 | 41539486 | 41539487 | E-F | Y | LANDMARK | + | - | 617 | 61 | 115 | NA |
| f | chr22 | 41544346 | 41544347 | chr22 | 41544348 | 41544349 | F-G | Y | LANDMARK | + | - | 502 | 86 | 95 | NA |
| g | chr22 | 41547182 | 41547183 | chr22 | 41547184 | 41547185 | G-H | Y | LANDMARK | + | - | 342 | 59 | 78 | NA |

| | | | | | | | | | | | | | | | |
|---|---|---|---|---|---|---|---|---|---|---|---|---|---|---|---|
| h | chr22 | 41550270 | 41550271 | chr22 | 41550272 | 41550273 | H-I | Y | LANDMARK | + | - | 385 | 70 | 90 | NA |
| i | chr22 | 41553266 | 41553267 | chr22 | 41553268 | 41553269 | I-J | Y | LANDMARK | + | - | 409 | 74 | 91 | NA |
| j | chr22 | 41553474 | 41553475 | chr22 | 41553476 | 41553477 | J-K | Y | LANDMARK | + | - | 244 | 41 | 86 | NA |
| k | chr22 | 41563385 | 41563386 | chr22 | 41563387 | 41563388 | K-L | Y | LANDMARK | + | - | 290 | 82 | 93 | NA |
| l | chr22 | 41573527 | 41573528 | chr22 | 41573529 | 41573530 | L-M | Y | LANDMARK | + | - | 104 | 14 | 24 | NA |
| 4a | chr4 | 19108228 | 19108229 | chr4 | 19108239 | 19108240 | 4A-4B | Y | LANDMARK | + | - | 68 | 16 | 23 | NA |
| 4b | chr4 | 19111939 | 19111940 | chr4 | 19111949 | 19111950 | 4B-4C | Y | LANDMARK | + | - | 82 | 12 | 24 | NA |
| 4c | chr4 | 19130827 | 19130828 | chr4 | 19130838 | 19130839 | 4C-4D | Y | LANDMARK | + | - | 99 | 14 | 27 | NA |
| v | HPV16 | 0 | 1 | HPV16 | 7905 | 7906 | V-V | Y | LANDMARK | - | + | 7940 | 1255 | 1264 | NA |

**Supplementary Table 3.1. Breakpoints and normal segment junctions in Tumor 2.** Genomic coordinates and other features of virus-virus, virus-host, and host-host breakpoints and of normal segment junctions detected in Tumor 2 upon analysis of WGS and LR-seq data. Description, defined segments connected at junction (see **Supplementary Table 3.2**); normal, presence (Y) or absence (N) of junction in reference genome sequence; type, junction classification by the Sniffles algorithm as duplication (DUP), deletion (DEL), inversion (INV), structural variation breakpoint (BND), or normal reference genome feature (LANDMARK); strand 1 and strand 2, + or - strand as per BEDPE format for WGS data; # reads, number of reads from indicated platform supporting junction; segment-defining, (Y) breakpoint used to define the termini of genomic segments typically >1 kb in length or (N) breakpoint not used to define genomic segments.

| chrom | start | stop | ID | size |
|---|---|---|---|---|
| chr22 | 40462973 | 41462973 | pad22_1 | 1000000 |
| chr22 | 41462974 | 41512974 | 22A | 50000 |
| chr22 | 41512975 | 41529727 | 22B | 16752 |
| chr22 | 41529728 | 41537063 | 22C | 7335 |
| chr22 | 41537064 | 41537402 | 22D | 338 |
| chr22 | 41537403 | 41539485 | 22E | 2082 |
| chr22 | 41539486 | 41544347 | 22F | 4861 |
| chr22 | 41544348 | 41547183 | 22G | 2835 |
| chr22 | 41547184 | 41550271 | 22H | 3087 |
| chr22 | 41550272 | 41553267 | 22I | 2995 |
| chr22 | 41553268 | 41553475 | 22J | 207 |
| chr22 | 41553476 | 41563386 | 22K | 9910 |
| chr22 | 41563387 | 41565573 | 22L | 2186 |
| chr22 | 41565574 | 41573528 | 22M | 7954 |
| chr22 | 41573529 | 41623529 | 22N | 50000 |
| chr22 | 41623530 | 42623530 | pad22_2 | 1000000 |
| chr4 | 18058226 | 19058226 | pad4_1 | 1000000 |
| chr4 | 19058227 | 19108227 | 4A | 50000 |
| chr4 | 19108228 | 19111938 | 4B | 3710 |
| chr4 | 19111939 | 19130826 | 4C | 18887 |
| chr4 | 19130827 | 19180827 | 4D | 50000 |
| chr4 | 19180828 | 20180828 | pad4_2 | 1000000 |
| HPV16 | 0 | 7906 | V | 7906 |

**Supplementary Table 3.2. Definition of genomic segments flanking HPV integrants in Tumor 2.** Shown here are chromosome coordinates for segments used for the analysis of Tumor 2 LR-seq and WGS data. Chromosome, segment start and stop positions, segment ID, and segment size are indicated. Coordinates are based on GRCh37 human reference genome sequence and HPV16 reference genome (NC_001526.3).

| ID | chrom1 | start1 | stop1 | chrom2 | start2 | stop2 | description | normal | type | strand1 | strand2 | # reads ILMN | # reads PacBio | # reads ONT | segment defining |
|---|---|---|---|---|---|---|---|---|---|---|---|---|---|---|---|
| 1 | chr8 | 128727579 | 128727580 | HPV16 | 2170 | 2171 | (V)-B | N | BND | - | - | 786 | 97 | 189 | Y |
| 2 | chr8 | 128731980 | 128731995 | chr8 | 128779516 | 128779536 | G-C | N | DUP | - | + | 24 | 2 | 5 | Y |
| 3 | chr8 | 128739520 | 128739521 | chr8 | 128792937 | 128792938 | C-I | N | DEL | + | - | 79 | 19 | 16 | Y |
| 4 | chr8 | 128741221 | 128741222 | chr8 | 128774888 | 128774889 | D-G | N | DEL | + | - | 107 | 8 | 21 | Y |
| 5 | chr8 | 128766130 | 128766131 | HPV16 | 1760 | 1761 | E-(V) | N | BND | + | + | 156 | 14 | 19 | Y |
| 6 | chr8 | 128793667 | 128793668 | HPV16 | 1803 | 1804 | I-(V) | N | BND | + | + | 692 | 105 | 173 | Y |
| a | chr8 | 128727577 | 128727578 | chr8 | 128727579 | 128727580 | A-B | Y | LANDMARK | + | - | 96 | 8 | 21 | NA |
| b | chr8 | 128731979 | 128731980 | chr8 | 128731981 | 128731982 | B-C | Y | LANDMARK | + | - | 978 | 49 | 167 | NA |
| c | chr8 | 128739519 | 128739520 | chr8 | 128739521 | 128739522 | C-D | Y | LANDMARK | + | - | 1070 | 95 | 179 | NA |
| d | chr8 | 128741220 | 128741221 | chr8 | 128741222 | 128741223 | D-E | Y | LANDMARK | + | - | 973 | 90 | 168 | NA |
| e | chr8 | 128766129 | 128766130 | chr8 | 128766131 | 128766132 | E-F | Y | LANDMARK | + | - | 864 | 96 | 211 | NA |
| f | chr8 | 128774887 | 128774888 | chr8 | 128774889 | 128774890 | F-G | Y | LANDMARK | + | - | 916 | 76 | 192 | NA |
| g | chr8 | 128779515 | 128779516 | chr8 | 128779517 | 128779518 | G-H | Y | LANDMARK | + | - | 714 | 36 | 112 | NA |
| h | chr8 | 128792936 | 128792937 | chr8 | 128792938 | 128792939 | H-I | Y | LANDMARK | + | - | 514 | 90 | 181 | NA |
| i | chr8 | 128793666 | 128793667 | chr8 | 128793668 | 128793669 | I-J | Y | LANDMARK | + | - | 68 | 4 | 15 | NA |
| v | HPV16 | 0 | 1 | HPV16 | 7904 | 7905 | V-V | Y | LANDMARK | - | + | 1124 | 103 | 191 | NA |

**Supplementary Table 3.3. Breakpoints and normal segment junctions in Tumor 5.** Genomic coordinates and other features of virus-virus, virus-host, and host-host breakpoints and of normal segment junctions detected in Tumor 5 upon analysis of WGS and LR-seq data. Description, defined segments connected at junction (see **Supplementary Table 3.4**); normal, presence (Y) or absence (N) of junction in reference genome sequence; type, junction classification by the Sniffles algorithm as duplication (DUP), deletion (DEL), inversion (INV), structural variation breakpoint (BND), or normal reference genome feature (LANDMARK); strand 1 and strand 2, + or - strand as per BEDPE format for WGS data; # reads, number of reads from indicated platform supporting junction; segment-defining, (Y) breakpoint used to define the termini of genomic segments typically >1 kb in length or (N) breakpoint not used to define genomic segments.

| chrom | start | stop | ID | size |
|---|---|---|---|---|
| chr8 | 127677577 | 128677577 | pad1 | 1000000 |
| chr8 | 128677578 | 128727578 | A | 50000 |
| chr8 | 128727579 | 128731980 | B | 4401 |
| chr8 | 128731981 | 128739520 | C | 7539 |
| chr8 | 128739521 | 128741221 | D | 1700 |
| chr8 | 128741222 | 128766130 | E | 24908 |
| chr8 | 128766131 | 128774888 | F | 8757 |
| chr8 | 128774889 | 128779516 | G | 4627 |
| chr8 | 128779517 | 128792937 | H | 13420 |
| chr8 | 128792938 | 128793667 | I | 729 |
| chr8 | 128793668 | 128843668 | J | 50000 |
| chr8 | 128843669 | 129843669 | pad2 | 1000000 |
| HPV16 | 0 | 7906 | V | 7906 |

**Supplementary Table 3.4. Definition of genomic segments flanking HPV integrants in Tumor 5.** Shown here are chromosome coordinates for segments used for the analysis of Tumor 5 LR-seq and WGS data. Chromosome, start and stop positions, segment ID, and segment size are indicated. Coordinates are based on GRCh37 human reference genome sequence and HPV16 reference genome (NC_001526.3).

| ID | chrom1 | start1 | stop1 | chrom2 | start2 | stop2 | description | type | normal | strand1 | strand2 | # reads ILMN | # reads PacBio | # reads ONT | segment defining |
|---|---|---|---|---|---|---|---|---|---|---|---|---|---|---|---|
| 1 | chr3 | 184019096 | 184019100 | HPV16 | 824 | 834 | A-(V) | BND | N | + | + | 12 | 3 | 5 | N |
| 2 | chr3 | 184019114 | 184019115 | HPV16 | 618 | 619 | V-B | BND | N | - | + | 60 | 12 | 30 | Y |
| 3 | chr3 | 184042829 | 184042833 | HPV16 | 5221 | 5233 | B-(V) | BND | N | + | + | 30 | 3 | 11 | Y |
| 4 | chr3 | 184158226 | 184158238 | HPV16 | 6277 | 6289 | (V)-D | BND | N | - | - | 6 | 0 | 4 | Y |
| 5 | chr3 | 184174613 | 184174633 | HPV16 | 2605 | 2608 | D-(V) | BND | N | + | + | 35 | 10 | 15 | Y |
| 6 | HPV16 | 3134 | 3154 | HPV16 | 4350 | 4351 | V-(V) | INV-DUP | N | + | + | 19 | 5 | 12 | N |
| 7 | HPV16 | 3916 | 3917 | HPV16 | 5435 | 5436 | V-V | DEL | N | + | - | 78 | 23 | 42 | N |
| a | chr3 | 184019113 | 184019114 | chr3 | 184019115 | 184019116 | A-B | LANDMARK | Y | + | - | 90 | 28 | 39 | NA |
| b | chr3 | 184042832 | 184042833 | chr3 | 184042834 | 184042835 | B-C | LANDMARK | Y | + | - | 161 | 18 | 45 | NA |
| c | chr3 | 184125373 | 184125374 | chr3 | 184158226 | 184158227 | C-D | LANDMARK | Y | + | - | 107 | 39 | 43 | NA |
| d | chr3 | 184174624 | 184174625 | chr3 | 184174626 | 184174627 | D-E | LANDMARK | Y | + | - | 97 | 30 | 30 | NA |
| v | HPV16 | 0 | 1 | HPV16 | 7905 | 7906 | V-V | LANDMARK | Y | - | + | 7858 | 1178 | 3156 | NA |

**Supplementary Table 3.5. Breakpoints and normal segment junctions in Tumor 3.** Genomic coordinates and other features of virus-virus, virus-host, and host-host breakpoints and of normal segment junctions detected in Tumor 3 upon analysis of WGS and LR-seq data. Description, defined segments connected at junction (see **Supplementary Table 3.6**); normal, presence (Y) or absence (N) of junction in reference genome sequence; type, junction classification by the Sniffles algorithm as duplication (DUP), deletion (DEL), inversion (INV), structural variation breakpoint (BND), or normal reference genome feature (LANDMARK); strand 1 and strand 2, + or - strand as per BEDPE format for WGS data; # reads, number of reads from indicated platform supporting junction; segment-defining, (Y) breakpoint used to define the termini of genomic segments typically >1 kb in length or (N) breakpoint not used to define genomic segments.

| chrom | start | stop | ID | size |
|---|---|---|---|---|
| chr3 | 182969113 | 183969113 | pad1 | 1000000 |
| chr3 | 183969114 | 184019114 | A | 50000 |
| chr3 | 184019115 | 184042833 | B | 23718 |
| chr3 | 184042834 | 184158225 | C | 115391 |
| chr3 | 184158226 | 184174625 | D | 16399 |
| chr3 | 184174626 | 184224626 | E | 50000 |
| chr3 | 184224627 | 185224627 | pad2 | 1000000 |
| HPV16 | 0 | 7906 | V | 7906 |

**Supplementary Table 3.6. Definition of genomic segments flanking HPV integrants in Tumor 3.** Shown here are chromosome coordinates for segments used for the analysis of Tumor 3 LR-seq and WGS data. Chromosome, segment start and stop positions, segment ID, and segment size are indicated. Coordinates are based on GRCh37 human reference genome sequence and HPV16 reference genome (NC_001526.3).

| ID | chrom1 | start1 | stop1 | chrom2 | start2 | stop2 | description | normal | type | strand1 | strand2 | # reads ILMN | # reads PacBio | # reads ONT | segment defining |
|---|---|---|---|---|---|---|---|---|---|---|---|---|---|---|---|
| 0 | chr8 | 43093982 | 43093983 | chr8 | 128809796 | 128809797 | E-(CEN8) | N | INV | - | - | 11* | 6 | 8 | Y |
| 1 | chr8 | 128743987 | 128743988 | chr8 | 128845445 | 128845446 | F-B | N | DUP | - | + | 3912 | 698 | 985 | Y |
| 2 | chr8 | 128746802 | 128746803 | chr8 | 128748660 | 128748661 | B-B | N | DEL | + | - | 102 | 3 | 9 | N |
| 3 | chr8 | 128752228 | 128752229 | chr8 | 128897785 | 128897786 | K-C | N | DUP | - | + | 129 | 25 | 36 | Y |
| 4 | chr8 | 128756497 | 128756515 | chr8 | 128843921 | 128843922 | F-C | N | DUP | - | + | 70 | 15 | 35 | N |
| 5 | chr8 | 128757517 | 128757518 | HPV16 | 4487 | 4488 | C-V | N | BND | + | - | 3404 | 734 | 1072 | Y |
| 6 | chr8 | 128840587 | 128840588 | HPV16 | 3520 | 3521 | E-V | N | BND | + | - | 108 | 15 | 20 | N |
| 7 | chr8 | 128840599 | 128840600 | HPV16 | 1760 | 1761 | V-F | N | BND | - | + | 4424 | 1049 | 1294 | Y |
| 8 | chr8 | 128845271 | 128845274 | HPV16 | 7013 | 7031 | F-V | N | BND | + | - | 35 | 12 | 11 | N |
| 9 | chr8 | 128857554 | 128857574 | chr8 | 128870481 | 128870501 | I-G | N | DUP | - | + | 33 | 9 | 10 | N |
| 10 | chr8 | 128859613 | 128859614 | chr8 | 128866118 | 128866119 | I-H | N | DUP | - | + | 82 | 9 | 19 | Y |
| 11 | chr8 | 128865903 | 128865904 | chr8 | 128871649 | 128871650 | J-I | N | DUP | - | + | 815 | 151 | 250 | Y |
| 12 | chr8 | 128870491 | 128870492 | HPV16 | 5645 | 5646 | I-V | N | BND | + | - | 1446 | 292 | 436 | Y |
| 13 | chr8 | 43097515 | 43097516 | chr21 | 10726025 | 10726026 | CEN8-CEN21 | N | BND | + | - | 4* | 6 | 26 | Y |
| a | chr8 | 128743986 | 128743987 | chr8 | 128743988 | 128743989 | A-B | Y | LANDMARK | + | - | 50 | 7 | 9 | NA |
| b | chr8 | 128752227 | 128752228 | chr8 | 128752229 | 128752230 | B-C | Y | LANDMARK | + | - | 2857 | 689 | 981 | NA |
| c | chr8 | 128757516 | 128757517 | chr8 | 128757518 | 128757519 | C-D | Y | LANDMARK | + | - | 45 | 6 | 9 | NA |
| d | chr8 | 128809797 | 128809798 | chr8 | 128809799 | 128809800 | D-E | Y | LANDMARK | + | - | 46 | 11 | 19 | NA |
| e | chr8 | 128840598 | 128840599 | chr8 | 128840600 | 128840601 | E-F | Y | LANDMARK | + | - | 0 | 0 | 0 | NA |
| f | chr8 | 128845448 | 128845449 | chr8 | 128845447 | 128845448 | F-G | Y | LANDMARK | + | - | 1557 | 260 | 208 | NA |
| g | chr8 | 128859612 | 128859613 | chr8 | 128859614 | 128859615 | G-H | Y | LANDMARK | + | - | 1664 | 247 | 468 | NA |
| h | chr8 | 128865902 | 128865903 | chr8 | 128865904 | 128865905 | H-I | Y | LANDMARK | + | - | 1343 | 298 | 464 | NA |
| i | chr8 | 128870490 | 128870491 | chr8 | 128870492 | 128870493 | I-J | Y | LANDMARK | + | - | 1003 | 90 | 219 | NA |
| j | chr8 | 128871648 | 128871649 | chr8 | 128871650 | 128871651 | J-K | Y | LANDMARK | + | - | 239 | 42 | 63 | NA |
| k | chr8 | 128897784 | 128897785 | chr8 | 128897786 | 128897787 | K-L | Y | LANDMARK | + | - | 77 | 21 | 23 | NA |
| v | HPV16 | 0 | 1 | HPV16 | 7904 | 7905 | V-V | Y | LANDMARK | - | + | 6010 | 1073 | 1333 | NA |

* We did not initially find WGS short reads supporting breakpoint 0, the junction between segment E and Chr. 8 centromere, and breakpoint 13, the junction between Chr. 8 centromere and Chr. 21 centromere, likely due to the repetitive sequences at centromeres. LR-seq enabled us to detect these breakpoints located at hard-to-map regions. Realignment to breakpoint junctions identified in LR-seq reads enabled us to find WGS pairs connecting these segments to CEN-8.

**Supplementary Table 4.1. Breakpoints and normal segment junctions in GUMC-395 cells.** Genomic coordinates and other features of virus-virus, virus-host, and host-host breakpoints and of normal segment junctions detected in GUMC-395 cells upon analysis of WGS and LR-seq data. Description, defined segments connected at junction (see **Supplementary Table 4.2**); normal, presence (Y) or absence (N) of junction in reference genome sequence; type, junction classification by the Sniffles algorithm as duplication (DUP), deletion (DEL), inversion (INV), structural variation breakpoint (BND), or normal reference genome feature (LANDMARK); strand 1 and strand 2, + or - strand as per BEDPE format for WGS data; # reads, number of reads from indicated platform supporting junction; segment-defining, (Y) breakpoint used to define the termini of genomic segments typically >1 kb in length or (N) breakpoint not used to define genomic segments.

| chrom | start | stop | ID | size |
|---|---|---|---|---|
| chr8 | 127693986 | 128693986 | pad1 | 1000000 |
| chr8 | 128693987 | 128743987 | A | 50000 |
| chr8 | 128743988 | 128752228 | B | 8240 |
| chr8 | 128752229 | 128757517 | C | 5288 |
| chr8 | 128757518 | 128809797 | D | 52279 |
| chr8 | 128809798 | 128840599 | E | 30801 |
| chr8 | 128840600 | 128845445 | F | 4845 |
| chr8 | 128845446 | 128859613 | G | 14167 |
| chr8 | 128859614 | 128865903 | H | 6289 |
| chr8 | 128865904 | 128870491 | I | 4587 |
| chr8 | 128870492 | 128871649 | J | 1157 |
| chr8 | 128871650 | 128897785 | K | 26135 |
| chr8 | 128897786 | 128947786 | L | 50000 |
| chr8 | 128947787 | 129947787 | pad2 | 1000000 |
| chr8 | 43093983 | 43097509 | CEN-8 | 3526 |
| chr21 | 10726025 | 10728406 | CEN-21 | 2381 |
| HPV16 | 0 | 7906 | V | 7906 |

**Supplementary Table 4.2. Definition of genomic segments flanking HPV integrants in GUMC-395 cells.** Shown here are chromosome coordinates for segments used for the analysis of GUMC-395 LR-seq and WGS data. Chromosome, start and stop positions, segment ID, and segment size are indicated. Coordinates are based on GRCh37 human reference genome sequence and HPV16 reference genome (NC_001526.3).

| ID | chrom1 | start1 | stop1 | chrom2 | start2 | stop2 | description | normal | type | strand1 | strand2 | # reads ILMN | # reads PacBio | # reads ONT | segment defining |
|---|---|---|---|---|---|---|---|---|---|---|---|---|---|---|---|
| 1 | chr8 | 128230631 | 128230632 | HPV18 | 5735 | 5736 | (V)-B | N | BND | - | - | 342 | 26 | 108 | Y |
| 2 | CP068256.2 (22T) | 51321578 | 51321579 | chr8 | 128232844 | 128232845 | 22T-C | N | BND | + | - | 34* | 1 | 9 | Y |
| 3 | chr8 | 128233364 | 128233367 | HPV18 | 3097 | 3100 | C-(V) | N | BND | + | + | 25 | 1 | 21 | Y |
| 4 | chr8 | 128234254 | 128234255 | HPV18 | 7856 | 7857 | D-(V) | N | BND | + | + | 150 | 14 | 93 | Y |
| 5 | chr8 | 128241547 | 128241548 | HPV18 | 2496 | 2497 | E-(V) | N | BND | + | + | 104 | 14 | 61 | Y |
| a | chr8 | 128230630 | 128230631 | chr8 | 128230632 | 128230633 | A-B | Y | LANDMARK | + | - | 47 | 2 | 34 | NA |
| b | chr8 | 128232843 | 128232844 | chr8 | 128232845 | 128232846 | B-C | Y | LANDMARK | + | - | 389 | 29 | 122 | NA |
| c | chr8 | 128233364 | 128233365 | chr8 | 128233366 | 128233367 | C-D | Y | LANDMARK | + | - | 361 | 29 | 121 | NA |
| d | chr8 | 128234253 | 128234254 | chr8 | 128234255 | 128234256 | D-E | Y | LANDMARK | + | - | 142 | 3 | 74 | NA |
| e | chr8 | 128241546 | 128241547 | chr8 | 128241548 | 128241549 | E-F | Y | LANDMARK | + | - | 72 | 9 | 52 | NA |
| v | HPV18 | 0 | 1 | HPV18 | 7856 | 7857 | V-V | Y | LANDMARK | - | + | 124 | 12 | 83 | NA |

* We did not initially find WGS short reads supporting breakpoint 2, the junction connecting Chr. 22 telomere to segment C. However, we detected this breakpoint, located at a hard-to-map region, by LR-seq. We found 34 WGS short read pairs connecting segment C to simple repeats at 22T after realignment to ONT LR-seq reads.

**Supplementary Table 5.1. Breakpoints and normal segment junctions in HeLa cells.** Genomic coordinates and other features of virus-virus, virus-host, and host-host breakpoints and of normal segment junctions detected in HeLa cells upon analysis of WGS and LR-seq data. Description, defined segments connected at junction (see **Supplementary Table 5.2**); normal, presence (Y) or absence (N) of junction in reference genome sequence; type, junction classification by the Sniffles algorithm as duplication (DUP), deletion (DEL), inversion (INV), structural variation breakpoint (BND), or normal reference genome feature (LANDMARK); strand 1 and strand 2, + or - strand as per BEDPE format for WGS data; # reads, number of reads from indicated platform supporting junction; segment-defining, (Y) breakpoint used to define the termini of genomic segments typically >1 kb in length or (N) breakpoint not used to define genomic segments.

| chrom | start | stop | ID | size |
|---|---|---|---|---|
| chr8 | 127180630 | 128180630 | pad1 | 1000000 |
| chr8 | 128180631 | 128230631 | A | 50000 |
| chr8 | 128230632 | 128232844 | B | 2212 |
| chr8 | 128232845 | 128233364 | C | 519 |
| chr8 | 128233365 | 128234254 | D | 889 |
| chr8 | 128234255 | 128241547 | E | 7292 |
| chr8 | 128241548 | 128291548 | F | 50000 |
| chr8 | 128291549 | 129291549 | pad2 | 1000000 |
| CP068256.2 | 51293895 | 51314834 | 22T | 20939 |
| HPV18 | 0 | 7857 | V | 7857 |

**Supplementary Table 5.2. Definition of genomic segments flanking HPV integrants in HeLa cells.** Shown here are chromosome coordinates for segments used for the analysis of HeLa LR-seq and WGS data. Chromosome, start and stop positions, segment ID, and segment size are indicated. Coordinates are based on GRCh37 human reference genome sequence and HPV18 reference genome (NC_001357.1).

| ID | chrom1 | start1 | stop1 | chrom2 | start2 | stop2 | description | normal | type | strand1 | strand2 | # reads ILMN | # reads PacBio | # reads ONT | segment defining |
|---|---|---|---|---|---|---|---|---|---|---|---|---|---|---|---|
| 1 | HPV16 | 2985 | 2986 | HPV16 | 4084 | 4085 | V-V | N | DUP | - | + | 96 | 32 | 45 | N |
| 2 | HPV16 | 1688 | 1690 | HPV16 | 2773 | 2793 | V-(V) | N | BND | + | + | 77 | 7 | 45 | N |
| 3 | HPV16 | 1942 | 1962 | HPV16 | 4733 | 4752 | V-V | N | DEL | + | - | 53 | 20 | 30 | N |
| 4 | HPV16 | 23 | 29 | HPV16 | 2431 | 2451 | V-(V) | N | BND | - | - | 26 | 1 | 13 | N |
| 5 | HPV16 | 957 | 959 | HPV16 | 7718 | 7720 | V-(V) | N | BND | - | - | 24 | 8 | 39 | N |
| 6 | HPV16 | 5439 | 5446 | HPV16 | 6826 | 6832 | V-V | N | DEL | + | - | 13 | 8 | 15 | N |
| 7 | HPV16 | 4834 | 4854 | HPV16 | 7884 | 7887 | V-(V) | N | DUP | - | + | 39 | 4 | 21 | N |
| 8 | HPV16 | 508 | 509 | HPV16 | 7351 | 7352 | V-(V) | N | BND | - | - | 119 | 35 | 98 | N |
| 9 | chr17 | 36423207 | 36423208 | HPV16 | 1066 | 1067 | V-17B | N | BND | - | + | 207 | 460 | 921 | Y |
| 10 | chr17 | 36424557 | 36424558 | HPV16 | 6457 | 6458 | (V)-17B | N | BND | - | - | 77 | 32 | 58 | N |
| 11 | chr17 | 36425034 | 36425035 | chr17 | 36442270 | 36442271 | 17B-17B | N | DUP | - | + | 16 | 8 | 13 | N |
| 12 | chr17 | 36425102 | 36425122 | HPV16 | 3336 | 3339 | V-(17B) | N | BND | + | + | 44 | 10 | 27 | N |
| 13 | chr17 | 36431389 | 36431390 | chr17 | 36439064 | 36439065 | 17B-(17B) | N | INV | + | + | 127 | 27 | 34 | N |
| 14 | chr17 | 36445082 | 36445083 | chr17 | 36479513 | 36479514 | 17B-17D | N | DEL | + | - | 211 | 468 | 438 | Y |
| 15 | chr17 | 36479587 | 36479588 | HPV16 | 1940 | 1941 | 17D-V | N | BND | + | - | 254 | 469 | 445 | Y |
| 16 | chrX | 33459201 | 33459202 | chrX | 33507238 | 33507239 | XC-XB | N | DUP | - | + | 163 | 21 | 121 | Y |
| 17 | chrX | 33479831 | 33479832 | HPV16 | 3810 | 3811 | XB-V | N | BND | + | - | 196 | 68 | 134 | Y |
| 18 | chrX | 33479834 | 33479835 | HPV16 | 137 | 138 | V-XC | N | BND | - | + | 180 | 110 | 119 | Y |
| 17a | chr17 | 36423205 | 36423206 | chr17 | 36423207 | 36423208 | 17A-17B | Y | LANDMARK | + | - | 16 | 18 | 26 | NA |
| 17b | chr17 | 36445081 | 36445082 | chr17 | 36445083 | 36445084 | 17B-17C | Y | LANDMARK | + | - | 26 | 15 | 27 | NA |
| 17c | chr17 | 36479512 | 36479513 | chr17 | 36479514 | 36479515 | 17C-17D | Y | LANDMARK | + | - | 11 | 16 | 16 | NA |
| 17d | chr17 | 36479586 | 36479587 | chr17 | 36479588 | 36479589 | 17D-17E | Y | LANDMARK | + | - | 9 | 12 | 16 | NA |
| xa | chrX | 33459199 | 33459200 | chrX | 33459201 | 33459202 | XA-XB | Y | LANDMARK | + | - | 62 | 8 | 31 | NA |
| xb | chrX | 33479830 | 33479831 | chrX | 33479832 | 33479833 | XB-XC | Y | LANDMARK | + | - | 13 | 4 | 11 | NA |
| xc | chrX | 33507237 | 33507238 | chrX | 33507239 | 33507240 | XC-XD | Y | LANDMARK | + | - | 43 | 4 | 29 | NA |
| v | HPV16 | 0 | 1 | HPV16 | 7905 | 7906 | V-V | Y | LANDMARK | - | + | 4830 | 1376 | 3184 | NA |

**Supplementary Table 5.3. Breakpoints and normal segment junctions in VU147 cells.** Genomic coordinates and other features of virus-virus, virus-host, and host-host breakpoints and of normal segment junctions detected in VU147 cells upon analysis of WGS and LR-seq data.

Description, defined segments connected at junction (see **Supplementary Table 5.4**); normal, presence (Y) or absence (N) of junction in reference genome sequence; type, junction classification by the Sniffles algorithm as duplication (DUP), deletion (DEL), inversion (INV), structural variation breakpoint (BND), or normal reference genome feature (LANDMARK); strand 1 and strand 2, + or - strand as per BEDPE format for WGS data; # reads, number of reads from indicated platform supporting junction; segment-defining, (Y) breakpoint used to define the termini of genomic segments typically >1 kb in length or (N) breakpoint not used to define genomic segments.

| chrom | start | stop | ID | size |
|---|---|---|---|---|
| chr17 | 35373205 | 36373205 | pad17_1 | 1000000 |
| chr17 | 36373206 | 36423206 | 17A | 50000 |
| chr17 | 36423207 | 36445082 | 17B | 21875 |
| chr17 | 36445083 | 36479513 | 17C | 34430 |
| chr17 | 36479514 | 36479587 | 17D | 73 |
| chr17 | 36479588 | 36529588 | 17E | 50000 |
| chr17 | 36529589 | 37529589 | pad17_2 | 1000000 |
| chrX | 32409199 | 33409199 | padX_1 | 1000000 |
| chrX | 33409200 | 33459200 | XA | 50000 |
| chrX | 33459201 | 33479831 | XB | 20630 |
| chrX | 33479832 | 33507238 | XC | 27406 |
| chrX | 33507239 | 33557239 | XD | 50000 |
| chrX | 33557240 | 34557240 | padX_2 | 1000000 |
| HPV16 | 0 | 7906 | V | 7906 |

**Supplementary Table 5.4. Definition of genomic segments flanking HPV integrants in VU147 cells.** Shown here are chromosome coordinates for segments used for the analysis of VU147 LR-seq and WGS data. Chromosome, segment start and stop positions, segment ID, and segment size are indicated. Coordinates are based on GRCh37 human reference genome sequence and HPV16 reference genome (NC_001526.3).

| ID | chrom1 | start1 | stop1 | chrom2 | start2 | stop2 | description | normal | type | strand1 | strand2 | # reads ILMN | # reads PacBio | # reads ONT | segment defining |
|---|---|---|---|---|---|---|---|---|---|---|---|---|---|---|---|
| 1 | chr8 | 123119884 | 123119885 | HPV16 | 5952 | 5953 | (B)-V | N | BND | - | - | 578 | 102 | 132 | Y |
| 2 | chr8 | 123129149 | 123129150 | chr8 | 128409177 | 128409178 | D-(B) | N | BND | + | + | 535 | 100 | 153 | Y |
| 3 | chr8 | 128397515 | 128397516 | HPV16 | 1990 | 1991 | V-D | N | BND | - | + | 533 | 106 | 125 | Y |
| a | chr8 | 123119884 | 123119885 | chr8 | 123119886 | 123119887 | A-B | Y | LANDMARK | + | - | 89 | 25 | 29 | NA |
| b | chr8 | 123129154 | 123129155 | chr8 | 123129156 | 123129157 | B-C1 | Y | LANDMARK | + | - | 95 | 22 | 30 | NA |
| c | chr8 | 128397515 | 128397516 | chr8 | 128397517 | 128397518 | C2-D | Y | LANDMARK | + | - | 109 | 20 | 35 | NA |
| d | chr8 | 128409178 | 128409179 | chr8 | 128409180 | 128409181 | D-E | Y | LANDMARK | + | - | 129 | 18 | 21 | NA |
| v | HPV16 | 0 | 1 | HPV16 | 7905 | 7906 | V-V | Y | LANDMARK | - | + | 567 | 107 | 129 | NA |

**Supplementary Table 5.5. Breakpoints and normal segment junctions in HTEC cells.** Genomic coordinates and other features of virus-virus, virus-host, and host-host breakpoints and of normal segment junctions detected in HTEC cells upon analysis of WGS and LR-seq data. Description, defined segments connected at junction (see **Supplementary Table 5.6**); normal, presence (Y) or absence (N) of junction in reference genome sequence; type, junction classification by the Sniffles algorithm as duplication (DUP), deletion (DEL), inversion (INV), structural variation breakpoint (BND), or normal reference genome feature (LANDMARK); strand 1 and strand 2, + or - strand as per BEDPE format for WGS data; # reads, number of reads from indicated platform supporting junction; segment-defining, (Y) breakpoint used to define the termini of genomic segments typically >1 kb in length or (N) breakpoint not used to define genomic segments..

| chrom | start | stop | ID | size |
|---|---|---|---|---|
| chr8 | 122069884 | 123069884 | pad1 | 1000000 |
| chr8 | 123069885 | 123119885 | A | 50000 |
| chr8 | 123119886 | 123129155 | B | 9269 |
| chr8 | 123129156 | 123179156 | C1 | 50000 |
| chr8 | 123179157 | 128347515 | pad2 | 5168358 |
| chr8 | 128347516 | 128397516 | C2 | 50000 |
| chr8 | 128397518 | 128409179 | D | 11661 |
| chr8 | 128409180 | 128459180 | E | 50000 |
| chr8 | 128459181 | 129459181 | pad3 | 1000000 |
| HPV16 | 0 | 7906 | V | 7906 |

**Supplementary Table 5.6. Definition of genomic segments flanking HPV integrants in HTEC cells.** Shown here are chromosome coordinates for segments used for the analysis of HTEC LR-seq and WGS data. Chromosome, start and stop positions, segment ID, and segment size are indicated. Coordinates are based on GRCh37 human reference genome sequence and HPV16 reference genome (NC_001526.3).
