## Supplementary Figures for "Intratumoral heterogeneity and clonal evolution induced by HPV integration"

### SUPPLEMENTAL FIGURES

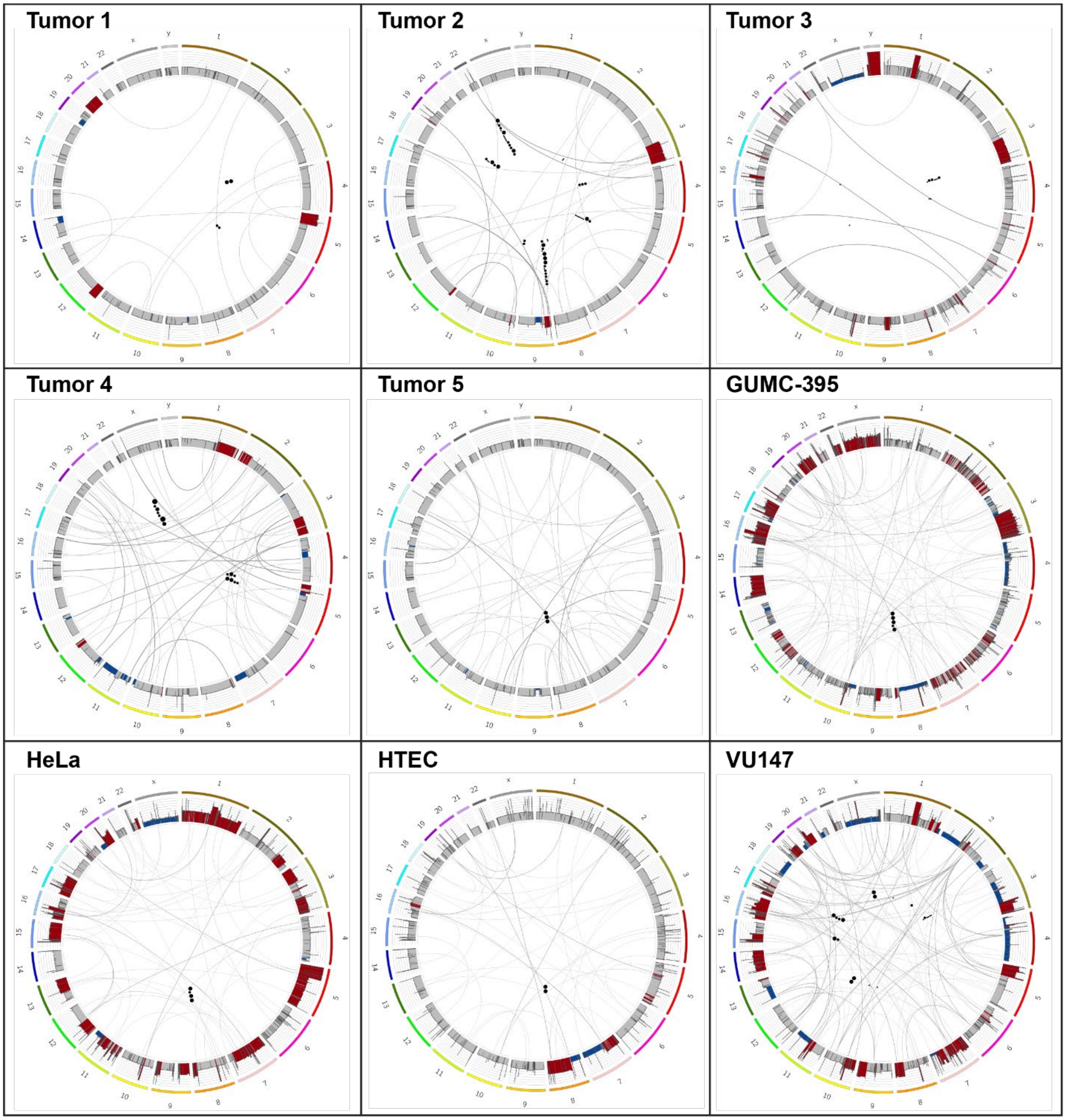

**Supplementary Figure 1.1. Circos plots of human genomes.** Circos plots depict alignments of WGS data (Illumina) to human reference (Chromosomes 1-22, X and Y) generated from HPV-positive primary oropharyngeal cancers and cancer cell lines as indicated. *Barplots*, human genome copy-number alterations: *red*, amplification; *blue*, deletion; *black arcs*, chromosomal translocations; *black dots*, virus-host breakpoints; *dot sizes*, larger with higher number of supporting reads.

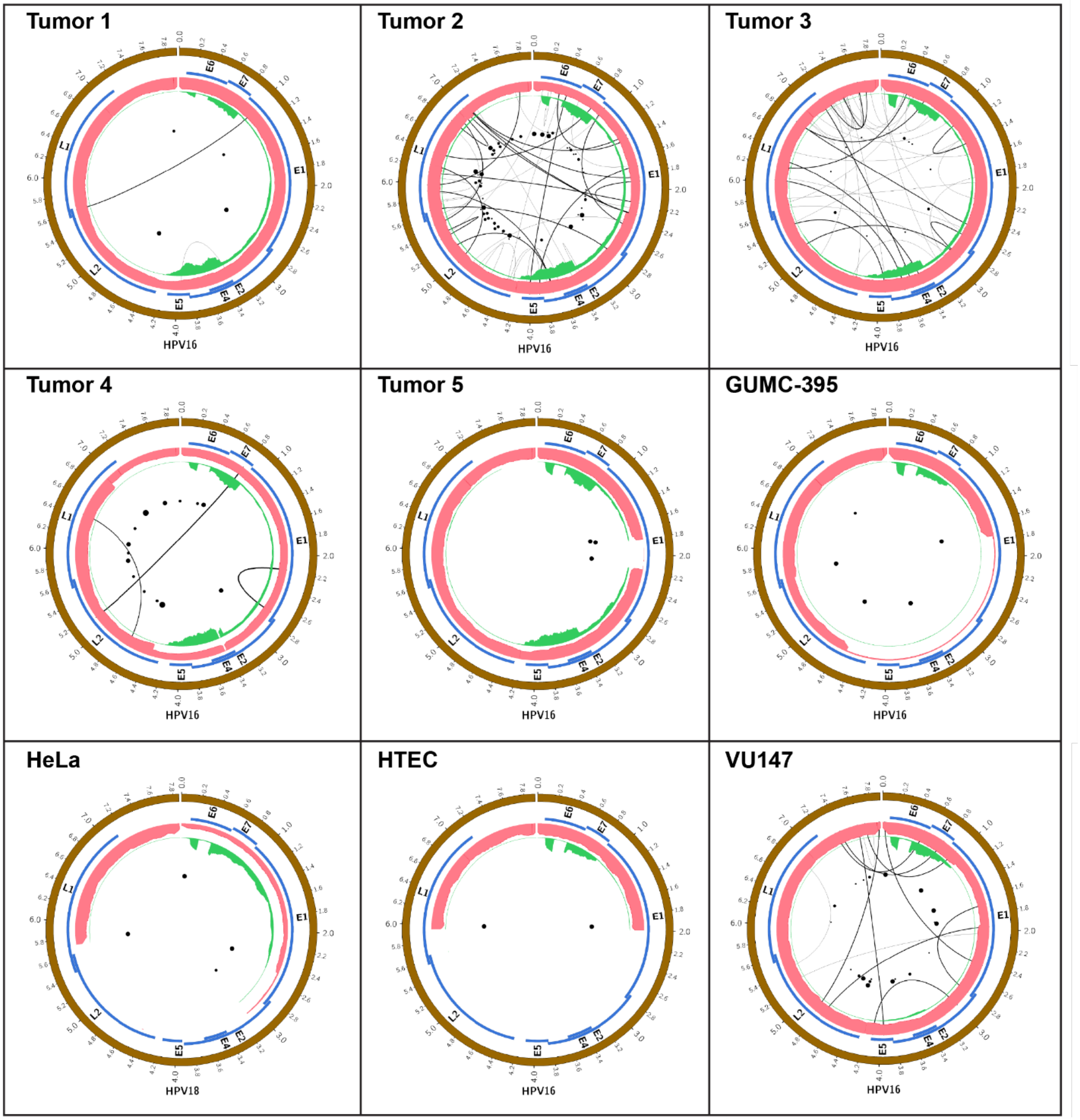

**Supplementary Figure 1.2. Circos plots of HPV genomes.** Circos plots depict alignments of WGS data (Illumina) to HPV genomes generated from HPV-positive primary oropharyngeal cancers and cancer cell lines as indicated. HPV16 or 18 as indicated; *black dots*, virus-host breakpoints; *dot sizes*, larger with higher number of supporting reads; *blue tiles*, virus genes as indicated; *pink histograms*, depth of WGS coverage; *green histograms*, counts of RNA-Seq reads; *arcs*, virus-virus rearrangements.

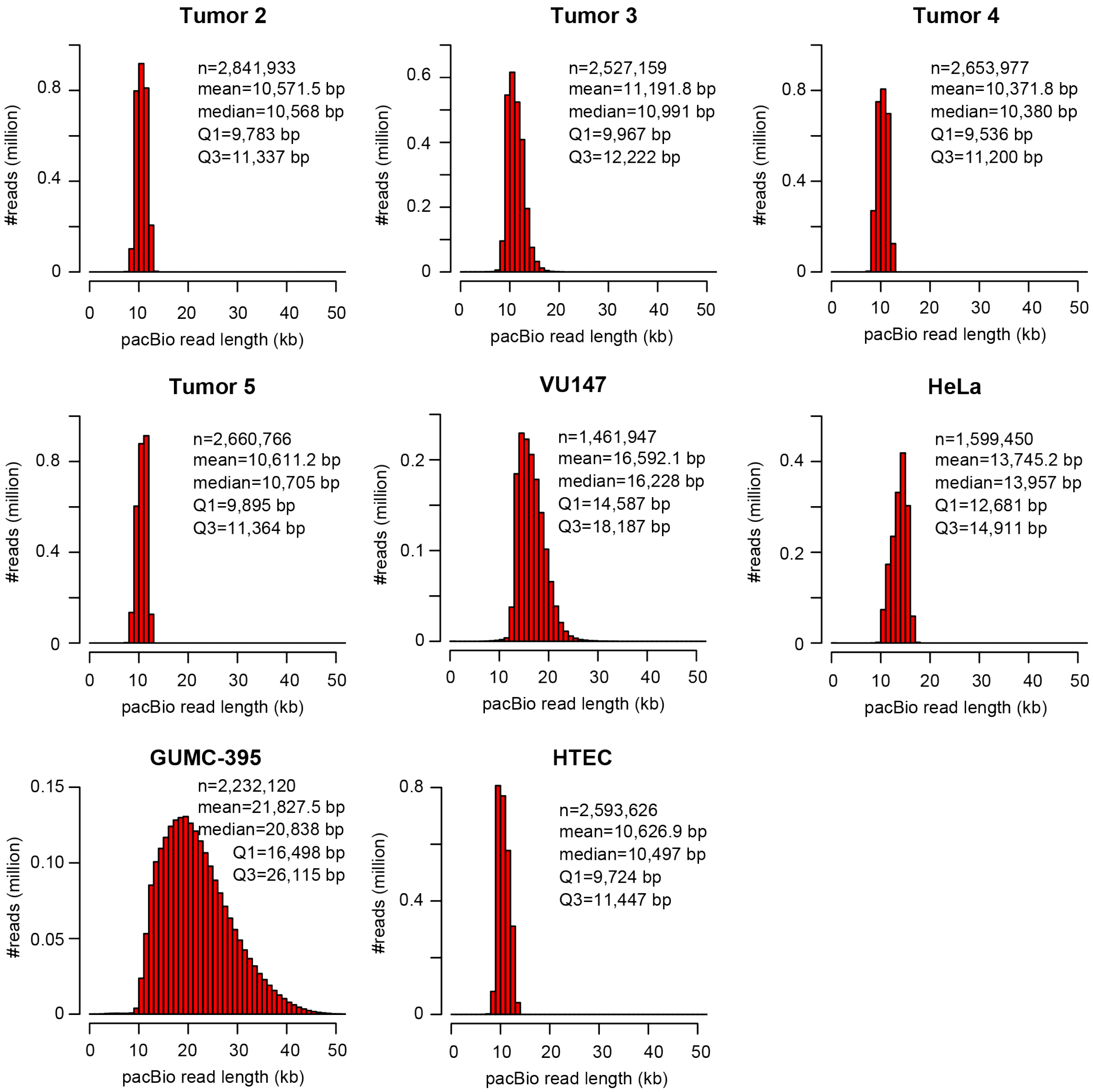

**Supplementary Figure 1.3. PacBio HiFi read length distributions.** Histograms display PacBio HiFi read lengths from primary human oropharyngeal cancers (Tumors 2-5) and cancer cell lines as indicated. Tumor 1 was not sequenced on this platform as genomic DNA was not available. *X-axis*, read length in kilobase pairs (kb); *y-axis*, number of reads per bin size. Shown for each cancer or cell line are total number of reads (n), mean read length, median read length, and the Q1 (25th percentile) and Q3 (75th percentile) for read lengths. Depths of sequencing coverage are reported in **Supplementary Table S1.1**.

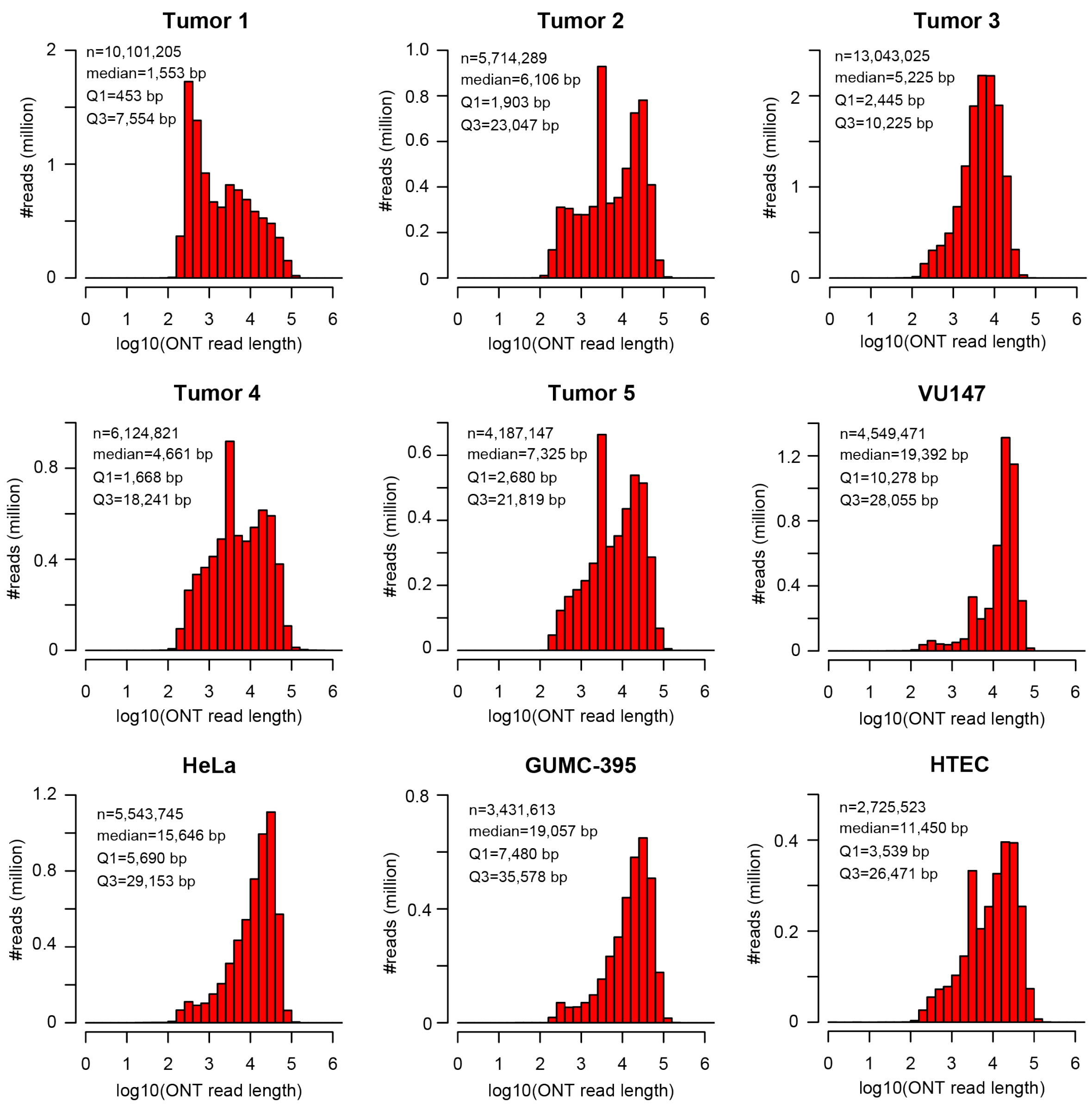

**Supplementary Figure 1.4. Oxford Nanopore (ONT) read length distributions.** Histograms display Oxford Nanopore (ONT) read lengths from primary human oropharyngeal cancers (Tumors 1 - 5) and cancer cell lines as indicated. *X-axis*, read length in kilobase pairs (kb); *y-axis*, number of reads per bin size. Shown within each panel are total number of reads (n), median read length, and the Q1 (25th percentile) and Q3 (75th percentile) for read lengths from each cancer or cell line. Depths of sequencing coverage are reported in **Supplementary Table S1.1**.

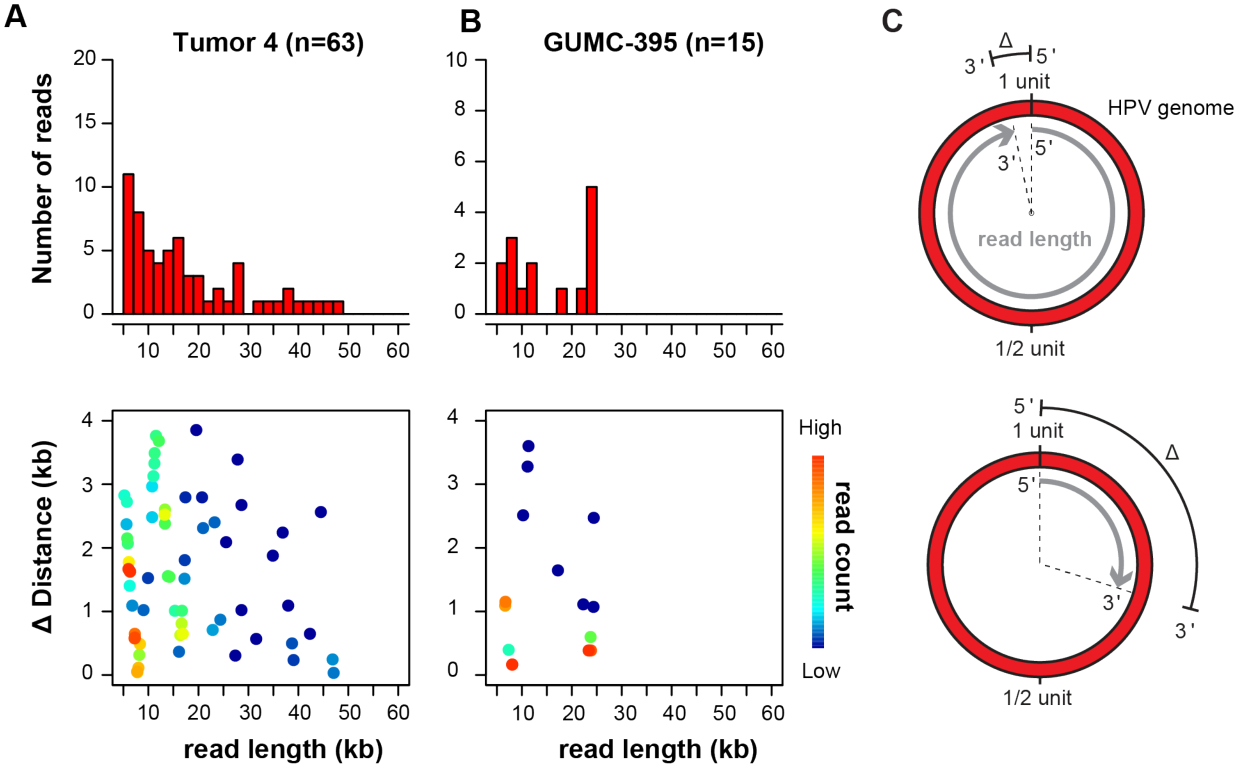

**Supplementary Figure 1.5. Detection of HPV concatemers by LR-seq reads.** ONT LR-seq reads revealed presence of HPV genome concatemers with and without structural variation in (A) Tumor 4 and (B) GUMC-395. Shown are (*top, y-axis*) read count histograms and (*bottom, y-axis*) plots of the distance (Δ) between 5' and 3' mapped coordinates when HPV-only ONT reads were aligned against the HPV16 reference genome. n, number of aligned ONT reads. *X-axis*, *top and bottom*, ONT read lengths in kilobase pairs (kb). *Bottom*, *heatmap*, read counts. (C) Schematic depicting distance (Δ) between read 5' and 3' ends (based on half-maximal genome unit circumference, 7906 ÷ 3953, bp). *Top and bottom*, two example ONT reads (*gray*) aligned against one unit circle of HPV16 genome (*red*). See **Supplementary Table S1.2** for additional details.

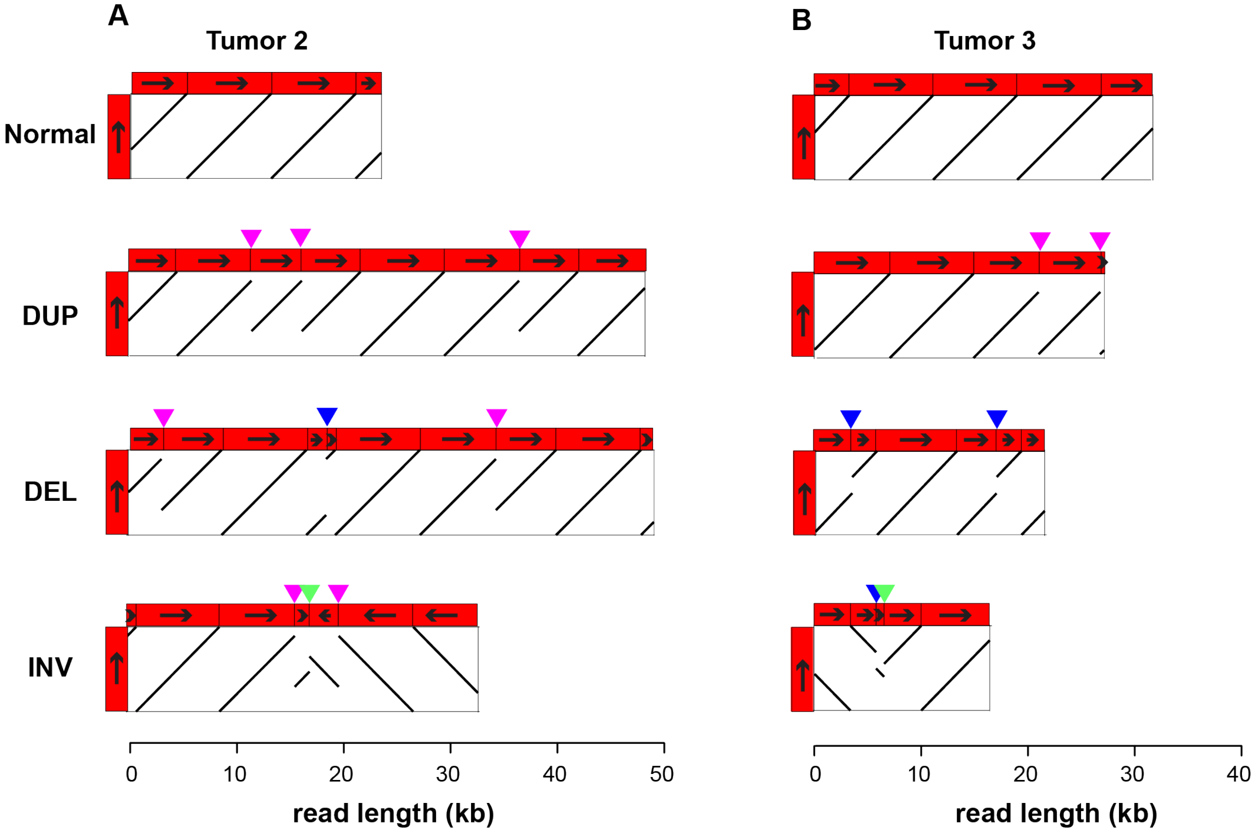

**Supplementary Figure 1.6. ONT reads show structural rearrangements in HPV concatemers.** Dotplots depict (*gray*) alignments of (*x-axis, top*) representative ONT reads of variable lengths against (*y-axis*) one ~7.9 kb unit of HPV genome (*red rectangle*) from (A) Tumor 2 and (B) Tumor 3. Black arrows, orientation of HPV genome from coordinate 1 to 7906. Normal, concatemerized HPV genome without SV; DUP, *pink triangle*, duplication; DEL, *blue triangle*, deletion; INV, *green triangle*, inversion. *X-axis, bottom*, ONT read length.

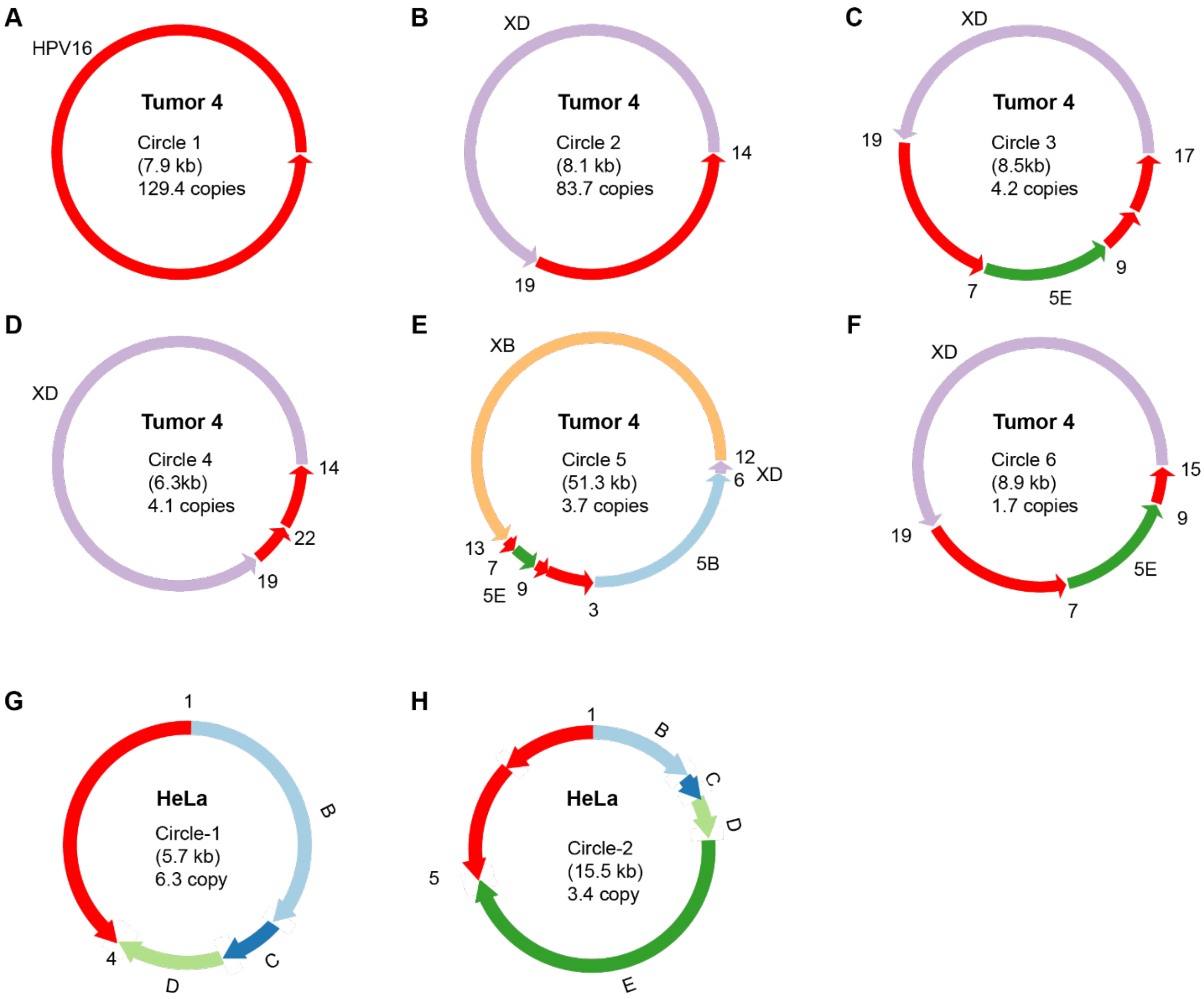

**Supplementary Figure 2. Inaccurate AmpliconArchitect predictions of HPV ecDNA structures in Tumor 4 and HeLa.** WGS (Illumina) short-read data from (A-F) Tumor 4 and (G-H) HeLa were evaluated for presence of ecDNA using AmpliconArchitect. AmpliconArchitect predicted 6 different HPV-containing ecDNA structures in Tumor 4: (A) HPV16 virus-only ecDNA; (B) host XD segment connected to HPV16 at breakpoints 19 and 14; (C) circles comprising XD-HPV-5E-HPV segments, connected at breakpoints 19, 7, 9 and 17; (D) XD segments connected to HPV16 at breakpoints 19, 22, and 14; (E) circles comprising XB-HPV-5E-HPV-5B-XD segments connected at breakpoints 13, 7, 9, 3, 6, and 12; and (F) circles comprising XD-HPV-5E-HPV segments connected at breakpoints 19, 7, 9, and 15. AmpliconArchitect predictions for Tumor 4 were oversimplified and inaccurate when compared structures supported by LR-seq data (see **Fig. 2**, **Supplementary Tables S2.1, S2.2**). AmpliconArchitect predicted 2 different ecDNA structures in HeLa: (G) Segment B-C-D connected to the HPV18 genome by breakpoints 1 and 4; and (H) Segment B-C-D-E connected to the HPV18 genome by breakpoints 1 and 5. In contrast, ONT data supported more complex structures containing alternating concatemers of segments B-C-D-(V) and B-C-D-E-(V) that were anchored into flanking sequences in Chr. 8 (see **Fig.6B**).
